# The landscape of regional missense mutational intolerance quantified from 730,947 exomes

**DOI:** 10.1101/2024.04.11.588920

**Authors:** Lily Wang, Katherine R. Chao, Ruchit Panchal, Calwing Liao, Haneen Abderrazzaq, Robert Ye, Patrick Schultz, John Compitello, Riley H. Grant, Jack A. Kosmicki, Ben Weisburd, William Phu, Michael W. Wilson, Kristen M. Laricchia, Julia K. Goodrich, Daniel Goldstein, Jacqueline I. Goldstein, Christopher Vittal, Timothy Poterba, Samantha Baxter, Nicholas A. Watts, Matthew Solomonson, Genome Aggregation Database Consortium, Grace Tiao, Heidi L. Rehm, Benjamin M. Neale, Michael E. Talkowski, Daniel G. MacArthur, Anne O’Donnell-Luria, Konrad J. Karczewski, Predrag Radivojac, Mark J. Daly, Kaitlin E. Samocha

## Abstract

Missense variants can have a range of functional impacts depending on factors such as the specific amino acid substitution and location within the gene. To interpret their deleteriousness, studies have sought to identify regions within genes that are specifically intolerant of missense variation. Here, we leverage the patterns of rare missense variation in 730,947 exome sequenced individuals in the Genome Aggregation Database (gnomAD v4.1.1) against a null mutational model to identify transcripts with regional differences in missense constraint. Missense-depleted regions are enriched for ClinVar pathogenic variants, *de novo* missense variants from individuals with neurodevelopmental disorders, and complex trait heritability. Following ClinGen calibration recommendations for the ACMG/AMP variant classification guidelines, we establish that variants within regions with <36% of their expected missense variation achieve moderate support for pathogenicity. We integrate this regional constraint measure into a missense deleteriousness metric (named MPC) that effectively stratifies rare and *de novo* missense variants in individuals with early-onset developmental conditions from controls. These results provide additional tools to aid in missense variant interpretation.

## Main text

Over the last decade, exome and genome sequencing have enabled variant discovery across hundreds of thousands of individuals^1–6^. As sequencing becomes more routine in both clinical and research settings, it is crucial to improve our ability to interpret the potential impacts of these variants, particularly in understanding disease risk. With the rapidly expanding catalog of observed genomic variation comes an increasingly large number of variants of uncertain significance (VUS) that cannot be conclusively considered harmful (pathogenic) or neutral (benign)^7^. Indeed, well over half of the variants in the ClinVar database^8^, which hosts disease-variant associations, are considered VUS, with thousands more added every year^9^.

Classification and interpretation of rare variants has been greatly aided by the release of large reference databases, which have provided the opportunity to study selective forces acting on the human genome and to identify genomic regions under selective constraint by, for example, identifying regions with fewer variants than expected based on mutational models^2,3,10–13^. In particular, gene-level metrics of predicted loss-of-function (pLoF) variant depletion have proven to be valuable in variant classification and identification of novel gene-disease relationships^14–18^. The functional impact and selective pressures relevant to missense variation, by contrast, remain challenging to predict, as the effect of a missense variant is governed by the gene housing the variant, the position of the variant in the gene, and the specific amino acid substitution caused by the variant. In fact, 90% of the missense variants in ClinVar are VUS^9,19^. To address this, prior work has sought to identify regions within coding genes that are specifically intolerant of missense variation as a way to improve classification^20–31^.

One major application of constraint metrics is in novel gene identification, particularly when searching for a significant burden of *de novo* (newly arising) variants. Both missense deleteriousness metrics and region-specific missense depletion information have been used to improve power to identify genes associated with neurodevelopmental disorders^15,16,32^, as both approaches help separate signal from noise by highlighting a subset of missense variants that are more likely to contribute to rare disorders. Beyond enrichment in statistical frameworks, these metrics can also be incorporated into clinical workflows for variant classification. This requires that the metrics are calibrated for use in the published guidelines for evaluating variant pathogenicity from the American College of Medical Genetics and Genomics and the Association for Molecular Pathology (ACMG/AMP)^33^. These ACMG/AMP guidelines provide recommendations for how various lines of evidence (e.g., computational, functional) can be applied towards determining if a variant is pathogenic or benign as well as the strength of the evidence (e.g., strong, moderate). For a given type of evidence, such as *in silico* computational tools^34,35^ or functional data from multiplex assays of variant effects^36–38^, probabilistic frameworks can be used to determine the strength of evidence.

Here, we expand upon previous work^20^ and develop a sub-genic measure of missense intolerance leveraging population-level variation. We show that this measure facilitates variant classification and risk stratification for association studies with *de novo*, rare, and common variants. Specifically, we identify that 36% of transcripts have evidence of variable missense constraint, allowing us to refine the missense constraint scores for 50% of all coding bases to the sub-genic region-level rather than gene-level scores. In missense-depleted coding regions, we find strong enrichments of disease-associated variants from ClinVar and *de novo* missense variants identified in individuals with neurodevelopmental disorders (NDDs). For the first time, we apply a clinical calibration framework to assign a strength of evidence for regional missense depletion that can be used in clinical classification workflows, demonstrating moderate support for pathogenicity. Finally, we combine our regional missense constraint results with variant- and amino acid-level evidence to create the MPC (Missense deleteriousness Prediction by Constraint) score that effectively stratifies rare variants identified in individuals with NDDs from those found in controls^39–41^.

## Results

### Identification of missense constraint regions within transcripts

We explored the patterns of rare missense variant presence or absence in 730,947 exomes in the Genome Aggregation Database (gnomAD) v4.1.1 to quantify missense depletion at the sub-genic level. We searched 17,841 MANE Select or canonical protein-coding transcripts that passed quality control (see **Methods, Supplementary Table 1**) for variability in missense constraint, quantified as the number of rare (allele frequency [AF] < 0.1%) missense variants observed in gnomAD divided by the number expected under neutral evolution as estimated from previously described mutational models^3^. For each transcript, we applied a recursive search based on likelihood ratio tests over all potential rare missense sites looking for breaks that divide the transcript coding sequence into sequences with statistically significantly different missense observed/expected (OE) values (see **Supplementary Note** and **Supplementary Fig. 1**), which we refer to as missense constraint regions (MCRs; **Fig. 1a-b**). Through this approach, we discover 6,361 transcripts (36%) that harbor regional variability in missense constraint (**Fig. 1c**), i.e., have two or more MCRs (minimum coding length 20bp, median 339bp; **Supplementary Figs. 2-3**). We thus refine the resolution of missense constraint for 50% of coding sites (15.7 coding Mb in the 6,361 transcripts vs. 31.2 coding Mb in all 17,841 transcripts assessed), discovering widespread signatures of negative and neutral selection obscured when measuring constraint at transcript-resolution (**Fig. 1d**). For example, relative to the transcript-resolution missense constraint distribution, a larger proportion of the coding exome lies within strongly constrained sequences (e.g., 3.3% vs. 0.7% at OE < 0.4 for all genes; see **Supplementary Note** for OE threshold selection) and the mode shifts toward an OE indicative of evolutionary neutrality at approximately 1 (49.8% vs. 46.4% at 0.9 < OE ≤ 1.1). Across coding sequences, the median missense OE is 0.92 and the top 10% most constrained sites have a missense OE ≤ 0.63 (**Supplementary Table 2**).

**Fig. 1:**
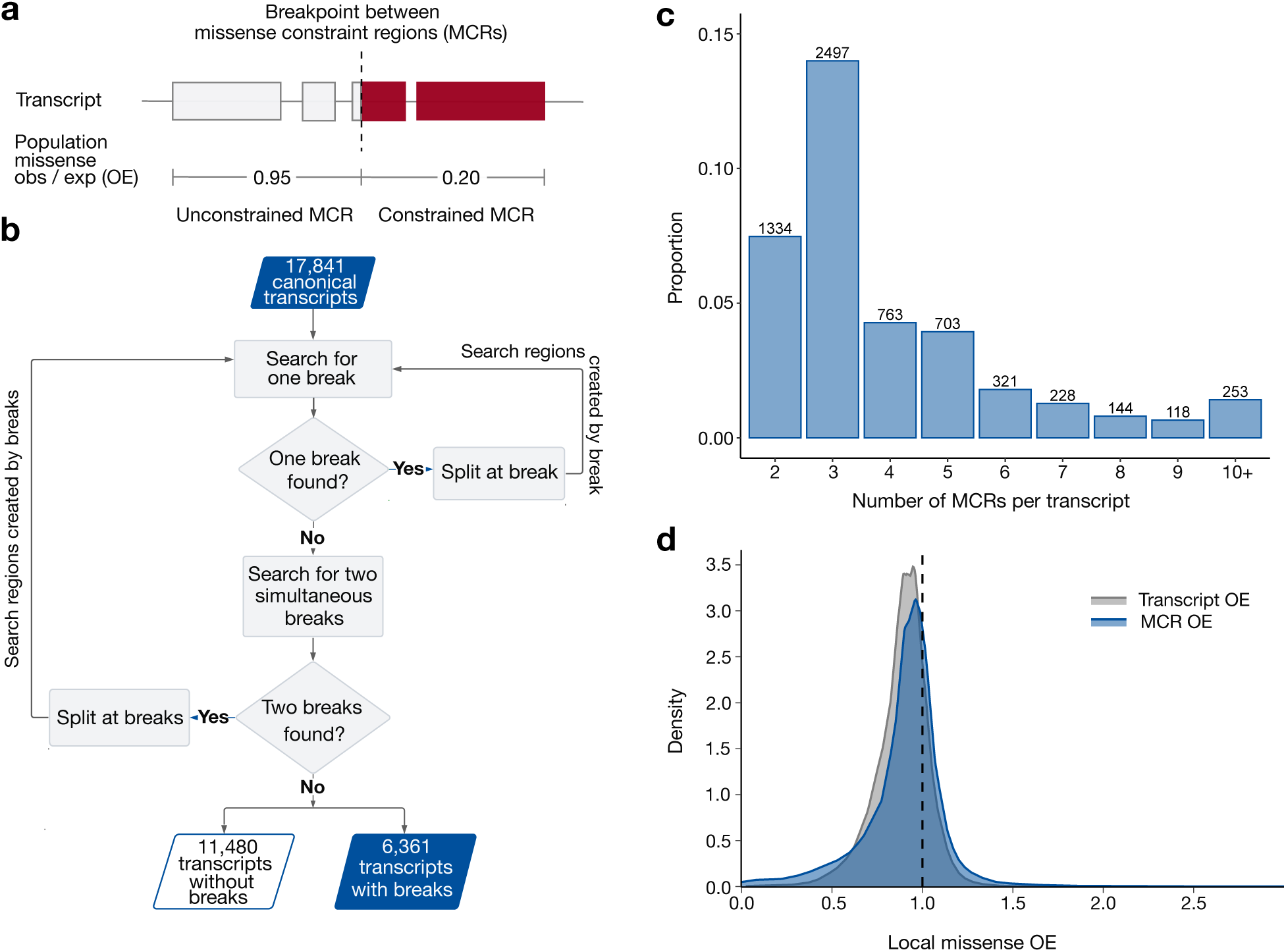
36% of protein-coding genes in the human genome harbor regional variation in population-level missense depletion. **a**, An example transcript that has two missense constraint regions (MCRs) with significantly different levels of population-wide missense depletion, defined as the number of missense variants observed in gnomAD at rare frequency (AF < 0.1%) divided by the number of rare missense variants expected under neutral evolution (observed/expected or OE). Lower OE values correspond to greater variant depletion and suggest stronger selective constraint. **b**, Flow chart describing the process of searching for breakpoints that divide a transcript into multiple MCRs. Searching for breakpoints is recursive and leverages likelihood ratio tests at a significance threshold of p = 0.001. **c**, The number of MCRs identified within the 6,361 transcripts where we discover regional differences in missense constraint. The other 11,480 transcripts are deemed to have a single MCR (i.e., a constant level of constraint across their entirety) and are not shown. **d**, The distribution of local missense OE at all coding sites in MANE Select or canonical transcripts. Local missense OE is defined as the OE calculated over the whole transcript (for “transcript OE”) or over the MCR (for “MCR OE”) where the site is located. Transcript OE and MCR OE are equivalent for transcripts with one MCR. The areas under each of the two curves sum to 1.

We find that missense constraint is able to identify regions associated with severe, early-onset disease. One example is in *BAP1*, which plays a key role in chromatin modeling by mediating histone deubiquitination^42–44^. BAP1 has a conserved N-terminal catalytic domain (UCH) and a predicted C-terminal UCHL5/UCH37-like domain (ULD) likely involved in complexing with other proteins and nuclear localization^45–47^. Disease-causing variants in this gene are linked to both cancer^42,48,49^ and a recently discovered rare NDD named Küry-Isidor syndrome^44^. We identify two highly constrained MCRs in this gene with missense OEs of 0.50 (6th percentile) and 0.57 (8th percentile) at the N- and C-termini, closely overlapping the annotated gene domains.

Despite covering only 43% of the gene (944/2,187 bp), these two constrained MCRs capture all 13 missense variants reported to be causal for Küry-Isidor in ClinVar or the initial publication^44^ (binomial p of enrichment in the constrained MCRs = 1.8×10^−5^, **Fig. 2a**). The only ClinVar pathogenic/likely pathogenic (P/LP) variants that do not fall within any missense-constrained MCRs in *BAP1* (n=2) are associated with cancer predisposition diseases and are likely under weaker selection than early-onset NDDs. Overall, 26/28 (93%) of all ClinVar P/LP variants in *BAP1* reside in constrained MCRs (binomial p = 4.2×10^−8^).

**Fig. 2:**
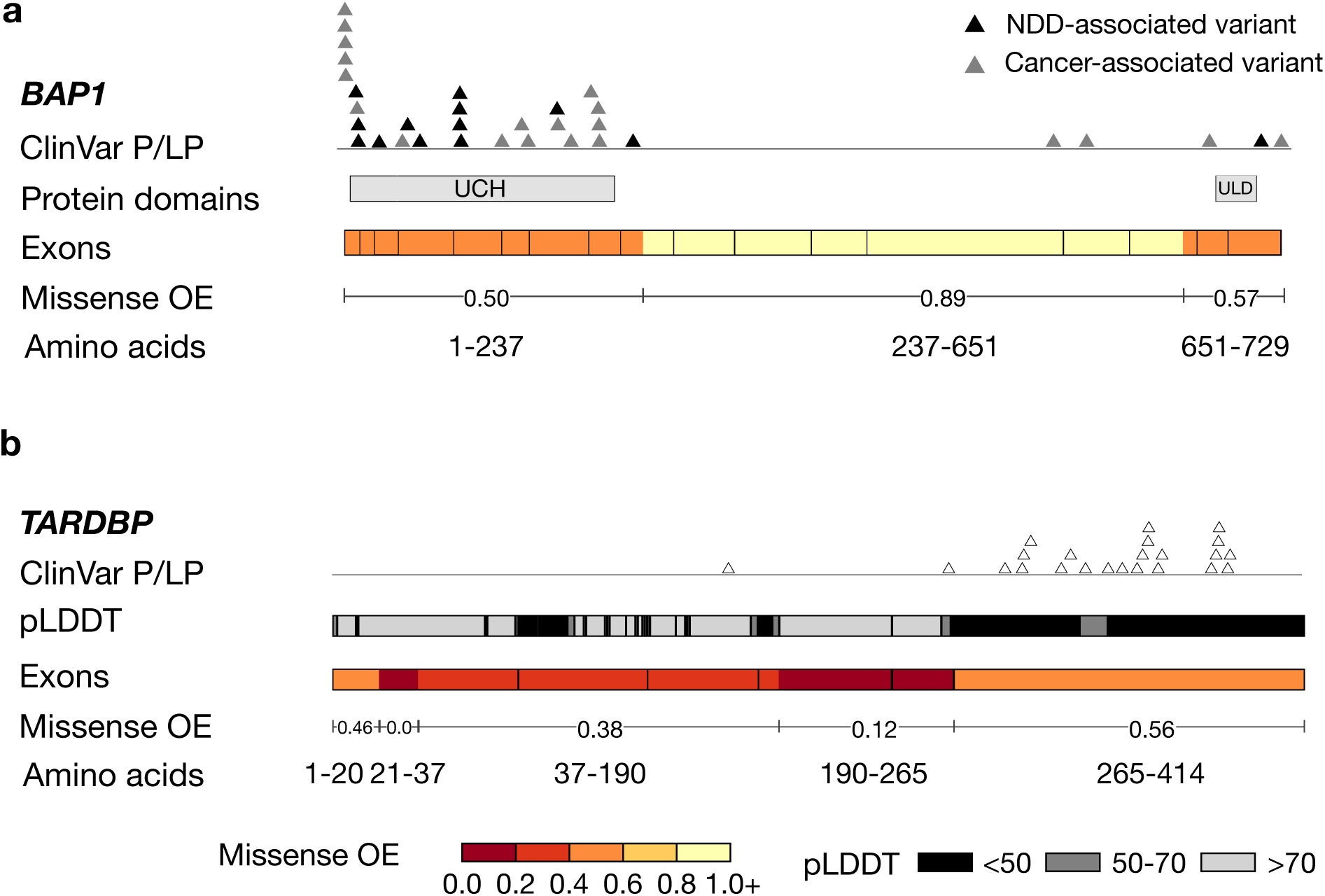
Missense constraint regions (MCRs) and the distribution of ClinVar pathogenic/likely pathogenic (P/LP) missense variants in two disease-associated genes. Exons are delineated with black outlines and MCRs are delineated by color based on their missense observed/expected (OE) ratio. **a,** *BAP1*. Variants in this gene can lead to cancer-predisposition syndromes, increased risk of certain cancers, or the neurodevelopmental disorder Küry-Isidor syndrome^44^. All of the variants associated with Küry-Isidor in ClinVar or the initial publication fall within the highly constrained N-terminal or C-terminal MCRs (6th and 8th percentiles, respectively). Küry-Isidor variants are colored in black and all other cancer-associated P/LP missense variants are colored in gray. UCH: Ubiquitin C-terminal hydrolase domain, ULD: UCHL5/UCH37-like domain. **b,** *TARDBP*. Missense variants in this gene can cause adult-onset neurodegenerative disorders. The C-terminal constrained MCR (8th percentile), where the majority of the ClinVar P/LP variants are located, aligns to a predicted-disordered region of the protein. pLDDT: predicted Local Distance Difference Test from AlphaFold.

We also find that missense constraint can identify disease-associated gene regions beyond annotated protein domains. For example, *TARDBP*, which encodes a regulator of RNA processing and metabolism, is an extremely pLoF-constrained gene^50^ and is classically associated with amyotrophic lateral sclerosis (ALS) and accompanying frontotemporal dementia (FTD)^51–54^. All 23 ClinVar P/LP variants in this gene cause a missense change, and all but one reside in the protein C-terminus, which is predicted to be disordered by AlphaFold^55^ and MobiDB-lite in UniProt^45,56^ (**Fig. 2b**). Our missense constraint metric assigns strong constraint over the entirety of this gene and identifies a highly constrained C-terminal MCR (OE = 0.56, 8th percentile; amino acids [AAs] 265-414) that aligns precisely with the predicted-disordered C-terminal region of the gene (AAs 259-414 with AlphaFold predicted Local Distance Difference Test [pLDDT] < 50) and includes 21 of the 22 ClinVar variants at this end of the gene. Notably, this particular MCR has the weakest constraint of all five MCRs identified in *TARDBP*, but would be missed by most 3D structural approaches that remove regions with low pLDDT^30^. Taken together with the extreme pLoF constraint in this gene yet lack of disease-associated pLoF variants, this suggests two things. First, it is likely the currently known neurodegenerative conditions are some of the least severe diseases caused by disruptions of *TARDBP,* and LoF or other damaging variants outside of the disordered C-terminus are likely to cause unrecognized severe rare disorders or be incompatible with life. Second, missense variants and constraint can provide a window into understanding gene functions and disease mechanisms when pLoF-based approaches are more opaque.

### Enrichment of disease-associated variation in missense-depleted regions

Next, we sought to determine if the signatures of selection revealed by MCRs more broadly recapitulated biological and disease relevance of coding sequences. Here, we filtered to high-quality, high-coverage sequences (95% of MCRs or 98.3% of coding space; see **Supplementary Note** and **Supplementary Fig. 4** for discussion of lower-coverage sequences). Overall, transcripts that are more intolerant to pLoF variation, as measured by the loss-of-function observed/expected upper bound fraction (LOEUF) score^3,50^, also tend to be more intolerant to missense variation. This trend is markedly more prominent when measuring intolerance via the most constrained MCR in a transcript vs. across the whole transcript (Spearman ρ = 0.58 vs. 0.51; **Supplementary Note** and **Supplementary Fig. 5**). We confirmed that, as expected, autosomal genes without disease associations in OMIM^57^ tended to be less missense-constrained than genes with autosomal dominant disease inheritance (Welch’s t p < 10^−50^ for transcript missense OE and Wilcoxon p < 10^−50^ for minimum MCR missense OE).

However, we found that these non-OMIM genes were also more missense-constrained than genes with autosomal recessive disease inheritance (Welch’s t p = 1.1×10^−16^ for transcript missense OE and Wilcoxon p = 9.0×10^−7^ for minimum MCR missense OE; **Supplementary Fig. 6**). Taken together with our finding that 62% (1,511/2,430) of genes that are both LOEUF-constrained and severely missense-depleted (first three deciles of LOEUF and MCR with OE < 0.4) do not have disease associations in OMIM, this highlights a potential search space for novel gene-disease relationships (**Supplementary Fig. 7**).

We defined a set of 695 strongly pLoF-intolerant genes as those in the first three deciles of LOEUF or either of two s_het_ measures^58,59^ and with monoallelic association with a developmental phenotype in Gene2Phenotype (G2P)^60^. In these genes, we observed strong transcript-wide missense depletion that was even stronger for genes with multiple MCRs (**Fig. 3a**; Wilcoxon p < 10^−50^). As a comparator, we defined 1,580 tolerant genes as those in the last three deciles of LOEUF and the two s_het_ measures and not associated with a phenotype in OMIM. Intolerant genes were much more likely to harbor multiple MCRs than tolerant genes, at a 62-fold increased rate after regressing out transcript coding length (612/695 intolerant genes vs.

**Fig. 3:**
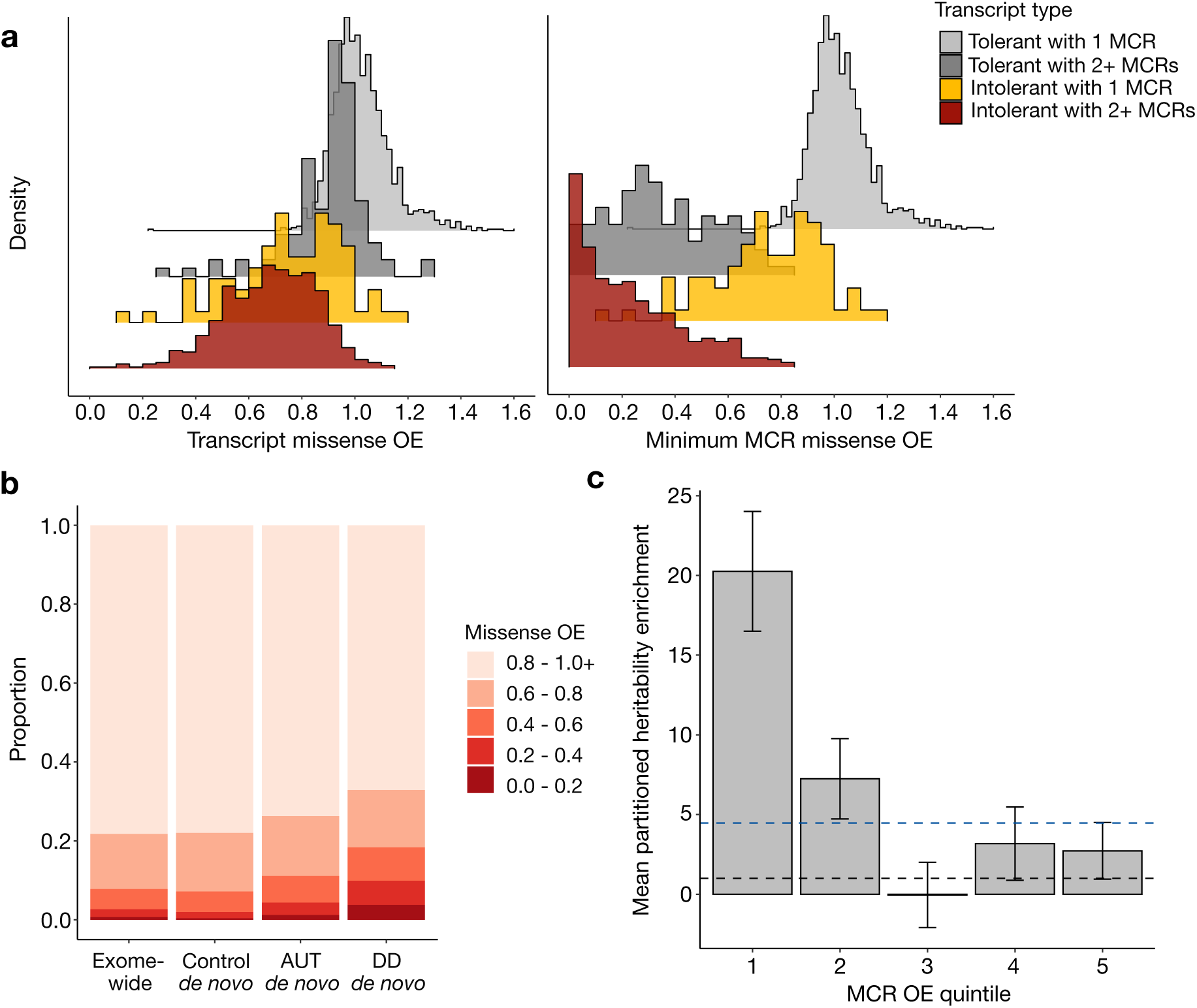
Regional missense depletion reveals constraint obscured by gene-level measures. **a**, Left: The distribution of transcript-wide missense observed/expected (OE) across 17,841 transcripts stratified by the combination of two factors: whether the transcript is strongly mutationally intolerant or tolerant and whether we detect multiple missense constraint regions (MCRs). Number of transcripts in each category are: strongly intolerant with multiple MCRs (n=612; red), strongly intolerant with one MCR (n=83; yellow), tolerant with multiple MCRs (n=103; dark gray), tolerant with one MCR (n=1,477; light gray). Right: Minimum MCR missense OE using the same groupings. For transcripts with a single MCR, minimum MCR missense OE is equivalent to transcript missense OE. **b**, MCR missense OE at all sites of possible exome-wide missense variants vs. sites of *de novo* missense variants in controls, autistic individuals (AUT), or individuals with developmental disorders (DDs). *De novo* variants from individuals with developmental phenotypes are enriched in more constrained sequences, with a more pronounced enrichment in DD than autism. **c**, Enrichment in per-variant heritability explained by common (AF > 5%) protein-coding SNPs stratified by MCR missense OE quintile, relative to the average SNP genome-wide. Enrichment is estimated by linkage disequilibrium score regression, accounting for number of SNPs in each quintile, and is averaged across 268 independent traits in the UK Biobank and other large genome-wide association studies. Black dashed line at 1 indicates no enrichment. Blue dashed line at 4.5 indicates average coding enrichment. Error bars represent 95% confidence intervals.

103/1,580 tolerant genes with multiple MCRs, logistic regression coefficient p < 10^−50^). Intolerant genes were also highly enriched for severely depleted regions (2.3-fold more likely to have minimum MCR OE < 0.4 after regressing out transcript length, logistic regression coefficient p = 4.3×10^−4^), whereas the most constrained MCRs in tolerant genes were overall less depleted and more evenly distributed across the OE spectrum. We next compared missense constraint to selection over longer timescales, measured by the phyloP^61^ score for conservation across placental mammals. At the transcript-level, we found that genes with more conserved coding sequences also tended to be more depleted of human missense variation overall (Spearman ρ = 0.50, p < 10^−50^). However, at the MCR-level, we discovered a substantial number of strongly constrained MCRs that appear widely unconserved across mammals, suggesting human-specific negative selection pressures that are obscured at the whole-transcript level (**Supplementary Fig. 8**, **Supplementary Tables 3-4**).

We next aggregated *de novo* missense variants from 31,058 individuals with a severe developmental disorder^15^ (DD), 15,036 autistic individuals (AUT), and their 5,492 siblings not diagnosed with a DD^16^ (**Fig. 3b**, **Supplementary Fig. 9, Supplementary Tables 5-8**). The distribution of *de novo* missense variants across the missense OE spectrum in unaffected siblings largely mirrored the exome-wide missense OE distribution. In contrast, *de novo* missense variants in autistic individuals are enriched in missense-constrained sequences, with an even stronger effect observed in individuals with DDs. For example, relative to unaffected siblings, the rate of *de novo* missense variants in MCRs with OE < 0.6 is 3-fold higher in individuals with DDs (Poisson two-sided p < 10^−50^) and 1.7-fold higher in autistic individuals (Poisson two-sided p < 10^−50^; **Supplementary Fig. 9**; see **Supplementary Note** for OE threshold selection). This is consistent with the expectations that a small subset of *de novo* missense variants in individuals with developmental phenotypes are causal for those traits and that variants causal for DD are generally more selectively deleterious than those for autism.

Beyond large-effect rare and *de novo* variation in traits under strong negative selection, we additionally investigated whether our MCR metric, which was calculated using rare variants, correlated with functional effects of common variants. Prior work found that pLoF-constrained genes and their flanking 100kb sequences are enriched for SNP heritability across hundreds of independent traits in the UK Biobank and other large genome-wide association studies (GWAS)^3^. We partitioned common (AF > 5%) variant heritability of the same 268 independent traits across MCRs to investigate relative enrichment. To establish a baseline, we computed the heritability enrichment relative to the average genome-wide SNP over all coding sequences comprising MCRs (4.5-fold). The most constrained MCRs have the strongest heritability enrichment: e.g., the first quintile of MCR missense OE harbors a 20-fold enrichment (**Fig. 3c**, **Supplementary Fig. 10, Supplementary Tables 9-10**). Coding SNPs in missense-unconstrained MCRs (e.g., in the three least constrained quintiles of MCR missense OE) harbor little to no detectable heritability enrichment. These findings suggest that regions depleted of rare missense variation can prioritize common coding variants important for complex traits (e.g., improve GWAS fine-mapping variant prioritization), and that there exists a subset of coding sequences with no appreciable heritability enrichment relative to the average genome-wide SNP, which constraint can help identify.

### Evaluating the clinical utility of missense depletion

We then assessed the ability of MCRs to recapitulate clinical disease information by examining the constraint distribution around rare ClinVar P/LP and benign/likely benign (B/LB) missense variants. Across genes with autosomal dominant disease inheritance and ≥2 MCRs, we found that P/LP missense variants tend to occur in regions that are more strongly constrained than the transcript average (n=529 genes, n=29,928 variants, paired Welch’s t p < 10^−50^), while the converse was true for B/LB variants (n=615 genes, n=51,705 variants, paired Welch’s t p < 10^−50^; **Fig. 4a**, **Supplementary Fig. 11a-b, Supplementary Table 11**). For example, in 337 genes with both constrained (missense OE < 0.6) and unconstrained (missense OE > 0.9) MCRs, P/LP variants occur 11-fold more frequently in constrained than unconstrained MCRs, while B/LB variants occur 9-fold more frequently in unconstrained than constrained MCRs (odds ratio [OR] = 93, Fisher’s exact p < 10^−50^). These patterns are also significant, though more mild, in genes associated with autosomal recessive inheritance (**Supplementary Fig. 11c-d, Supplementary Table 12**). Together, these results demonstrate that our constraint metric is able to distinguish specific sequences within genes that are enriched for pathogenic or benign variants.

**Fig. 4:**
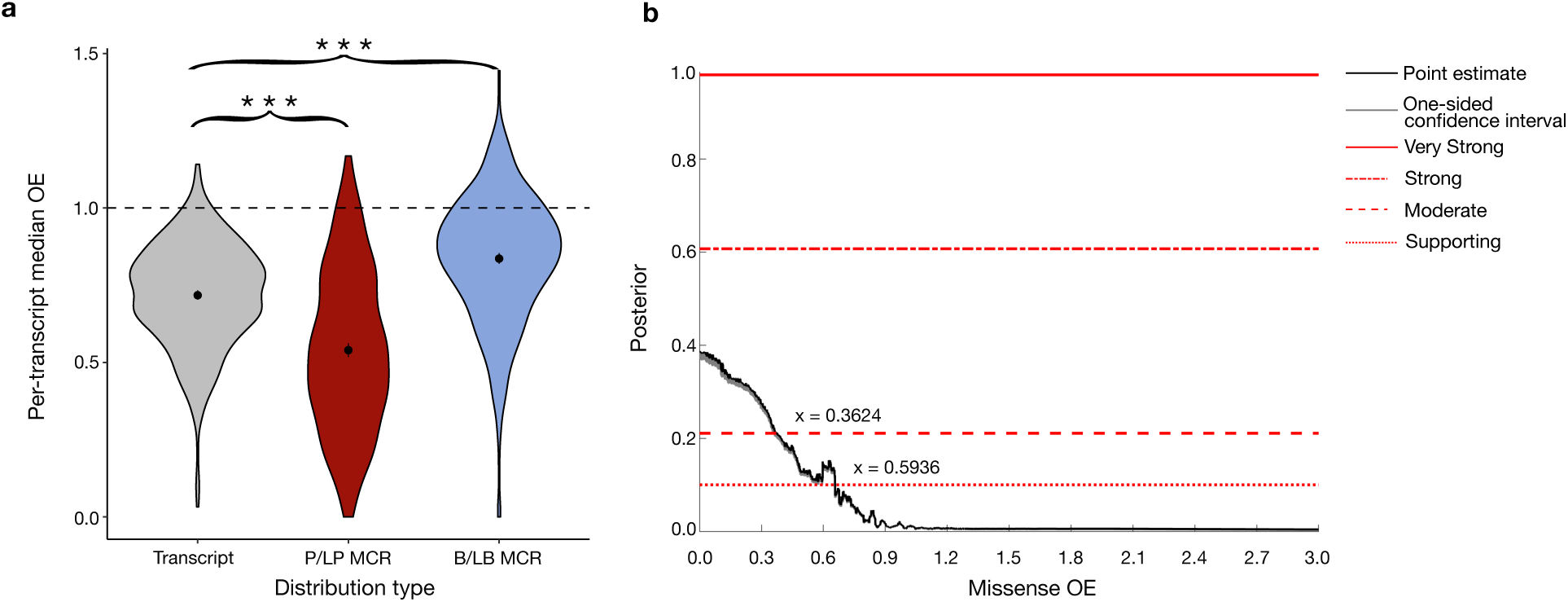
Clinical utility in application of missense constraint. **a,** The distribution of transcript-wide missense observed/expected (OE; gray) and missense constraint region (MCR) OE for ClinVar pathogenic/likely pathogenic missense variants (P/LP; red) and benign/likely benign missense variants (B/LB; blue) in genes with ≥2 MCRs and autosomal dominant disease associations. We filtered to 501 transcripts with at least one P/LP and one B/LB missense variant. For the P/LP and B/LB distributions, we annotated each variant with the missense OE across the MCR they fell in and collapsed these values within each transcript by taking the respective medians.

Having confirmed a correlation between constraint and clinical disease information, we then sought to quantify the utility of our constraint metric as a point of evidence for clinical variant classification. To do so, we leveraged a probabilistic framework established by ClinGen^34^ that formally identifies score thresholds for computational metrics that match each variant classification tier (e.g., “strong” or “moderate”) under the ACMG/AMP guidelines^33^. We applied this robust quantitative framework to determine the missense OE thresholds that met different levels of clinical evidence strengths evaluated under the population data-derived pathogenic (PP2) and benign (BP1) criteria codes^33^. We did not apply a restriction on gnomAD coverage for this assessment, as individuals leveraging our constraint metric for clinical interpretation may not make such considerations, though we do provide coverage information and scores are coverage-adjusted (see **Supplementary Note**). The resulting thresholds are thus conservative estimates. We discovered that our constraint metric met moderate (OE ≤ 0.36) and supporting (OE ≤ 0.59) evidence for pathogenicity, as well as supporting (OE > 0.97) and moderate (OE > 1.23) evidence for benignity (**Fig. 4b, Supplementary Fig. 12**). These correspond respectively to the 3rd, 9th, 65th, and 99th percentiles of the overall missense OE distribution across coding sites. Similar assessment of another regional constraint metric, the missense tolerance ratio (MTR)^62^, revealed that MTR also reaches supporting and moderate support for pathogenicity, but does not meet any level of evidence to support benignity (**Supplementary Fig. 12, Supplementary Table 13**). These results establish formal thresholds on our missense constraint metric that may be used within the ACMG/AMP clinical variant classification guidelines.

Asterisks denote p < 10^−50^. **b,** Local posterior probabilities of variant pathogenicity given MCR missense OE calculated across all transcripts. Gray shading indicates the one-sided 95% confidence interval on the more stringent side. Horizontal lines indicate thresholds required to meet ACMG/AMP evidence levels — from bottom to top: supporting, moderate, strong, very strong. MCR missense OE reaches supporting (OE ≤ 0.59) and moderate (OE ≤ 0.36) level evidence for the population data-derived pathogenic (PP2) criteria code.

### Integrating missense constraint into a variant-level deleteriousness score

We integrated our regional missense constraint measure into a variant-level predictor of missense deleteriousness named MPC (Missense deleteriousness Prediction by Constraint) that additionally incorporates information about amino acid substitution type and local context.

The XGBoost-based model integrates regional and gene constraint-derived metrics together with PolyPhen-2^63^ and phyloP^61^ and is trained on ClinVar pathogenic variants in 2,987 haploinsufficient genes^64^ and 359 genes with non-LoF DD associations in G2P^60^ vs. ClinVar benign variants and gnomAD variants with AF > 0.1% (**Supplementary Tables 14-15, Supplementary Fig. 13**). Higher scores predict greater deleteriousness (**Supplementary Figs. 14-15**).

We assessed the utility of MPC in prioritizing potentially disease-causing variation by evaluating its ability to stratify case and control rare and *de novo* missense variation. Consistent with the regional constraint results, the *de novo* missense variants in DD and AUT are enriched for high MPC scores compared to the control siblings without DDs (**Supplementary Fig. 16**). We then calculated burden enrichments in DD and AUT compared to the controls, stratifying by presence in 373 genes previously associated with NDD^16^ and three bins of MPC scores (<2, 2-2.5, ≥2.5; see **Supplementary Note** for threshold selection; **Fig. 5a-b**, **Supplementary Fig. 17, Supplementary Table 16**). We found a very strong enrichment of *de novo* variants in NDD-associated genes for most MPC bins (from most- to least-damaging MPC: rate ratios [RRs] = 5.1, 3.9, 1.2 for AUT; 15.1, 8.0, and 1.4 for DD), consistent with known causal genetic architectures for these phenotypes. We additionally discovered meaningful enrichments of predicted-damaging *de novo* variants in genes not yet associated with NDDs. For AUT, this was only significant in the most severe MPC bin (RR = 1.7, p = 1.3×10^−5^), suggesting a sizable reservoir of *de novo* missense variants causal for autism in as-of-yet unassociated genes that will likely require strict deleteriousness criteria to statistically ascertain. Substantial *de novo* enrichments were found outside of NDD genes for DD across all missense deleteriousness bins (RRs = 3.6, 1.4, and 1.1), pointing to considerable latent risk outside of currently associated genes and even outside of known OMIM genes (RR for MPC ≥2.5 = 2.2, p = 2.0×10^−9^).

**Fig. 5:**
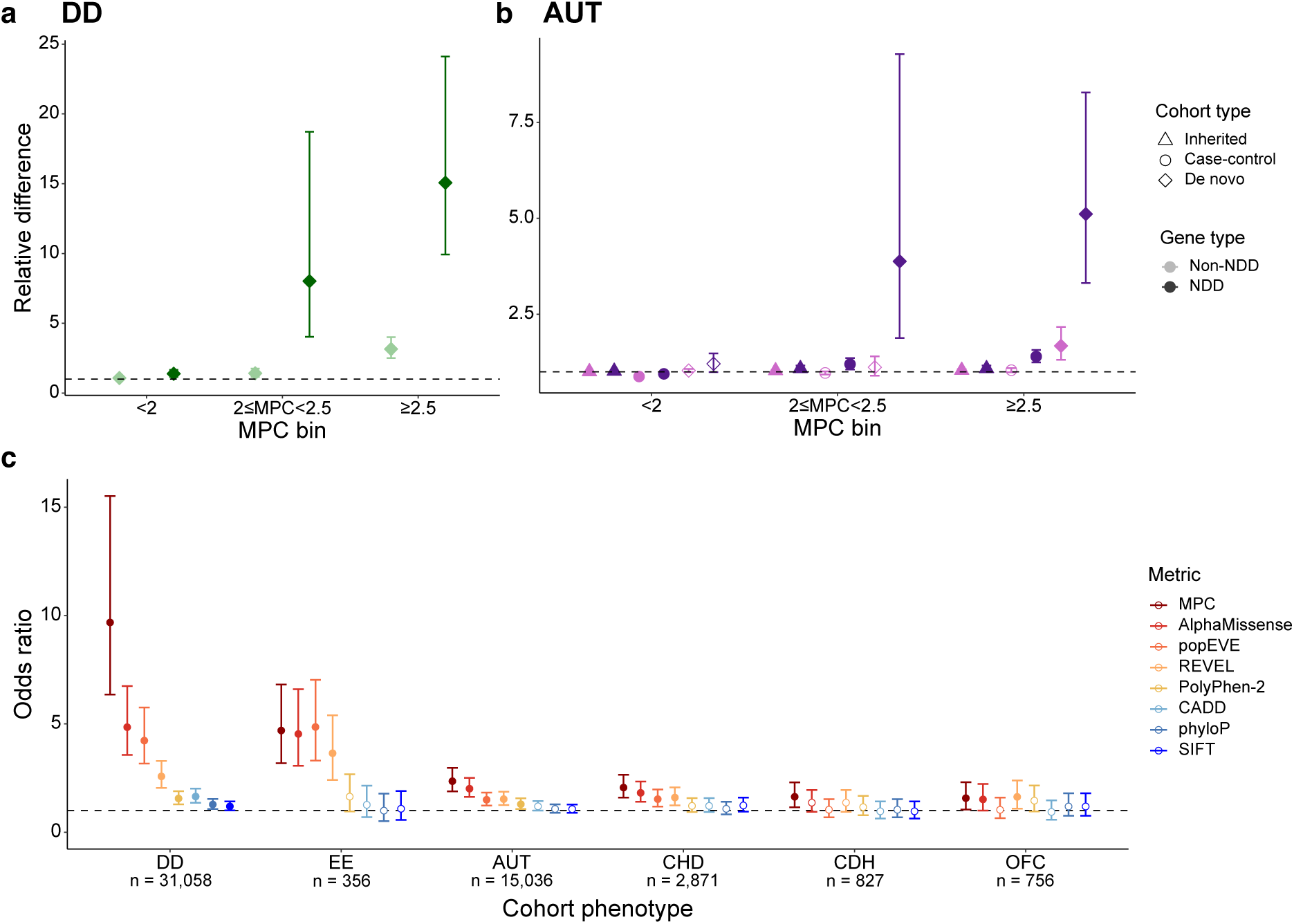
MPC effectively stratifies case and control variation. **a**, The difference relative to controls of missense variants stratified by MPC score and localization to genes associated with neurodevelopmental disorders (NDDs) for **a,** individuals with a developmental disorder (DD) and **b**, autistic individuals (AUT). Relative difference is calculated as: for *de novo* variants, the average rate of variants in probands divided by that in sibling controls; for case-control, the average rate of variants in cases divided by that in controls from case-control data; for inherited, the average rate in probands of transmitted variants divided by that of untransmitted variants. Error bars represent 95% confidence intervals calculated from a binomial test. **c**, The odds ratio of case to control *de novo* missense variants in the top 5% vs. bottom 95% of respective rankings. *De novo* missense variants from each case cohort are ranked against those in the 5,492 controls for each predictor. EE: epileptic encephalopathy, OFC: orofacial cleft, CHD: congenital heart disease, CDH: congenital diaphragmatic hernia. Error bars represent 95% confidence intervals. Only variants scored by all predictors are included. In all plots, points are solid colored if the difference from 1 is statistically significant (binomial or Fisher’s exact p < 0.05).

We additionally assessed enrichments of rare missense variants in 5,591 autistic individuals and 8,597 controls from case-control studies without *de novo* information, as well as parental transmission for 13,384 autistic individuals^16^. While we did not find meaningful parental transmission differences for any MPC bin, we did observe substantial enrichment in the case-control analysis for variants in NDD genes in the two most damaging MPC bins (RRs = 1.4 and 1.2, p = 9.8×10^−9^ and 3.8×10^−3^, respectively), most likely from *de novo* variants. No significant case-control enrichment was found in NDD-unassociated genes, pointing to the importance of *de novo* information for autism gene discovery.

We extended our assessment of case-control *de novo* stratification for a comparison of our model against several other *in silico* missense deleteriousness predictors: AlphaMissense^40^, popEVE^41^, REVEL^39^, Polyphen-2^63^, CADD^65,66^, phyloP^61^, and SIFT^67^. For this assessment, we evaluated four additional early-onset developmental phenotypes: epileptic encephalopathy (EE)^68^, orofacial cleft (OFC)^69^, congenital heart disease (CHD)^70^, and congenital diaphragmatic hernia (CDH)^71^. To compare across predictors with different score distributions, we used a ranking-based performance assessment. For each predictor, we ranked the *de novo* missense variants from each case cohort against those in the 5,492 controls without DDs from the autism study and computed the case-control variant ORs in the top percentiles of these rankings (**Fig. 5c, Supplementary Table 17**). At the top 5% of variants, MPC displays the highest OR for DD (OR = 9.7, Fisher’s exact p < 10^−50^), AUT (OR = 2.4, p = 1.6×10^−16^), CHD (OR = 2.1, p = 1.4×10^−8^), and CDH (OR = 1.6, p = 4.8×10^−3^), demonstrating that MPC effectively ranks high-impact *de novo* variants in the most deleterious prediction regimes. Of the other predictors, AlphaMissense, popEVE, and REVEL tend to perform best. There is generally substantial overlap in the 95% confidence intervals between the top-performing predictors, indicating that *de novo* variant stratification performance tends to be fairly comparable across those models. The exception is in DD, where MPC has much higher performance than the other predictors.

This is likely driven by the way MPC is designed to predict deleteriousness, which focuses on strong fitness effects (see **Discussion**). These observations are generally consistent over a range of thresholds used to define the top percentiles for ranking (**Supplementary Fig. 18**). Across the phenotype categories assessed, the predictors typically had the highest efficacy in DD and EE, then AUT and CHD, and finally CDH and OFC, consistent with the expected contribution of Mendelian *de novo* missense variants to relative risk for each condition. In the top 1% ranking, CDH and OFC missense enrichments reach substantial suggestive enrichments, though not Bonferroni significant (CDH OR = 2.8 and p = 3.8×10^−3^, OFC OR = 3.7 and p = 4.7×10^−4^), indicating that while *de novo* missense variants contribute to risk for these disorders, they do so at a lower rate than for more severe phenotypes.

## Discussion

We developed a method to identify sub-genic regions with differential intolerance to missense variation. We demonstrate that coding regions depleted for missense variation in the general population are enriched for established disease-associated variation, *de novo* variants from individuals with developmental disorders, and heritability for 268 complex traits from the UK Biobank and other large GWAS. Additionally, we have calibrated these constraint scores to establish that regions with less than 36% of their expected missense variation can achieve moderate evidence towards clinical classification of pathogenicity for missense variants that occur in these regions, following ACMG/AMP guidelines. Finally, we incorporated regional missense intolerance information into the missense deleteriousness metric, MPC, and show that MPC effectively separates potentially risk-carrying variants identified in various developmental disorder cases from those seen in controls.

Our method of summarizing constraint strength information over contiguous base pair sites into discretized regions allows for identification of gene segments under distinct selection pressures that can point to different functions and structures within a gene. This approach can provide more easily interpretable information about broad gene functions than very fine resolution metrics at the site- or amino acid-level. For example, we show that MCRs, which do not incorporate any protein structural information, can align precisely with the locations of protein domains and functional intrinsically disordered regions as well as the pathogenic disease variants therein.

Other metrics that have been developed to identify gene segments with distinct missense intolerance linearly along the genome include subRVIS^21^, Constrained Coding Regions (CCRs)^27^, and the constrained segments calculated with MTR^62^, among others. SubRVIS is limited to quantification at the level of exons or domains, lacking dynamic ascertainment of functional gene regions. The CCR metric is generally more akin to a site-level metric: 60% of CCR gene segments are 1bp in size due to being the site of a nonsynonymous variant observed in gnomAD (assigned a score of 0 or unconstrained), and 0.1% of segments are >50bp.

Notably, both subRVIS and CCRs were calculated on datasets much smaller than the dataset underlying the MCRs presented here (gnomAD v4.1.1): subRVIS used the Exome Sequencing Project^72^ (∼112x smaller than gnomAD v4.1.1) and CCRs used gnomAD v2.1.1 (∼6x smaller than v4.1.1). We previously provided a comparison of an earlier version of MCRs calculated with gnomAD v2.1.1^73^ to CCRs. MTR was calculated on the Regeneron Genetics Center Million Exome (RGC-ME) dataset consisting of 821,979 unrelated samples, which is a comparably sized dataset to gnomAD v4.1.1 and should therefore afford similar statistical power to our MCR method. The RGC-ME study released MTR intolerance scores at the amino acid level as well as at the gene segment level, the latter calculated by aggregating over the former. We find that the

MTR method of segmenting genes into regions is prone to identifying likely spurious “unconstrained” regions as there are no restrictions placed on the size of unconstrained regions, only constrained regions. This results in approximately 16% of unconstrained regions falling under the 10 amino acid minimum applied to constrained regions. Furthermore, the binary nature of the assignment of amino acids as “constrained” or “unconstrained” for input to segmentation means that there is no distinction between regions of varying levels of constraint below the 15th percentile threshold designated “constrained”. As a result, consecutive regions that fall under the “constrained” threshold but have different levels of constraint will be unlikely to be distinguished from each other by the MTR method unless an “unconstrained region” is identified that separates them. In contrast, our MCR method delineates gene segments based on having differing levels of constraint without this binarization, and the size minimum of 16 expected variants is applied to all MCRs for robustness.

We note that constraint, as quantified here by population depletion of observed rare variants, is essentially a footprint of selection on heterozygous variants. Our methodology is thus most powerful in sequences undergoing appreciable heterozygous selection, e.g., genes where severe and early onset diseases are caused by dominant missense variants. However, as demonstrated in **Supplementary Fig. 11**, disease variants in genes currently known to be only associated with conditions with recessive inheritance do also associate with detectable constraint, albeit to a lesser degree than dominant variants. This lends further support to a body of evidence that few genes are purely recessive: many “recessive-only” genes are likely only partially recessive, with heterozygotes having measurable phenotypic differences^74–76^. As such, we show that this constraint method is indeed amenable, to an extent, to characterizing selective effects even in genes considered to be solely associated with recessive inheritance.

An inherent strength of our regional missense constraint metric is its grounding in population genetics and independence of, and thus lack of bias from, disease genetics databases and domain or structural knowledge. Given that constraint measures fitness effects, we expect—and indeed observe—associations between strong constraint and risk variants in early developmental disorders which are under strong selective pressure (**Fig. 2a**, **Fig. 3b**), while genes where germline variants cause adult-onset diseases like hereditary cancers generally harbor little detectable constraint on missense or pLoF variants. This makes it all the more notable when we clearly identify highly constrained gene sequences where clinically relevant missense variants are currently only known to be associated with adult-onset disease, such as in *TARDBP* (**Fig. 2b**), where the highly constrained C-terminal MCR bounds nearly all clinically reported variants causing ALS and FTD, diseases that typically begin in post-reproductive life but can start in earlier adulthood^77–80^. The rest of the gene is even more strongly constrained, which points to severe fitness effects unexplainable by the known gene-disease associations, demonstrating the value of this constraint method in revealing novel evidence for gene function. Additionally, while missense predictive models that rely on 3D structure exclude or have lower performance in intrinsically disordered regions^81–83^, there is no such structure-based incompatibility for the genetics-derived inference of MCRs. Returning to *TARDBP* as an example, nearly all of the 23 P/LP missense variants in ClinVar lie in the disordered regions of this gene. Of the 23, AlphaMissense provides support for pathogenicity for only 7 while supporting benignity for the remaining 16 (using the developer-recommended score thresholds of >0.564 and <0.34, respectively^40^; the average score for all 23 is 0.36). Meanwhile MPC, where missense constraint is a key component, predicts scores >2 for all 23 variants (average score of 2.76 or 97th percentile). Still, detachment from structural information means that MCR identification is restricted to a linear search along the genome, and we are thus less powered to discover 3D-specific constraint clusters such as small binding pockets.

For the MPC variant-level deleteriousness predictor, certain protein domain and structural information is learned with the PolyPhen-2 model feature, though this has limitations compared to full 3D structure-based methods like AlphaMissense. We thus evaluated the utility of combining MPC and AlphaMissense in predicting missense deleteriousness. We re-categorized the DD, AUT, and control *de novo* missense variants by whether they were predicted to have high or low deleteriousness scores by MPC and AlphaMissense (≥2.5 and <2 for MPC, ≥0.9985 and <0.761 for AlphaMissense; see **Supplementary Note** for threshold selection) and calculated case-control enrichments. Variants predicted to be highly deleterious by both MPC and AlphaMissense have stronger enrichment than when predicting by either metric alone, and variants where deleteriousness is high in one metric and low in the other also harbor detectable enrichments (**Supplementary Fig. 19, Supplementary Table 18**). This demonstrates that MPC and AlphaMissense offer complementary information and joint application can improve prediction performance. However, we caution against taking the simple maximum of the MPC or AlphaMissense relative deleteriousness for any variant — this sharply decreases predictive performance in ranking case vs. control *de novo* variants (**Supplementary Fig. 18, Supplementary Table 17**), arising from decreased specificity wherein MPC and AlphaMissense each assign relatively high predicted deleteriousness to subsets of the control *de novo* variants.

Lastly, to guide usage of the MPC model, we note that MPC is best suited to modeling strong fitness effects: the variants composing the deleterious training set were chosen to be those likely under strong selection given constraint and disease association evidence, and the missense constraint feature of the MPC model is, as aforementioned, grounded in heterozygous selection. This is demonstrated by MPC’s excellent performance in distinguishing *de novo* variants in a severe DD cohort from controls (**Fig. 5c**), which is driven by a combination of model design and unavoidable training biases as the training set naturally includes known DD genes that severely disrupt development.

In summary, we identify 36% of MANE Select or canonical transcripts with variable levels of missense constraint and demonstrate that coding regions specifically depleted of missense variation in the general population are enriched for disease-associated variation. Additionally, we show that this depletion of missense variation provides moderate strength evidence when clinically classifying variants according to ACMG/AMP guidelines and that incorporation of regional missense constraint into an *in silico* predictor effectively prioritizes a subset of *de novo* missense variation in individuals with developmental phenotypes for association testing. We have publicly released these data for use in both research and clinical settings. We anticipate refined resolution of these metrics as datasets grow, both in size and in ancestral representation, and with the incorporation of complementary structural or functional data.

## Methods

### Transcripts

This study followed the same steps of transcript annotation and the Ensembl Variant Effect Predictor (VEP) table as described in gnomAD v4.1.1^50^ and analyzed only coding transcripts as defined by MANE Select v0.95 and, when not present, the canonical transcript as listed in GENCODE v39/Ensembl VEP v105. For high quality transcripts, we excluded transcripts that had outlier variant counts: zero expected or too many observed pLoF, missense, or synonymous variants; or too few observed synonymous variants. This totaled 17,841 transcripts, 96% from MANE Select and 4% canonical. We also make MCR and MPC information available for the 1,534 transcripts with outlier counts, but caution that scores may be less accurate in these sequences.

### Genome Aggregation Database (gnomAD) population data

All analyses in this paper were conducted using the 730,947 gnomAD v4.1.1 exomes on GRCh38^50^. Coverage for gnomAD v4.1.1 was calculated using information from sample genomic VCF (gVCF) data rather than from read data. As a result, coverage information for gnomAD v4.1.1 is not as granular due to the reference block structure within gVCFs. To remedy this, we used allele number percent (%AN) to proxy coverage for all analyses to improve our ability to capture constraint in lower-coverage sites. Sites with %AN < 20 (0.68% of all possible missense sites with %AN > 0) were excluded from analysis. “High-coverage” sequences refer to MCRs with %AN ≥ 90. The gnomAD allele frequencies used in this study are those calculated across all individuals. LOEUF and transcript missense OE scores are from the gnomAD v4.1.1 gene constraint data^50^.

### ClinVar variants

We annotated functional consequences for ClinVar^8^ (v.20250504) variants using the Ensembl VEP table. For this study, we selected high-quality missense ClinVar variants with non-conflicting pathogenic (P), likely pathogenic (LP), variant of uncertain significance (VUS), benign (B), or likely benign (LB) classification and a review status of at least one star. For these high-quality missense variants, 89.9% are VUS (1,585,522/1,764,005).

### OMIM data

We used OMIM (v.20230215) to annotate gene-phenotype modes of inheritance (MoI). Genes were considered autosomal dominant if their only MoI noted in OMIM was autosomal dominant, likewise for autosomal recessive. In total, 1,036 genes were considered autosomal dominant, and 2,176 were considered autosomal recessive.

### Gene2Phenotype (G2P) data

Phenotype associations and mechanistic information for developmental disorder (DD) genes were taken from the G2P DD panel (v.20251127). We filtered to 810 genes with definitive or strong evidence and a monoallelic mechanism for DD association. The 359 genes considered to be non-LoF were those where the noted variant consequence included “altered gene product structure” or “increased gene product level”, and the noted molecular mechanism was not loss-of-function.

### Missense Tolerance Ratio (MTR) scores

We obtained MTR scores estimated from the exomes of 983,578 individuals^62^. This method provides a window-based estimate of observed/expected missense variant counts, similar to the gnomAD missense OE presented here. Scores are available at the amino acid-level (smoothed across a window of 31 amino acids), or a segment (region) level calculated by aggregating consecutive amino acids below a certain “constrained” OE threshold. For the analysis calibrating metric thresholds for clinical interpretation, we used the amino acid-level scores.

### Rare and *de novo* variants from developmental disorder (DD) cohorts

Case *de novo* variants for association analyses were obtained from studies of severe DD^15^, autism^16^ (AUT), congenital heart disease^70^ (CHD), orofacial cleft^69^ (OFC), congenital diaphragmatic hernia^71^ (CDH), and epileptic encephalopathy^68^ (EE). Control *de novo* mutations were obtained from siblings of the autistic probands without DDs^16^. Variants were lifted over from GRCh37 to GRCh38 using the “liftover” function in Hail for the DD, CDH, and EE cohorts. Variant functional consequences were re-annotated using the Ensembl VEP table. Variants transmitted and not transmitted from parents to autistic probands were procured from previously published ASC-SSC and SPARK cohorts, and case-control variants for autism were procured from previously published iPSYCH and Swedish cohorts^16^. Both the transmitted/untransmitted and case-control variant sets were filtered to gnomAD AF < 0.1%.

### Identifying breakpoints within transcripts of regional missense constraint

We defined the set of sites with possible missense variants using a synthetic Hail Table (HT) containing all possible single nucleotide variants in the exome. We annotated this HT with the Ensembl VEP table and filtered it to variants with the consequence “missense_variant” in the MANE Select or canonical coding transcripts as defined in *Transcripts*. We then further filtered to (1) variants observed in gnomAD with allele count (AC) > 0, AF < 0.001, %AN ≥ 20, and QC PASS; or (2) variants unobserved in gnomAD with AC = 0. We then aggregated the total number of observed variant counts per base pair site by summing the number of observed variants over each possible substitution (reference and alternate allele) at the locus.

As described in more detail for the gnomAD v4.1.1 release^50^, we applied two models (plateau and coverage) to calculate the proportion of expected missense variation per variant (chromosome/locus plus reference and alternate alleles). As with the observed missense counts, we then summed the number of expected variants to get the total expected rare missense variant count per locus. We applied a likelihood ratio test to determine whether the missense observed/expected (OE) ratio was uniform along a transcript or whether a transcript had evidence for discrete subsections (regions) with distinct missense constraint strengths. We assumed that the observed missense counts should follow a Poisson distribution around the expected missense counts. We defined our null model as transcripts not having any evidence of regional variability in missense depletion (where the OE ratio is consistent across the length of the transcript). Our alternative model was that transcripts exhibited evidence of distinct subsections of missense depletion (OE ratio calculated per transcript subsection). Because the alternative model should always have a better fit than the null model, we require a chi-squared value above a given threshold (p = 0.001) to establish significance. We used the following formulas to determine the significance of a breakpoint that would split a transcript into two subsections, A and B:

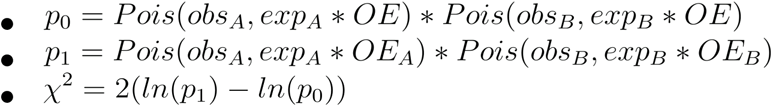

where OE is the missense observed/expected ratio across the entire transcript, obs_A_ is the number of observed missense variants in transcript subsection A, exp_A_ is the number of expected missense variants in transcript subsection A, OE_A_ is the OE ratio across transcript subsection A, obs_B_ is the number of observed missense variants in transcript subsection B, exp_B_ is the number of expected missense variants in transcript subsection B, OE_B_ is the OE ratio across transcript subsection B, and Pois is the Poisson likelihood.

Likewise, we used the following formulas to determine the significance of a breakpoint that would split a transcript into three subsections, A, B, and C:

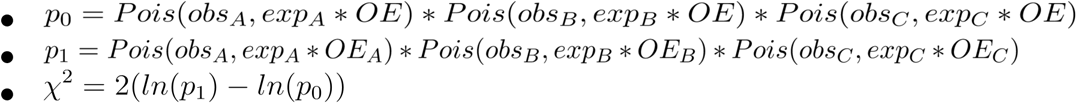

For the purposes of this breakpoint search, all transcript subsections with more observed variants than expected were capped at an OE of 1, as we were looking for areas of missense depletion and not missense enrichment. We also converted the expected counts for transcript subsections with zero expected variants from 0 to 10^−9^ to prevent nonfinite OE values. We implemented a minimum number of expected missense variants (16) to prevent identification of breakpoint positions that would create very small (i.e., a handful of base pairs in size) transcript subsections that are more likely to be spurious (see **Supplementary Note**).

To search for a single breakpoint that would divide a transcript into two subsections, we calculated chi-squared statistics to conduct likelihood ratio tests simultaneously for every eligible position within a transcript. The positions we considered were base pair sites with a possible missense variant substitution that had at least 16 expected missense counts in either direction (i.e., both transcript subsections created by dividing the transcript at this point would have at least 16 expected missense variants). Breakpoints were determined by the site with the maximum chi-squared value that was significant at p = 0.001. Any transcripts that did not have a single significant breakpoint were searched for two simultaneous breakpoints that would divide the transcript into three sections. As in the single-breakpoint scenario, we tested all pairs of eligible base pair sites where the resulting transcript subsections would harbor at least 16 expected missense variants. For transcripts where we identified significant breakpoints, we repeated this process (single-breakpoint search, then two-breakpoint search) recursively on each subsection until no further significant breakpoints were found.

### Clinical calibration of missense constraint regions

We filtered to genes that had at least one pathogenic missense variant in ClinVar and selected missense variants in those genes with a non-conflicting pathogenic (P), likely pathogenic (LP), benign (B), or likely benign (LB) classification and a review status of at least one star. Missense variants with an AF ≥ 1% in gnomAD v4.1.1 were removed, leaving a total of 122,269 ClinVar variants in 4,334 genes. For this set of genes, we also retrieved QC PASS variants with AF < 1% from gnomAD v4.1.1, which resulted in 3,847,501 gnomAD variants from 4,302 genes.

All variants were annotated with MCR missense OE and the amino acid-level missense tolerance ratio (MTR) metric^62^. Using our previously developed method^34^ and the prior probability of pathogenicity of 4.41%, we computed the local posterior probability curves for all scores. P/LP and B/LB ClinVar variants were used to estimate the local values of the posterior probability of pathogenicity, and gnomAD variants were used to smooth out the posterior estimates. We defined score threshold ranges for each strength of evidence (supporting, moderate, strong, and very strong) for pathogenicity and benignity based on the ACMG/AMP criteria (**Supplementary Table 13**). For both metrics, we plotted the local posterior probability curve estimate along with the one-sided 95% confidence interval, determined using 10,000 bootstrapping iterations.

### Modeling deleteriousness of individual missense variants

We designed our missense variant deleteriousness predictor (Missense deleteriousness Prediction by Constraint or MPC) to explicitly incorporate information on amino acid substitution class and position-specific variant effects informed by missense constraint. Briefly, we trained a machine learning model to differentiate “pathogenic” missense variants likely under strong heterozygous selection from “benign” missense variants likely under neutral to near-neutral selection. The “pathogenic” training set consisted of high-quality ClinVar variants (see *ClinVar variants*) labeled as P or LP in 2,987 predicted-haploinsufficient genes (defined as having probability of haploinsufficiency (pHaplo) ≥ 0.86^64^) or in 359 genes with DD associations in G2P through non-LoF mechanisms (see *Gene2Phenotype data*). The “benign” training set consisted of high-quality gnomAD variants AF > 0.1% or ClinVar B/LB variants in the same genes as the “pathogenic” set (see *gnomAD variants* and *ClinVar variants*). Variants matching criteria for both the “benign” and “pathogenic” sets were removed from the training data. We further removed variants with %AN < 20 in the gnomAD v4.1.1 exomes and variants that were not in the 17,841 MANE Select/canonical transcripts.

We evaluated logistic regression and XGBoost (gradient-boosted tree) architectures, incorporating features of MCR and gene constraint, amino acid substitution class severity, PolyPhen-2 for orthogonal information from protein homology and structure, and phyloP for evolutionary selection information across longer timescales. Only variants with all relevant annotations were retained for training and for the full variant score table. The XGBoost model was chosen over logistic regression based on performance on a hold-out gene set and then retrained on all data (20,931 “pathogenic” variants and 93,638 “benign” variants) for the final model. We applied the XGBoost model across the 70,313,598 possible exome-wide missense variants in the Ensembl VEP table with all features to obtain a probability of pathogenicity for each variant. The MPC score for any missense variant *i* is given as:

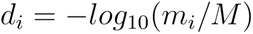

where *d_i_* is the MPC score, *m_i_* is the number of “benign” missense variants in the training set with a fitted probability of pathogenicity value that is less than the fitted value for variant *i*, and *M* is the total number of “benign” missense variants in the training set. When *m_i_* is 0, i.e. when the fitted score for variant *i* is more severe than for all “benign” variants, this introduces a log-zero error so we set *d_i_* to 6 (the maximum real value for *d_i_* is just over 5 when *m_i_* is 1). Larger values of *d_i_* indicate stronger predicted-deleteriousness. See **Supplementary Note** and **Supplementary Figs. 13-16** for further detail on these methods and the distribution of MPC scores.

### Comparison of MPC to other predictors

We compared our model to the following *in silico* missense deleteriousness predictors: AlphaMissense^40^, popEVE^41^, REVEL^39^, CADD^65,66^, PolyPhen-2^63^, and SIFT^67^. AlphaMissense scores were downloaded from https://github.com/google-deepmind/alphamissense and mapped to variants by locus and gene name, as the transcript provided was not always the MANE Select transcript. popEVE scores were downloaded from https://pop.evemodel.org. REVEL scores were downloaded from https://sites.google.com/site/revelgenomics/. PolyPhen-2 and SIFT scores were obtained from VEP^84^. CADD scores were downloaded from the CADD website (https://cadd.gs.washington.edu/download). phyloP scores were downloaded from the UCSC browser (https://genome.ucsc.edu/cgi-bin/hgTrackUi?db=hg38&g=cons241way). We annotated the case and control *de novo* missense variants described in *Rare and de novo variants from developmental disorder cohorts* and ranked the variants based on their annotated scores. To assess each predictor’s ability to stratify case and control variation, we assessed the proportion of case to control variants among the variants with the top 1%, 5%, or 10% for each score and compared to the remaining 99%, 95%, and 90% of variants, respectively, using Fisher’s exact test.

## Supporting information

Supplementary Tables 1 through 18

## Data availability

The missense constraint regions (MCRs) are displayed on the gnomAD v4 browser (https://gnomad.broadinstitute.org) and available for download on the gnomAD website (https://gnomad.broadinstitute.org/data#v4) and in the gnomAD v4 public datasets on Google and Amazon clouds (gs://gcp-public-data--gnomad/papers/2026-rmc). MPC scores for all possible variants in MANE Select canonical transcripts are available in the gnomAD v4 public datasets on Google (gs://gcp-public-data--gnomad/papers/2026-rmc/gnomad_v4.1.1_mpc.ht). gnomAD v4 exome, genome, and allele number data and the table of all possible single nucleotide polymorphisms used to calculate mutational models and search for MCRs are also available in the gnomAD public buckets and are accessible using code in the gnomAD Hail utilities GitHub repository (https://github.com/broadinstitute/gnomad_methods/blob/b61aff692041dbf1118773b18104f13d1 31008fa/gnomad/resources/grch38/gnomad.py#L456 and https://github.com/broadinstitute/gnomad_methods/blob/b61aff692041dbf1118773b18104f13d1 31008fa/gnomad/resources/grch38/reference_data.py#L147).

ClinVar data were downloaded from ClinVar’s FTP server (https://ftp.ncbi.nlm.nih.gov/pub/clinvar/). *De novo* variants were extracted from the supplementary files of the cited studies.

## Code availability

Code to determine missense constraint regions (MCRs) and calculate MPC is available at https://github.com/broadinstitute/regional_missense_constraint. Code used to generate the mutational models is available at https://github.com/broadinstitute/gnomad_methods/blob/main/gnomad/utils/constraint.py and https://github.com/broadinstitute/gnomad-constraint. Code used to generate figures is available at https://github.com/broadinstitute/rmc-2026-figures. The Hail library is available at https://hail.is/.

## Acknowledgments

We thank all of the individuals who contributed their data to gnomAD for enabling the research presented here and in numerous other publications. We also thank members of the gnomAD Production and Browser Teams for their behind-the-scenes efforts in producing and sharing the gnomAD dataset, Dan King and the rest of the Hail team for enabling data processing at scale, and Emma Pierce-Hoffman for her work curating gene lists. This work was supported by the National Human Genome Research Institute (NHGRI; U24HG011450 to H.L.R. and M.J.D.; U01HG011755 to A.O.D.L., H.L.R, and M.E.T.; U24HG006834 to H.L.R, U01HG012022 to P.R.; 5T32HG002295 to L.W.; R01HG012867 to K.E.S.), the Eunice Kennedy Shriver National Institute of Child Health and Human Development (NICHD; F31HD111109 to L.W.), the National Science Foundation (NSF; GRFP#2022339661 to L.W.; GRFP#2023361442 to H.A.), the Simons Foundation (SFARI 1009802 to M.E.T. and K.E.S.), a National Health and Medical Research Council investigator grant (2009982) to D.G.M., and the Massachusetts General Hospital Executive Committee on Research, Interim Support Funding (K.E.S.).

## Author information

These authors contributed equally: Lily Wang and Katherine R. Chao.

## Contributions

L.W., K.R.C., K.E.S., B.M.N, and M.J.D conceived and designed experiments. L.W., K.R.C, and K.E.S. performed primary writing of the manuscript. L.W., K.R.C., R.P., C.L., H.A., and R.Y. performed the analyses and generated figures. P.S. and J.C. were instrumental to developing methods. R.H.G., N.A.W., and M.S. developed visualizations for the web browser. B.W., W.P., M.W.W., K.M.L., J.K.G, K.J.K, and G.T. completed code review for methods. D.G., J.I.G., C.V., and T.P. helped debug runtime compute. J.A.K. provided data and analysis suggestions. S.B. contributed analysis suggestions. H.L.R., B.M.N., M.E.T, D.G.M, A.O.D.L., K.J.K., P.R., M.J.D., and K.E.S. supervised the research. All authors listed under the Genome Aggregation Database Consortium contributed to the generation of the primary data incorporated into the gnomAD resource. All authors reviewed the manuscript.

## Genome Aggregation Database Consortium

Maria Abreu^1^, Amina Abubakar^2,3^, Rolf Adolfsson^4^, Carlos A. Aguilar Salinas^5^, Tariq Ahmad^6^, Elissa Alarmani^7^, Christine M. Albert^8,9^, Jessica Alföldi^7,10^, Matthieu Allez^11^, Irina M. Armean^7,10^, Elizabeth G. Atkinson^12,13^, Gil Atzmon^14,15^, Eric Banks^16^, John Barnard^17^, Daniel Ben-Isvy^7,18^, Emelia J. Benjamin^19,20,21^, David Benjamin^16^, Louis Bergelson^16^, Charles Bernstein^22,23^, Douglas Blackwood^24^, Michael Boehnke^25^, Erwin P. Bottinger^26^, Donald W. Bowden^27^, Matthew J. Bown^28^, Harrison Brand^18,29^, Steven Brant^30,31,32^, Ted Brookings^16,33^, Sam Bryant^10,34^, Shawneequa L. Callier^35,36^, Hannia Campos^37,38^, John C. Chambers^39,40,41^, Juliana C. Chan^42^, Sinéad Chapman^7,10,43^, Daniel I. Chasman^8,44^, Lea A. Chen^45^, Siwei Chen^7,10^, Rex Chisholm^46^, Judy Cho^26^, Mina K. Chung^47^, Wendy K. Chung^48^, Kristian Cibulskis^16^, Bruce Cohen^49,50^, Ryan L. Collins^7,18,51^, Kristen M. Connolly^52^, Adolfo Correa^53^, Aiden Corvin^54^, Miguel Covarrubias^16^, Nick Craddock^55^, Beryl B. Cummings^7,51^, Dana Dabelea^56^, John Danesh^57^, Dawood Darbar^58^, Philip W. Darnowsky^7^, Stacey Donnelly^59^, Richard H. Duerr^60,61,62^, Ravindranath Duggirala^63^, Josée Dupuis^64,65^, Patrick T. Ellinor^7,66^, Roberto Elosua^67,68,69^, Eleina England^7,70^, Jeanette Erdmann^71,72,73^, Tõnu Esko^74^, Emily Evangelista^7^, Yossi Farjoun^75^, Diane Fatkin^76,77,78^, William Faubion^79^, Steven Ferriera^80^, Gemma Figtree^81,82,83^, Jose Florez^44,84,85^, Laurent Francioli^7,10^, Andre Franke^86,87^, Adam Frankish^88^, Jack Fu^7,18,89^, Martti Färkkilä^90,91,92^, Stacey Gabriel^80^, Kiran Garimella^16^, Laura D. Gauthier^16^, Jeff Gentry^16^, Michel Georges^93^, Gad Getz^44,94,95^, David C. Glahn^96,97^, Benjamin Glaser^98^, Stephen J. Glatt^99^, Fernando S. Goes^100^, Clicerio Gonzalez^101^, Leif Groop^102,103^, Sanna Gudmundsson^7,104,105^, Jeremy Guez^7,10^, Michael Guo^7^, Namrata Gupta^7,80^, Andrea Haessly^16^, Christopher Haiman^106^, Craig L. Hanis^107^, James Hanyok^108^, Matthew Harms^109,110^, Qin He^7^, Mikko Hiltunen^111^, Matti M. Holi^112^, Christina M. Hultman^113,114^, Steve Jahl^7,10^, Chaim Jalas^115^, Thibault Jeandet^16^, Mikko Kallela^116^, Diane Kaplan^16^, Jaakko Kaprio^103^, Sekar Kathiresan^18,44,117^, Eimear E. Kenny^118^, Bong-Jo Kim^119^, Young Jin Kim^119^, George Kirov^120^, Zan Koenig^10,43^, Jaspal Kooner^40,121,122^, Pragati Kore^12^, Seppo Koskinen^123^, Harlan M. Krumholz^124,125^, Subra Kugathasan^126^, Juozas Kupcinskas^127^, Nehir E. Kurtas^18^, Soo Heon Kwak^128^, Markku Laakso^129,130^, Nicole Lake^131^, Mikael Landén^113,132^, Matthew Lebo^133,134,135^, Terho Lehtimäki^136,137^, Monkol Lek^131^, James D. Lewis^138^, Cecilia M. Lindgren^139,140,141^, Christopher Llanwarne^16^, Ruth J.F. Loos^26,142^, Edouard Louis^143^, Chelsea Lowther^18^, Wenhan Lu^7^, Steven A. Lubitz^7,66^, Ronald C.W. Ma^42,144,145^, Dara S. Manoach^146,147^, Gregory M. Marcus^148^, Jaume Marrugat^149,150^, Nicholas A. Marston^151^, Daniel M. Marten^7,105^, Alicia R. Martin^7,10,43^, Steven McCarroll^43,152^, Mark I. McCarthy^153,154,155^, Jacob L. McCauley^156,157^, Dermot McGovern^158^, Ruth McPherson^159^, Andrew McQuillin^160^, James B. Meigs^7,44,161^, Olle Melander^162^, Andres Metspalu^163^, Deborah Meyers^164^, Eric V. Minikel^7^, Braxton D. Mitchell^165^, Paul Moayyedi^166,167,168^, Sanghamitra Mohanty^169^, Andrés Moreno Estrada^170^, Nicola J. Mulder^171,172^, “ NA, Joshua N. Nadeau^7^, Aliya Naheed^173^, Andrea Natale^174,175,176^, Saman Nazarian^177,178^, Charles Newton^179,180^, Peter M. Nilsson^181^, Sam Novod^16^, Michael C. O’Donovan^55^, Yukinori Okada^182,183,184^, Dost Ongur^44,49^, Roel A. Ophoff^185,186^, Lorena Orozco^187,188^, Willem Ouwehand^189^, Michael J. Owen^55^, Nick Owen^190^, Colin Palmer^191^, Nicholette D. Palmer^27^, Aarno Palotie^10,43,103^, Mara Parellada^192,193,194^, Kyong Soo Park^128,195^, Carlos Pato^196^, Nancy L. Pedersen^113^, Tina Pesaran^197^, Emma Pierce-Hoffman^7^, Sharon Plon^198^, Danielle Posthuma^199,200^, Ann E. Pulver^201^, Aaron Quinlan^202^, Dan Rader^177,203^, Nazneen Rahman^204^, Andreas Reif^205^, Alex P. Reiner^206,207^, Anne M. Remes^208,209^, Stephen Rich^210,211^, John D. Rioux^212,213^, Samuli Ripatti^59,103,214^, Jason Roberts^215,216^, Elise Robinson^18^, Dan M. Roden^217,218^, Jerome I. Rotter^219^, Guy Rouleau^220^, Christian T. Ruff^151^, Heiko Runz^221^, Marc S. Sabatine^151^, Nareh Sahakian^16^, Andrea Saltzman^7^, Nilesh J. Samani^222,223^, Alba Sanchis-Juan^18^, Akira Sawa^224,225,226^, Jeremiah Scharf^7,18,43^, Molly Schleicher^7^, Heribert Schunkert^227,228^, Sebastian Schönherr^229^, Eleanor G. Seaby^7,230^, Svati H. Shah^231,232^, Megan Shand^16^, Ted Sharpe^16^, Moore B. Shoemaker^233^, Tai Shyong^234,235^, Edwin K. Silverman^236,237^, Moriel Singer-Berk^7^, Jurgita Skieceviciene^127^, Jonathan T. Smith^16^, J. Gustav Smith^238,239,240^, Jordan W. Smoller^43,237,241^, Hilkka Soininen^242^, Harry Sokol^243,244,245^, Rachel G. Son^7^, Tim Spector^246^, David St Clair^247^, Sarah Stenton^7^, Christine Stevens^7,10,43^, Nathan O. Stitziel^248,249,250^, Patrick F. Sullivan^113,251^, Jaana Suvisaari^252^, Yekaterina Tarasova^7^, Kent D. Taylor^219^, Yik Ying Teo^253,254,255^, Kathleen Tibbetts^16^, Ming Tsuang^256,257^, Tiinamaija Tuomi^103,258,259^, Dan Turner^260^, Teresa Tusie-Luna^261,262^, Rachel Ungar^7^, Grace VanNoy^7^, Erkki Vartiainen^263^, Marquis Vawter^264^, Elisabet Vilella^265,266,267^, Gordon Wade^16^, Mark Walker^16^, Qingbo Wang^7,268^, Arcturus Wang^7,10,43^, James S. Ware^7,269,270^, Hugh Watkins^271^, Rinse K. Weersma^272^, Christopher Whelan^7^, Nicola Whiffin^7,273,274^, James G. Wilson^275^, Lauren Witzgall^7^, Ramnik J. Xavier^276,277^, Mary T. Yohannes^7^, Robert Yolken^278^, Xuefang Zhao^7^

^1^University of Miami Miller School of Medicine, Gastroenterology, Miami, USA

^2^Neuroscience Research Group, Department of Clinical Sciences, Kenyan Medical Research Institute, Wellcome Trust, Kilifi, Kenya

^3^Institute for Human Development, Aga Khan University, Nairobi, Kenya

^4^Department of Clinical Sciences, Psychiatry, Umeå University, Sweden

^5^Unidad de Investigacion de Enfermedades Metabolicas, Instituto Nacional de Ciencias Medicas y Nutricion, Mexico City, Mexico

^6^Peninsula College of Medicine and Dentistry, Exeter, UK

^7^Program in Medical and Population Genetics, Broad Institute of MIT and Harvard, Cambridge, MA, USA

^8^Division of Preventive Medicine, Brigham and Women’s Hospital, Boston, MA, USA

^9^Division of Cardiovascular Medicine, Brigham and Women’s Hospital and Harvard Medical School, Boston, MA, USA

^10^Analytic and Translational Genetics Unit, Massachusetts General Hospital, Boston, MA, USA

^11^Gastroenterology Department, Hôpital Saint-Louis - APHP, Université Paris Cité, INSERM U^1160^, Paris, France

^12^Department of Molecular and Human Genetics, Baylor College of Medicine, Houston, TX, USA

^13^Stanley Center for Psychiatric Research, The Broad Institute of MIT and Harvard, Cambridge MA, USA

^14^Department of Biology Faculty of Natural Sciences, University of Haifa, Haifa, Israel

^15^Departments of Medicine and Genetics, Albert Einstein College of Medicine, Bronx, NY, USA

^16^Data Science Platform, Broad Institute of MIT and Harvard, Cambridge, MA, USA

^17^Department of Quantitative Health Sciences, Cleveland Clinic Research, Cleveland, OH, USA

^18^Center for Genomic Medicine, Massachusetts General Hospital, Boston, MA, USA

^19^NHLBI and Boston University’s Framingham Heart Study, Framingham, MA, USA

^20^Department of Medicine, Boston University Chobanian & Avedisian School of Medicine, Boston, MA, USA

^21^Department of Epidemiology, Boston University School of Public Health, Boston, MA, USA

^22^Department of Internal Medicine, Max Rady College of Medicine, University of Manitoba, Winnipeg, Canada

^23^Rady Faculty of Health Sciences, University of Manitoba, Winnipeg, Canada

^24^Anaesthesia and Perioperative Medicine, Division of Surgery and Interventional Science, University College London Hospitals NHS Foundation Trust, University College London, London, UK

^25^Department of Biostatistics and Center for Statistical Genetics, University of Michigan, Ann Arbor, MI, USA

^26^The Charles Bronfman Institute for Personalized Medicine, Icahn School of Medicine at Mount Sinai, New York, NY, USA

^27^Department of Biochemistry, Wake Forest School of Medicine, Winston-Salem, NC, USA

^28^British Heart Foundation Centre for Research Excellence, NIHR Leicester Biomedical Research Centre, Division of Cardiovascular Sciences, University of Leicester, Leicester, UK

^29^“Department of Neurology, Massachusetts General Hospital and Harvard Medical School, Boston, MA, USA

^30^Department of Medicine, Rutgers Robert Wood Johnson Medical School, Rutgers, The State University of New Jersey, New Brunswick, NJ, USA

^31^Department of Genetics and the Human Genetics Institute of New Jersey, School of Arts and Sciences, Rutgers, The State University of New Jersey, Piscataway, NJ, USA

^32^Meyerhoff Inflammatory Bowel Disease Center, Johns Hopkins University School of Medicine, Baltimore, MD, USA

^33^Fulcrum Genomics, Boulder, CO, USA

^34^Stanley Center for Psychiatric Research, The Broad Intitute of MIT and Harvard, Cambridge MA, USA

^35^Department of Clinical Research and Leadership, George Washington University School of Medicine and Health Sciences, Washington, DC, USA

^36^Center for Research on Genomics and Global Health, National Human Genome Research Institute, National Institutes of Health, Bethesda, MD, USA

^37^Harvard School of Public Health, Boston, MA, USA

^38^Central American Population Center, San Pedro, Costa Rica

^39^Department of Epidemiology and Biostatistics, Imperial College London, London, UK

^40^Department of Cardiology, Ealing Hospital, NHS Trust, Southall, UK

^41^Imperial College, Healthcare NHS Trust Imperial College London, London, UK

^42^Department of Medicine and Therapeutics, The Chinese University of Hong Kong, Hong Kong, China

^43^Stanley Center for Psychiatric Research, Broad Institute of MIT and Harvard, Cambridge, MA, USA

^44^Department of Medicine, Harvard Medical School, Boston, MA, USA

^45^Department of Medicine, Rutgers Robert Wood Johnson Medical School, New Brunswick, NJ, USA

^46^Northwestern University, Evanston, IL, USA

^47^Departments of Cardiovascular, Medicine Cellular and Molecular Medicine Molecular Cardiology, Quantitative Health Sciences, Cleveland Clinic, Cleveland, OH, USA

^48^Department of Pediatrics, Boston Children’s Hospital, Harvard Medical School, Boston, MAUSA

^49^McLean Hospital, Belmont, MA, USA

^50^Department of Psychiatry, Harvard Medical School, Boston, MA, USA

^51^Division of Medical Sciences, Harvard Medical School, Boston, MA, USA

^52^Genomics Platform, Broad Institute of MIT and Harvard, Cambridge, MA, USA

^53^Department of Medicine, University of Mississippi Medical Center, Jackson, MI, USA

^54^Neuropsychiatric Genetics Research Group, Dept of Psychiatry and Trinity Translational Medicine Institute, Trinity College Dublin, Dublin, Ireland

^55^Centre for Neuropsychiatric Genetics & Genomics, Cardiff University School of Medicine, Cardiff, Wales

^56^Department of Epidemiology Colorado School of Public Health Aurora, CO, USA

^57^University of Cambridge, Cambridge, England

^58^Department of Medicine and Pharmacology, University of Illinois at Chicago, Chicago, IL, USA

^59^Broad Institute of MIT and Harvard, Cambridge, MA, USA

^60^Department of Medicine, School of Medicine, University of Pittsburgh, Pittsburgh, PA, USA

^61^Department of Human Genetics, School of Public Health, University of Pittsburgh, Pittsburgh, PA, USA

^62^Clinical and Translational Science Institute, University of Pittsburgh, Pittsburgh, PA, USA

^63^Department of Life Sciences, College of Arts and Scienecs, Texas A&M University-San Antonio, San Antonio, TX, USA

^64^Department of Biostatistics, Boston University School of Public Health, Boston, MA, USA

^65^Department of Epidemiology, Biostatistics and Occupational Health, McGill University, Montreal, QC, Canada

^66^Cardiac Arrhythmia Service and Cardiovascular Research Center, Massachusetts General Hospital, Boston, MA, USA

^67^Cardiovascular Epidemiology and Genetics, Hospital del Mar Medical Research Institute (IMIM), Barcelona, Catalonia, Spain

^68^CIBER CV, Spain

^69^Departament of Medicine, Faculty of Medicine, University of Vic-Central University of Catalonia, Vic Catalonia, Spain

^70^Clalit Genomics Center, Ramat-Gan, Israel

^71^Institute for Cardiogenetics, University of Lübeck, Lübeck, Germany

^72^German Research Centre for Cardiovascular Research, Hamburg/Lübeck/Kiel, Lübeck, Germany

^73^University Heart Center Lübeck, Lübeck, Germany

^74^Estonian Genome Center, Institute of Genomics University of Tartu, Tartu, Estonia

^75^Richards Lab, Lady Davis Institute, Montreal, QC, Canada

^76^Victor Chang Cardiac Research Institute, Darlinghurst, NSW, Australia

^77^Faculty of Medicine and Health, UNSW Sydney, Kensington, NSW, Australia

^78^Cardiology Department, St Vincent’s Hospital, Darlinghurst, NSW, Australia

^79^Mayo Clinic, Arizona, USA

^80^Broad Genomics, Broad Institute of MIT and Harvard, Cambridge, MA, USA

^81^Cardiovascular Discovery Group, Kolling Institute of Medical Research, University of Sydney, Australia

^82^Department of Cardiology, Royal North Shore Hospital, Australia

^83^Faculty of Medicine and Health, University of Sydney, Australia

^84^Diabetes Unit (Department of Medicine) and Center for Genomic Medicine, Massachusetts General Hospital, Boston, MA, USA

^85^Programs in Metabolism and Medical & Population Genetics, Broad Institute of MIT and Harvard, Cambridge, MA, USA

^86^Institute of Clinical Molecular Biology, Christian-Albrechts-University of Kiel, Kiel, Germany

^87^University Hospital Schleswig-Holstein, Kiel, Germany

^88^European Molecular Biology Laboratory, European Bioinformatics Institute, Wellcome Genome Campus, Hinxton, UK

^89^Department of Neurology, Massachusetts General Hospital and Harvard Medical School, Boston, MA, USA

^90^Helsinki University and Helsinki University Hospital Clinic of Gastroenterology, Helsinki, Finland

^91^Helsinki University and Helsinki University Hospital, Helsinki, Finland

^92^Abdominal Center, Helsinki, Finland

^93^Unit of Animal Genomics, GIGA & Faculty of Veterinary Medicine, University of Liège, Liège, Belgium

^94^Bioinformatics Program MGH Cancer Center and Department of Pathology, Boston, MA, USA

^95^Cancer Genome Computational Analysis, Broad Institute of MIT and Harvard, Cambridge, MA, USA

^96^Department of Psychiatry and Behavioral Sciences, Boston Children’s Hospitaland Harvard Medical School, Boston, MA, USA

^97^Harvard Medical School Teaching Hospital, Boston, MA, USA

^98^Department of Endocrinology and Metabolism, Hadassah Medical Center and Faculty of Medicine, Hebrew University of Jerusalem, Israel

^99^Department of Psychiatry and Behavioral Sciences, SUNY Upstate Medical University, Syracuse, NY, USA

^100^Department of PSychiatry and Behavioral Sciences, Johns Hopkins University School of Medicine, Baltimore, MD, USA

^101^Centro de Investigacion en Salud Poblacional, Instituto Nacional de Salud Publica, Mexico

^102^Lund University Sweden, Sweden

^103^Institute for Molecular Medicine Finland, (FIMM) HiLIFE University of Helsinki, Helsinki, Finland

^104^Science for Life Laboratory, Department of Gene Technology, KTH Royal Institute of Technology, Stockholm, Sweden

^105^Division of Genetics and Genomics, Boston Children’s Hospital, Boston, MA, USA

^106^Center for Genetic Epidemiology, Department of Population and Public Health Sciences, University of Southern California, Los Angeles, CA, USA

^107^Human Genetics Center, University of Texas Health Science Center at Houston, Houston, TX, USA

^108^Daiichi Sankyo, Basking Ridge, NJ, USA

^109^Department of Neurology Columbia University, New York City, NY, USA

^110^Institute of Genomic Medicine, Columbia University, New York City, NY, USA

^111^Institute of Biomedicine, University of Eastern Finland, Kuopio, Finland

^112^Department of Psychiatry, Helsinki University Central Hospital Lapinlahdentie, Helsinki, Finland

^113^Department of Medical Epidemiology and Biostatistics, Karolinska Institutet, Stockholm, Sweden

^114^Icahn School of Medicine at Mount Sinai, New York, NY, USA

^115^Bonei Olam, Center for Rare Jewish Genetic Diseases, Brooklyn, NY, USA

^116^Department of Neurology, Helsinki University, Central Hospital, Helsinki, Finland

^117^Cardiovascular Disease Initiative and Program in Medical and Population Genetics, Broad Institute of MIT and Harvard, Cambridge, MA, USA

^118^Institute for Genomic Health, Icahn School of Medicine at Mount Sinai, New York, NY, USA

^119^Division of Genome Science, Department of Precision Medicine, National Institute of Health, Republic of Korea

^120^MRC Centre for Neuropsychiatric Genetics & Genomics, Cardiff University School of Medicine, Cardiff, Wales

^121^Imperial College Healthcare NHS Trust, London, UK

^122^National Heart and Lung Institute Cardiovascular Sciences, Hammersmith Campus, Imperial College London, London, UK

^123^Department of Health THL-National Institute for Health and Welfare, Helsinki, Finland

^124^Section of Cardiovascular Medicine, Department of Internal Medicine, Yale School of Medicine, New Haven, Connecticut

^125^Center for Outcomes Research and Evaluation, Yale-New Haven Hospital, New Haven, Connecticut

^126^Division of Pediatric Gastroenterology, Emory University School of Medicine, Atlanta, GA, USA

^127^Institute for Digestive Research and Department of Gastroenterology, Lithuanian University of Health Sciences, Kaunas, Lithuania

^128^Department of Internal Medicine, Seoul National University Hospital, Seoul, Republic of Korea

^129^The University of Eastern Finland, Institute of Clinical Medicine, Kuopio, Finland

^130^Kuopio University Hospital, Kuopio, Finland

^131^Department of Genetics, Yale School of Medicine, New Haven, CT, USA

^132^Department of Neuroscience and Physiology, University of Gothenburg, Gothenburg, Sweden

^133^Personalized Medicine, Mass General Brigham, Cambridge, MA, USA

^134^Department of Pathology, Brigham and Women’s Hospital, Boston, MA, USA

^135^Department of Pathology, Harvard Medical School, Boston, MA, USA

^136^Department of Clinical Chemistry Fimlab Laboratories, Tampere University, Finland

^137^Finnish Cardiovascular Research Center-Tampere Faculty of Medicine and Health Technology, Tampere University, Finland

^138^Department of Medicine, Perelman School of Medicine, University of Pennsylvania, Philadelphia, PA, USA

^139^Big Data Institute, Li Ka Shing Centre for Health Information and Discovery, University of Oxford, Oxford, UK

^140^Wellcome Trust Centre Human Genetics, University of Oxford, Oxford, UK

^141^Medical and Population Genetics Program, Broad Institute of MIT and Harvard, Cambridge,MA, USA

^142^The Novo Nordisk Foundation Center for Basic Metabolic Research, Faculty of Health and Medical Sciences, University of Copenhagen, Denmark

^143^Department of Gastroenterology, University Hospital CHU of Liège, Liège, Belgium

^144^Li Ka Shing Institute of Health Sciences, The Chinese University of Hong Kong, Hong Kong, China

^145^Hong Kong Institute of Diabetes and Obesity, The Chinese University of Hong Kong, Hong Kong, China

^146^Department of Psychiatry, Massachusetts General Hospital, Harvard Medical School, Boston, MA, USA

^147^A. A. Martinos Center, Massachusetts General Hospital, Harvard Medical School, Boston, MA, USA

^148^Division of Cardiology, University of California San Francisco, San Francisco, CA, USA

^149^Hospital del Mar Medical Research Institute (IMIM), Barcelona, Spain

^150^CIBERCV, Madrid, Spain

^151^TIMI Study Group, Division of Cardiovascular Medicine, Brigham and Women’s Hospital, Boston, MA, USA

^152^Department of Genetics, Harvard Medical School, Boston, MA, USA

^153^Oxford Centre for Diabetes, Endocrinology and Metabolism, University of Oxford, Churchill Hospital Old Road Headington, Oxford, OX, LJ, UK

^154^Welcome Centre for Human Genetics, University of Oxford, Oxford, OX, BN, UK

^155^Oxford NIHR Biomedical Research Centre, Oxford University Hospitals, NHS Foundation Trust, John Radcliffe Hospital, Oxford, OX, DU, UK

^156^John P. Hussman Institute for Human Genomics, Leonard M. Miller School of Medicine, University of Miami, Miami, FL, USA

^157^The Dr. John T. Macdonald Foundation Department of Human Genetics, Leonard M. Miller School of Medicine, University of Miami, Miami, FL, USA

^158^F. Widjaja Foundation Inflammatory Bowel and Immunobiology Research Institute Cedars-Sinai Medical Center, Los Angeles, CA, USA

^159^Atherogenomics Laboratory University of Ottawa, Heart Institute, Ottawa, Canada

^160^Division of Psychiatry, University College London, London, UK

^161^Division of General Internal Medicine, Massachusetts General Hospital, Boston, MA, USA

^162^Department of Clinical Sciences University, Hospital Malmo Clinical Research Center, Lund University, Malmö, Sweden

^163^Estonian Genome Center, Institute of Genomics, University of Tartu, Tartu, Estonia

^164^University of Arizona Health Science, Tuscon, AZ, USA

^165^University of Maryland School of Medicine, Baltimore, MD, USA

^166^The Population Health Research Institute, McMaster University and Hamilton Health Sciences, Hamilton, Ontario, Canada

^167^Division of Gastroenterology, Department of Medicine, McMaster University, Hamilton, Ontario, Canada

^168^Farncombe Family Digestive Health Research Institute, McMaster University, Hamilton, Ontario, Canada

^169^Department of Electrophysiology, Texas Cardiac Arrhythmia Institute, St. David’s MedicalCenter, Austin, Texas, USA

^170^Aging Research Center, Cinvestav Sede Sur, Center for Research and Advanced Studies of the National Polytechnic Institute, Mexico City, Mexico

^171^Computational Biology Division, Department of Integrative Biomedical Sciences, University of Cape Town, Cape Town, South Africa

^172^Institute of Infectious Disease & Molecular Medicine, Faculty of Health Sciences, University of Cape Town, Cape Town, South Africa

^173^Nutrition Research Division, International Centre for Diarrhoeal Disease Research, Bangladesh

^174^Texas Cardiac Arrhythmia Institute, St. David’s Medical Center, Austin, TX;

^175^Department of Biomedicine and Prevention, Division of Cardiology, University of Tor Vergata, Rome, Italy

^176^MetroHealth Medical Center, Case Western Reserve University School of Medicine, Cleveland, OH, USA

^177^Perelman School of Medicine, University of Pennsylvania, Philadelphia, PA, USA

^178^Johns Hopkins Bloomberg School of Public Health, Baltimore, MD, USA

^179^Kenya Medical Research Institute-Wellcome Trust Collaborative Programme, Kilifi, Kenya

^180^Dept of Psychiatry, University of Oxford, United Kingdom

^181^Lund University, Dept. Clinical Sciences, Skåne University Hospital, Malmö, Sweden

^182^Department of Genome Informatics, Graduate School of Medicine, the University of Tokyo, Tokyo, Japan

^183^Laboratory for Systems Genetics, RIKEN Center for Integrative Medical Sciences, Yokohama, Japan

^184^Premium Research Institute for Human Metaverse Medicine (WPI-PRIMe), Osaka University, Japan

^185^Center for Neurobehavioral Genetics, Semel Institute for Neuroscience and Human Behavior, University of California Los Angeles

^186^Department of Human Genetics, David Geffen School of Medicine, University of California Los Angeles, Los Angeles, CA, USA

^187^Instituto Nacional de Medicina Genómica, (INMEGEN) Mexico City, Mexico

^188^Laboratory of Immunogenomics and Metabolic Diseases, INMEGEN,Mexico City, Mexico

^189^Department of Haematology, University of Cambridge, Cambridgeshire, United Kingdom of Great Britain and Northern Ireland

^190^Applied Biomechanics Department, Swansea University, Singleton Park, Swansea SA^2 8^PP, UK

^191^Medical Research Institute, Ninewells Hospital and Medical School University of Dundee, Dundee, UK

^192^Centro de Investigación Biomédica en Red de Salud Mental (CIBERSAM), Instituto de Salud Carlos III, Madrid, Spain

^193^Department of Child and Adolescent Psychiatry, Institute of Psychiatry and Mental Health, Madrid, Spain

^194^Hospital General Universitario Gregorio Marañón, School of Medicine, Universidad Complutense, IiSGM, Madrid, Spain

^195^Department of Molecular Medicine and Biopharmaceutical Sciences, Graduate School of Convergence Science and Technology, Seoul National University, Seoul, Republic of Korea

^196^Department of Psychiatry Keck School of Medicine at the University of Southern California, Los Angeles, CA, USA

^197^Ambry Genetics, Aliso Viejo, CA, USA

^198^Department of Pediatrics/Hematology-Oncology, Baylor College of Medicine, Houston, Texas, USA

^199^Department of Complex Trait Genetics, Center for Neurogenomics and Cognitive Research, Amsterdam Neuroscience, Vrije Universiteit Amsterdam, Amsterdam, The Netherlands

^200^Department of Clinical Genetics, Amsterdam Neuroscience, Vrije Universiteit Medical Center, Amsterdam, The Netherlands

^201^Department of Psychiatry and Behavioral Sciences, Johns Hopkins University School of Medicine, Baltimore, MD, USA

^202^Departments of Human Genetics, University of Utah, Salt Lake City, UT, USA

^203^Children’s Hospital of Philadelphia, Philadelphia, PA, USA

^204^Division of Genetics and Epidemiology, Institute of Cancer Research, London, UK

^205^Department of Psychiatry, Psychosomatic Medicine and Psychotherapy; University Hospital Frankfurt - Goethe University, Frankfurt am Main, Germany

^206^University of Washington, Seattle, WA, USA

^207^Fred Hutchinson Cancer Research Center, Seattle, WA, USA

^208^Medical Research Center, Oulu University Hospital, Oulu Finland

^209^Research Unit of Clinical Neuroscience Neurology University of Oulu, Oulu, Finland

^210^Department of Genome Sciences, University of Virginia, Charlottesville, VA, USA

^211^Department of Public Health Sciences, University of Virginia, Charlottesville, VA, USA

^212^Research Center Montreal Heart Institute, Montreal, Quebec, Canada

^213^Department of Medicine, Faculty of Medicine Université de Montréal, Québec, Canada

^214^Department of Public Health Faculty of Medicine, University of Helsinki, Helsinki, Finland

^215^Section of Cardiac Electrophysiology, Division of Cardiology, Department of Medicine, Western University, London, Ontario, Canada

^216^Population Health Research Institute, Hamilton Health Sciences, and McMaster University, Hamilton, Ontario, Canada

^217^Department of Medicine, Vanderbilt, University Medical Center, Nashville, TN, USA

^218^Departments of Pharmacology and Biomedical Informatics Vanderbilt, University Medical Center, Nashville, TN, USA

^219^The Institute for Translational Genomics and Population Sciences, Department of Pediatrics, The Lundquist Institute for Biomedical Innovation at Harbor-UCLA Medical Center, Torrance, CA, USA

^220^Department of Neurology and Neurosurgery, Montreal Neurological Institute and Hospital, McGill University Health Center, Montreal, Canada

^221^Translational Sciences, Research & Development, Biogen Inc., Cambridge, MA, USA

^222^Department of Cardiovascular Sciences, University of Leicester, Leicester, UK

^223^NIHR Leicester Biomedical Research Centre, Glenfield Hospital, Leicester, UK

^224^Departments of Neuroscience, Johns Hopkins University School of Medicine, Baltimore, MD, USA

^225^Departments of Psychiatry, Johns Hopkins University School of Medicine, Baltimore, MD,USA

^226^Departments of Biomedical Engineering, Johns Hopkins University School of Medicine, Baltimore, MD, USA

^227^Department of Cardiology, Deutsches Herzzentrum München, Technical University of Munich, DZHK Munich Heart Alliance, Germany

^228^Technische Universität München, Germany

^229^Institute of Genetic Epidemiology, Department of Genetics, Medical University of Innsbruck, ^6020^ Innsbruck, Austria

^230^Faculty of Medicine, University of Southampton, Southampton, SO^16 6^YD, UK

^231^Duke Molecular Physiology Institute, Durham, NC

^232^Division of Cardiology, Department of Medicine, Duke University School of Medicine, Durham, NC, USA

^233^Division of Cardiovascular Medicine, Nashville VA Medical Center, Vanderbilt University School of Medicine, Nashville, TN, USA

^234^Division of Endocrinology, National University Hospital, Singapore

^235^NUS Saw Swee Hock School of Public Health, Singapore

^236^Channing Division of Network Medicine, Brigham and Women’s Hospital, Boston, MA, USA

^237^Harvard Medical School, Boston, MA, USA

^238^The Wallenberg Laboratory/Department of Molecular and Clinical Medicine, Institute of Medicine, Gothenburg University

^239^Department of Cardiology, Wallenberg Center for Molecular Medicine and Lund University Diabetes Center, Clinical Sciences, Lund University and Skåne University Hospital, Lund, Sweden

^240^Department of Cardiology, Sahlgrenska University Hospital, Gothenburg, Sweden

^241^Psychiatric and Neurodevelopmental Genetics Unit, Center for Genomic Medicine, Massachusetts General Hospital, Boston, MA, USA

^242^Institute of Clinical Medicine Neurology, University of Eastern Finad, Kuopio, Finland

^243^Sorbonne Université, INSERM, Centre de Recherche Saint-Antoine, CRSA, AP-HP, Saint Antoine Hospital, Gastroenterology department, F-^75012^ Paris, France

^244^INRA, UMR^1319^ Micalis, Jouy en Josas, France

^245^Paris Center for Microbiome Medicine, (PaCeMM) FHU, Paris, France

^246^Department of Twin Research and Genetic Epidemiology King’s College London, London, UK

^247^Institute of Medical Sciences, University of Aberdeen, Aberdeen, Scotland, UK

^248^Department of Medicine, Washington University School of Medicine, Saint Louis, MO, USA

^249^Department of Genetics, Washington University School of Medicine, Saint Louis, MO, USA

^250^The McDonnell Genome Institute at Washington University, Saint Louis, MO, USA

^251^Departments of Genetics and Psychiatry, University of North Carolina, Chapel Hill, NC, USA

^252^National Institute for Health and Welfare, Helsinki, Finland

^253^Saw Swee Hock School of Public Health National University of Singapore, National University Health System, Singapore

^254^Life Sciences Institute, National University of Singapore, Singapore

^255^Department of Statistics and Applied Probability, National University of Singapore, Singapore

^256^Center for Behavioral Genomics, Department of Psychiatry, University of California, San Diego, CA, USA

^257^Institute of Genomic Medicine, University of California San Diego, San Diego, CA, USA

^258^Endocrinology, Abdominal Center, Helsinki University Hospital, Helsinki, Finland

^259^Institute of Genetics, Folkhalsan Research Center, Helsinki, Finland

^260^Juliet Keidan Institute of Pediatric Gastroenterology, Eisenberg R&D Authority, Shaare Zedek Medical Center, The Hebrew University of Jerusalem, Jerusalem, Israel

^261^Instituto de Investigaciones Biomédicas, UNAM, Mexico City, Mexico

^262^Instituto Nacional de Ciencias Médicas y Nutrición Salvador Zubirán, Mexico City, Mexico

^263^Department of Public Health Faculty of Medicine University of Helsinki, Helsinki, Finland

^264^Department of Psychiatry and Human Behavior, University of California Irvine, Irvine, CA, USA

^265^Hospital Universitari Institut Pere Mata, Reus, Tarragona, Spain

^266^Institut de Recerca IRB CatSud (formerly i-CERCA), Reus, Tarragona, Spain

^267^Centro de investigación biomédica en red- CIBERSAM, Madrid, Spain

^268^Department of Statistical Genetics, Osaka University Graduate School of Medicine, Suita, Japan

^269^National Heart and Lung Institute, Imperial College London, London, UK

^270^MRC Laboratory of Medical Sciences, Imperial College London, London, UK

^271^Radcliffe Department of Medicine, University of Oxford, Oxford, UK

^272^Department of Gastroenterology and Hepatology, University of Groningen and University Medical Center Groningen, Groningen, Netherlands

^273^Big Data Institute, University of Oxford, UK

^274^Wellcome Centre for Human Genetics, University of Oxford, UK

^275^Division of Cardiology, Beth Israel Deaconess Medical Center, Boston, MA USA

^276^Program in Infectious Disease and Microbiome, Broad Institute of MIT and Harvard, Cambridge, MA, USA

^277^Center for Computational and Integrative Biology, Massachusetts General Hospital, Boston, MA, USA

^278^Stanley Division of Developmental Neurovirology, Department of Pediatrics, Johns Hopkins School of Medicine, Baltimore, MD, USA

Authors received funding as follows:

Elizabeth G. Atkinson: R01 HG012869

Emelia J. Benjamin: R01HL092577; American Heart Association AF AHA_18SFRN34110082

Matthew J. Bown: British Heart Foundation awards CS/14/2/30841 and RG/18/10/33842 Steven Brant: National Institutes of Health DK062431

Lea A. Chen: NIH, New York Crohn’s Foundation, Crohn’s and Colitis Foundation Richard H. Duerr: U01 DK062420

Ravindranath Duggirala: U01 DK085524 National Institute for Diabetes and Digestive and Kidney Diseases (NIDDK)

Josée Dupuis: National Institute for Diabetes and Digestive and Kidney Diseases (NIDDK) R DK Roberto Elosua: Agència de Gestió d’Ajuts Universitaris i de Recerca: 2021 SGR 00144 Jeanette Erdmann: VIAgenomics, Leducq network PlaqOmics, Deutsche Forschungsgemeinschaft Cluster of Excellence “Precision Medicine in Chronic Inflammation” (EXC2167);

Martti Färkkilä: State funding for university level health research

Laura D. Gauthier: Intel, Illumina

Benjamin Glaser: 5U01 DK085584

Stephen J. Glatt: U.S. NIMH Grant R MH

Leif Groop: The Academy of Finland and University of Helsinki: Center of Excellence for Complex Disease Genetics (grant number 312063 and 336822), Sigrid Jusélius Foundation; IMI 2 (grant No 115974 and 15881 )

Christopher Haiman: U01CA164973

Mikko Hiltunen: Academy of Finland (grant 338182, 353053), Sigrid Jusélius Foundation, the Strategic Neuroscience Funding of the University of Eastern Finland

Chaim Jalas: Bonei Olam

Mikko Kallela: Grants from State funding for university level health research and from Department of Neurology, Helsinki University, Central Hospital; Grant from Maire Taponen Foundation

Jaakko Kaprio: Academy of Finland (grant 352792)

Mikael Landén: Swedish Medical Research Council (2022-00642)

Terho Lehtimäki: Academy of Finland (grant 356405), VTR Pirha

Ruth J.F. Loos: Novo Nordisk Foundation (NNF18CC0034900, NNF20OC0059313); NIH (R01DK110113; R01DK124097)

Ronald C.W. Ma: Research Grants Council of the Hong Kong Special Administrative Region (CU R4012-18), Research Grants; Council Theme-based Research Scheme (T12-402/13N), University Grants Committee Research Grants Matching Scheme

Jaume Marrugat: Agència de Gestió d’Ajuts Universitaris i de Recerca: 2021 SGR 00144 Jacob L. McCauley: National Institute of Diabetes and Digestive and Kidney Disease Grant R01DK104844

Michael C. O’Donovan: Medical Research Council UK: Centre Grant No. MR/L010305/1, Program Grant No. G0800509

Yukinori Okada: JSPS KAKENHI (25H01057), and AMED (JP24km0405217, JP24ek0109594, JP24ek0410113, JP24kk0305022, JP223fa627001, JP223fa627002, JP223fa627010, JP223fa627011, JP22zf0127008, JP24tm0524002, JP24wm0625504, JP24gm1810011), JST

Moonshot R&D (JPMJMS2021, JPMJMS2024), Takeda Science Foundation, Ono Pharmaceutical Foundation for Oncology, Immunology, and Neurology, Bioinformatics Initiative of Osaka University Graduate School of Medicine, Institute for Open and Transdisciplinary Research Initiatives, Center for Infectious Disease Education and Research (CiDER), and Center for Advanced Modality and DDS (CAMaD), Osaka University, RIKEN TRIP initiative (AGIS)

Michael J. Owen: Medical Research Council UK: Centre Grant No. MR/L010305/1, Program Grant No. G0800509

Aarno Palotie: the Academy of Finland Center of Excellence for Complex Disease Genetics (grant numbers 312074 and 336824) and Sigrid Jusélius Foundation

John D. Rioux: National Institute of Diabetes and Digestive and Kidney Diseases (NIDDK; DK062432), from the Canadian Institutes of Health (CIHR GPG 102170), from Genome Canada/Génome Québec (GPH-129341), and a Canada Research Chair (#230625)

Samuli Ripatti: the Academy of Finland Center of Excellence for Complex Disease Genetics (grant number ) Sigrid Jusélius Foundation

Jerome I. Rotter: Trans-Omics in Precision Medicine (TOPMed) program was supported by the National Heart, Lung and Blood Institute (NHLBI). WGS for “NHLBI TOPMed: Multi-Ethnic Study of Atherosclerosis (MESA)” (phs001416.v1.p1) was performed at the Broad Institute of MIT and Harvard (3U54HG003067-13S1). Core support including centralized genomic read mapping and genotype calling, along with variant quality metrics and filtering were provided by the TOPMed Informatics Research Center (3R01HL-117626-02S1; contract HHSN268201800002I). Core support including phenotype harmonization, data management, sample-identity QC, and general program coordination were provided by the TOPMed Data Coordinating Center (R01HL-120393; U01HL-120393; contract HHSN268201800001I). We gratefully acknowledge the studies and participants who provided biological samples and data for MESA and TOPMed. JSK was supported by the Pulmonary Fibrosis Foundation Scholars Award and grant K23-HL-150301 from the NHLBI. MRA was supported by grant K23-HL-150280, AJP was supported by grant K23-HL-140199, and AM was supported by R01-HL131565 from the NHLBI. EJB was supported by grant K23-AR-075112 from the National Institute of Arthritis and Musculoskeletal and Skin Diseases.The MESA project is conducted and supported by the National Heart, Lung, and Blood Institute (NHLBI) in collaboration with MESA investigators. Support for MESA is provided by contracts 75N92020D00001, HHSN268201500003I, N01-HC-95159, 75N92020D00005, N01-HC-95160, 75N92020D00002, N01-HC-95161, 75N92020D00003, N01-HC-95162, 75N92020D00006, N01-HC-95163, 75N92020D00004, N01-HC-95164, 75N92020D00007, N01-HC-95165, N01-HC-95166, N01-HC-95167, N01-HC-95168, N01-HC-95169, UL1-TR-000040, UL1-TR-001079, and UL1-TR-001420. Also supported in part by the National Center for Advancing Translational Sciences, CTSI grant UL1TR001881, and the National Institute of Diabetes and Digestive and Kidney Disease Diabetes Research Center (DRC) grant DK063491 to the Southern California Diabetes Endocrinology Research Center Kaitlin E. Samocha: R01HG012867

Jeremiah Scharf: NIH Grants U01 NS40024, K02 NS085048, NS102371

Eleanor G. Seaby: Kerkut Charitable Trust, Foulkes Fellowship, University of Southampton Presidential Scholarship

Edwin K. Silverman: NIH Grants U01 HL089856 and U01 HL089897

J. Gustav Smith: The Swedish Heart-Lung Foundation (2022-0344, 2022-0345), the Swedish Research Council (2021-02273), the European Research Council (ERC-STG-2015-679242), Gothenburg University, Skåne University Hospital, governmental funding of clinical research within the Swedish National Health Service, a generous donation from the Knut and Alice Wallenberg foundation to the Wallenberg Center for Molecular Medicine in Lund, and funding from the Swedish Research Council (Linnaeus grant Dnr 349-2006-237, Strategic Research Area Exodiab Dnr 2009-1039) and Swedish Foundation for Strategic Research (Dnr IRC15-0067) to the Lund University Diabetes Center

Harry Sokol: AgroParisTech, Jouy en Josas, France

Nathan O. Stitziel: National Human Genome Reserach Institute Grant UM1HG008853

Kent D. Taylor: Trans-Omics in Precision Medicine (TOPMed) program was supported by the National Heart, Lung and Blood Institute (NHLBI). WGS for “NHLBI TOPMed: Multi-Ethnic Study of Atherosclerosis (MESA)” (phs001416.v1.p1) was performed at the Broad Institute of MIT and Harvard (3U54HG003067-13S1). Core support including centralized genomic read mapping and genotype calling, along with variant quality metrics and filtering were provided by the TOPMed

Informatics Research Center (3R01HL-117626-02S1; contract HHSN268201800002I). Core support including phenotype harmonization, data management, sample-identity QC, and general program coordination were provided by the TOPMed Data Coordinating Center (R01HL-120393; U01HL-120393; contract HHSN268201800001I). We gratefully acknowledge the studies and participants who provided biological samples and data for MESA and TOPMed. JSK was supported by the Pulmonary Fibrosis Foundation Scholars Award and grant K23-HL-150301 from the NHLBI. MRA was supported by grant K23-HL-150280, AJP was supported by grant K23-HL-140199, and AM was supported by R01-HL131565 from the NHLBI. EJB was supported by grant K23-AR-075112 from the National Institute of Arthritis and Musculoskeletal and Skin Diseases.The MESA project is conducted and supported by the National Heart, Lung, and Blood Institute (NHLBI) in collaboration with MESA investigators. Support for MESA is provided by contracts 75N92020D00001, HHSN268201500003I, N01-HC-95159, 75N92020D00005, N01-HC-95160, 75N92020D00002, N01-HC-95161, 75N92020D00003, N01-HC-95162, 75N92020D00006, N01-HC-95163, 75N92020D00004, N01-HC-95164, 75N92020D00007, N01-HC-95165, N01-HC-95166, N01-HC-95167, N01-HC-95168, N01-HC-95169, UL1-TR-000040, UL1-TR-001079, and UL1-TR-001420. Also supported in part by the National Center for Advancing Translational Sciences, CTSI grant UL1TR001881, and the National Institute of Diabetes and Digestive and Kidney Disease Diabetes Research Center (DRC) grant DK063491 to the Southern California Diabetes Endocrinology Research Center Tiinamaija Tuomi: The Academy of Finland and University of Helsinki: Center of Excellence for Complex Disease Genetics (grant number 312072 and 336826 ), Folkhalsan Research Foundation, Helsinki University Hospital, Ollqvist Foundation, Liv och Halsa foundation; NovoNordisk Foundation

Teresa Tusie-Luna: CONACyT Project 312688

James S. Ware: JSW is supported by Medical Research Council (UK), Sir Jules Thorn Charitable Trust [21JTA], British Heart Foundation [RE/24/130023], and the NIHR Imperial Biomedical Research Centre

Rinse K. Weersma: The Lifelines Biobank initiative has been made possible by subsidy from the Dutch Ministry of Health Welfare and Sport the Dutch Ministry of Economic Affairs the University Medical Centre Groningen (UMCG the Netherlands ) the University of Groningen and the Northern Provinces of the Netherlands

Conflicts of interest are as follows:

Mikko Kallela: No related COI

Eimear E. Kenny: EEK has received personal fees from Regeneron Pharmaceuticals, 23&Me, Allelica, and Illumina; has received research funding from Allelica; and serves on the advisory boards for Encompass Biosciences, Foresite Labs, and Galateo Bio

Mikael Landén: M.L. has received lecture honoraria from Lundbeck pharmaceutical.

Ruth J.F. Loos: R.J.F.L has received consultancy and speaker fees from Novo Nordisk and Eli Lilly and Company

Ronald C.W. Ma: No related COI

Edwin K. Silverman: Research grants from GSK and Bayer

James S. Ware: JSW has received research support from Bristol Myers Squibb, has acted as a paid advisor to MyoKardia, Pfizer, Foresite Labs, Health Lumen, Tenaya Therapeutics, and Solid Biosciences, and is a founder with equity in Saturnus Bio.

