## Supplementary information for "The landscape of regional missense mutational intolerance quantified from 730,947 exomes"

### Supplementary Note

### Identification of regions with distinct missense constraint strengths within genes

Broadly, we were interested in identifying variability in missense constraint (as measured by the depletion of missense variation compared to a null mutational model) within protein-coding transcripts. Transcripts where we did not find evidence of regional variability in missense constraint are deemed to have one missense constraint region (MCR) from the start to the end of the coding sequence. As reported in the main text, we found that 36% of 17,841 MANE Select or canonical transcripts have ≥2 MCRs.

#### Determining minimum missense constraint region (MCR) size

We ran a power analysis to determine the minimum MCR size that our search should emit, based on the simplest single break scenario, i.e. a single breakpoint dividing a transcript into two MCRs, where one MCR had a missense observed/expected ratio (OE) of 0, and the other MCR had an OE of 1, as this is the maximum difference in OE values for our break search. We calculated a p-value for exome-wide significance of $2.8 \times 10^{-6}$ (0.05 / 17,841 transcripts), determining that the minimum number of expected variants on either side of a breakpoint position needed to meet this significance threshold was 16 (**Supplementary Fig. 1**).

#### MCRs and gene length

Gene coding length and number of MCRs are positively correlated (Spearman $\rho = 0.41$, $p < 10^{-18}$; **Supplementary Fig. 2**). This occurs for two reasons: 1. larger mutational space and thus greater statistical power in calling MCRs, especially when constraint differences are modest; and 2. greater probability of distinct functional components within genes. We thus control for gene coding length by regression when comparing MCR count across gene sets.

#### MCR size

In transcripts with ≥2 MCRs, the minimum MCR coding length is 20bp, and the median is 339 bp (**Supplementary Fig. 3a**). In these transcripts, we observe that regions estimated to be severely missense-depleted tend to be smaller: MCRs with OE < 0.4 have a median coding length of 83.5bp, while MCRs with OE ≥ 0.4 have a median coding length of 464bp (Wilcoxon $p < 10^{-50}$; **Supplementary Fig. 3b-c**). This difference is not driven by small low-coverage MCRs: high-coverage MCRs (%AN ≥ 90) with OE < 0.4 have a median coding length of 80bp and high-coverage MCRs with OE ≥ 0.4 have a median coding length of 474 bp.

#### Sequencing coverage and quantification of missense depletion

As detailed in the Methods, we used allele number percent (%AN) in the gnomAD v4.1.1[1] exomes to proxy coverage for all analyses to improve our ability to capture constraint in lower coverage sites. Prior to searching for MCR breakpoints within transcripts, we removed sites with %AN < 20 (0.68% of all possible missense variant sites) and adjusted expected variant counts for %AN as described elsewhere[1]. We evaluated post-hoc if there was any residual relationship between coverage and estimated MCR constraint. In transcripts with ≥2 MCRs (i.e. where breakpoints were identified), we binned MCRs into three bins based on their median %AN: 20-60, 60-90 and ≥90 (**Supplementary Fig. 4**) and found that low-coverage MCRs tend to be enriched for stronger constraint estimates. For example, a larger proportion of MCRs are highly missense-depleted (MCR missense OE < 0.4) in the 20-60 %AN bin (124/242; 51%) as compared to the ≥90 %AN bin (5,489/24,841; 22%; Fisher's exact $p = 9.7 \times 10^{-13}$). We thus caution against overinterpretation of highly constrained MCRs within low coverage sequences, though MCRs with median %AN < 90 and missense OE < 0.4 make up only a small (<1.5%) portion of the data. All MCRs regardless of coverage are made available in the data release and those with %AN < 90 have been flagged.

#### Transcripts with outlier variant counts

In addition to the 17,841 high-quality QC PASS MANE Select or canonical transcripts, we also performed MCR breakpoint identification on 1,534 MANE Select or canonical transcripts with outlier variant counts. Outlier transcripts were defined as those with zero expected or too many observed pLoF, missense, or synonymous variants; or too few observed synonymous variants as described in gnomAD v4.1.1[1]. 378 of the 1,534 (25%) transcripts were found to have ≥2 MCRs. MCR information in these transcripts is available in the data release but with an outlier transcript flag. We caution that these constraint estimates are likely less well-calibrated, for example being biased towards higher or lower OE.

#### Unconstrained MCRs and RNA expression

While we analyze the MANE Select or Ensembl canonical transcript for each gene in this study, it is possible that a given transcript may not be the most biologically appropriate or disease-appropriate gene model. We thus assessed whether unconstrained MCRs could actually represent exons that are excluded from the most relevant gene transcripts as an explanation for their lack of constraint. To do so, we evaluated the RNA expression output, quantified using the site-level proportion expressed across transcripts (pext) metric, across MCRs[1,2]. For transcripts with ≥2 MCRs, we calculated the median pext score across each MCR and compared the constraint distribution for MCRs with low median pext (<0.1) vs. MCRs with high median pext (≥0.9). We found slightly more MCRs with low median pext scores (377/1,062 or 36%) are unconstrained (MCR missense OE > 0.9) compared to MCRs with high median pext scores (3,108/9,826 or 32%), but the difference is not significant (Fisher's exact $p = 0.070$).

#### Comparing missense constraint to other measures of constraint, conservation, and disease association

##### Comparing missense constraint to pLoF constraint

We observed that transcripts that are more intolerant to pLoF variation, as measured by LOEUF, tend to be more intolerant to missense variation. This pattern is evident when quantifying missense intolerance with transcript missense OE, but grows stronger when quantifying with missense OE in the most constrained MCR in a transcript (Spearman ρ = 0.51 vs. 0.58; **Supplementary Fig. 5**). For example, we found that 12% (646) of 5,270 pLoF-constrained genes (within the first three LOEUF deciles) are missense-constrained (missense OE < 0.6) across their entire transcript. An additional 49% (2,573) genes are missense-constrained at the same threshold when assessing their minimum MCR OE. In contrast, in the three most pLoF-unconstrained deciles (last three deciles of LOEUF), only 2% (93/4,701) of genes are missense-constrained at the transcript level, and only an additional 8% (369) are MCR missense-constrained.

To assess the added value of missense constraint in identifying disease genes beyond pLoF constraint, we looked at genes that had definitive or strong evidence of monoallelic association with a DD in G2P but which were also LOEUF-unconstrained (highest three LOEUF deciles; n=34). In these genes, the LOEUF metric does not suggest a strong fitness effect for pLoF variants even though there is clear evidence for association with a severe disease. We found that 38% (13) of these genes have a constrained MCR with OE < 0.6. This represents a nearly 6-fold enrichment for having a constrained MCR compared to genes in the same LOEUF deciles without OMIM or G2P disease associations (385/3,929; OR = 5.7, Fisher's exact $p = 1.1 \times 10^{-5}$). This demonstrates that missense constraint can potentially identify genes that are causally linked to severe diseases that LOEUF does not capture well. This is particularly pertinent for genes where LOEUF is less well-powered (11/13 or 85% of these genes have <10 expected LoF variants) or that act through non-LoF mechanisms (4/13 or 31% of these genes are DD-associated in G2P through a non-LoF mechanism only). When performing this same comparison for missense constraint at the transcript-level rather than MCR-level, we find that the enrichment for missense depletion in the DDG2P vs. non-disease associated genes is stronger but captures fewer overall disease genes (5/34 monoallelic DDG2P genes, 77/3,929 non-DDG2P and non-OMIM genes, or OR = 8.6, Fisher's exact $p = 5.8 \times 10^{-4}$).

#### Comparing missense constraint to OMIM disease association

Compared to autosomal genes with dominant disease association (n=1,036), autosomal genes not associated with disease in OMIM (n=12,623) tended to be less missense-constrained with higher transcript-wide missense OE (median transcript-wide missense OE = 0.92 vs 0.81, Welch's t $p < 10^{-50}$) as well as minimum MCR missense OE (median minimum MCR missense OE = 0.89 vs. 0.43, Wilcoxon $p < 10^{-50}$). However, compared to autosomal genes with recessive disease association (n=2,176), these non-OMIM genes appear to be generally more missense-constrained with lower transcript-wide missense OE (median transcript-wide missense OE = 0.92 vs. 0.93, Welch's t $p = 1.1 \times 10^{-16}$) as well as minimum MCR missense OE (median minimum MCR missense OE = 0.89 vs. 0.91, Wilcoxon $p = 9.0 \times 10^{-7}$). This relationship persists when subsetting to genes with ≥2 MCRs: across genes not in OMIM (n= 4,083), we again observe that transcript-wide missense OE tends to be higher than for autosomal dominant genes (n=665; median = 0.82 vs. 0.75, Welch's t $p = 5.0 \times 10^{-22}$) but lower than for autosomal recessive genes (n=594; median = 0.82 vs. 0.87, Welch's t $p = 5.0 \times 10^{-21}$) and likewise for minimum MCR missense OE (median = 0.38 vs. 0.24, Wilcoxon $p = 7.8 \times 10^{-36}$ for non-OMIM vs. dominant; median = 0.38 vs. 0.43, Wilcoxon $p = 4.2 \times 10^{-5}$ for non-OMIM vs. recessive)

(**Supplementary Fig. 6**). This suggests that a sizable subset of as-of-yet disease-unassociated genes are under substantial heterozygous selection against missense variants. Accordingly, when incorporating pLoF constraint, we find that 62% (1,511/2,430) of genes that are both pLoF-constrained and severely missense-depleted (first three deciles of LOEUF and MCR with OE < 0.4) do not have disease associations in OMIM (**Supplementary Fig. 7**).

#### Comparing missense constraint to conservation

We investigated potential sites of divergence between recent selection on human variation (measured by missense constraint) compared to selection over longer timescales (measured by evolutionary conservation in placental mammals, phyloP[3]). Comparing phyloP to missense constraint quantified transcript-wide, we found that transcripts with more conserved coding sequences tended to also be more depleted of human missense variation (Spearman $\rho = 0.50$, $p < 10^{-50}$), consistent with previous correlations found between stronger mammalian conservation and lower human allele frequency[3] (**Supplementary Fig. 8a, c**). However, when comparing phyloP to MCR-wide missense constraint, we discover a substantial number of MCRs that appear strongly constrained against missense variants in humans but widely unconserved across mammals (Spearman $\rho = 0.44$, $p < 10^{-50}$), potentially pointing to human-specific negative selection pressures at specific sites within genes that are obscured when smoothing constraint over whole transcripts (**Supplementary Fig. 8b, d**). There is no significant difference in location of genes with these MCRs on autosomes vs. allosomes (Fisher's exact p = 0.16).

We found 274 autosomal genes have at least one severely depleted MCR (missense OE ≤ 0.4) where ≥70% of coding bases are unconserved[3] across placental mammals (phyloP < 2.7). A Gene Ontology (GO) enrichment analysis of these genes against a background set of all 4,896 autosomal genes with ≥2 MCRs using shinyGO v0.85[4] revealed enrichments concentrated in immune-related pathways (**Supplementary Tables 3-4**). We do not see a similar accumulation of mammalian-conserved yet human-unconstrained regions, indicating that negative selection pressures that have been maintained across mammalian evolution are unlikely to be waived in humans.

| Enrichment FDR | Fold Enrichment | N Genes | Total N Pathway Genes | Pathways |
|---|---|---|---|---|
| 0.01 | 22.5 | 3 | 49 | MHC class I receptor activity |
| 0.01 | 22.5 | 3 | 41 | Inhibitory MHC class I receptor activity |
| 0.0088 | 18 | 4 | 21 | Sialic acid binding |
| 0.01 | 6.6 | 7 | 276 | Immune receptor activity |

**Supplementary Table 3**
GO enrichment for molecular function of generally unconserved autosomal genes (≥70% of coding bases have phyloP < 2.7) with a severely depleted MCR (MCR missense OE < 0.4).

| Enrichment FDR | Fold Enrichment | N Genes | Total N Pathway Genes | Pathway |
|---|---|---|---|---|
| 0.031 | 11.3 | 4 | 48 | Antigen processing and presentation of peptide antigen via MHC class Ib |
| 0.025 | 9.4 | 5 | 98 | T cell mediated cytotoxicity |
| 0.025 | 9.4 | 5 | 90 | Reg. of t cell mediated cytotoxicity |
| 0.025 | 7.1 | 6 | 106 | Pos. reg. of leukocyte mediated cytotoxicity |
| 0.018 | 6.9 | 7 | 166 | Reg. of leukocyte mediated cytotoxicity |
| 0.018 | 6.4 | 8 | 182 | Reg. of cell killing |
| 0.031 | 6.4 | 6 | 115 | Pos. reg. of cell killing |
| 0.025 | 6.1 | 7 | 180 | Antigen processing and presentation of peptide antigen |
| 0.018 | 5.2 | 9 | 228 | Leukocyte mediated cytotoxicity |
| 0.0078 | 5 | 12 | 332 | Cell killing |
| 0.024 | 4.4 | 10 | 196 | Pos. reg. of lymphocyte mediated immunity |
| 0.018 | 4.1 | 12 | 272 | Reg. of lymphocyte mediated immunity |
| 0.025 | 4.1 | 10 | 348 | Humoral immune response |
| 0.018 | 3.5 | 14 | 334 | Reg. of leukocyte mediated immunity |
| 0.027 | 3.1 | 14 | 424 | Lymphocyte mediated immunity |
| 0.025 | 2.8 | 17 | 501 | Reg. of immune effector proc. |

**Supplementary Table 4**
GO enrichment for biological processes of generally unconserved autosomal genes (≥70% of coding bases have phyloP < 2.7) with a severely depleted MCR (MCR missense OE < 0.4)

### Selection of missense OE thresholds in enrichment tests

We chose the thresholds used to define missense-constrained regions referred to throughout this study based on a number of lines of evidence. First, we compared the rate of *de novo* missense variation in individuals with developmental disorders (DD; n=31,058)[5] to the rate of *de novo* missense variation in sibling controls (n=5,492)[6] across five different bins of missense OE, filtered to high-coverage sequences (**Supplementary Fig. 9, Supplementary Tables 5-8**). Given that only a subset of the *de novo* missense variants in DD individuals are causal for disease, we expected that regions intolerant of missense variation would have higher rate ratios (RR), and regions tolerant of missense variation would have a RR close to 1 (equal rates of *de novo* missense variation in cases and controls). We observed that the bins with missense OE < 0.6 had RRs substantially exceeding 1 compared to the bins with missense OE ≥ 0.6. Second, after applying a previously established probabilistic framework estimating thresholds on variant scores[7] (see *Clinical calibration of regional constraint metrics*), we found that missense OE ≤ 0.59 met supporting and missense OE ≤ 0.36 met moderate level evidence for pathogenicity

(**Supplementary Table 13**). We thus chose a threshold of OE < 0.6 to define “constrained” MCRs and a threshold of OE < 0.4 to define “severely depleted” MCRs. These correspond to the 9th and 4th percentiles of constraint, respectively.

### Partitioning heritability enrichment over missense constraint

We estimated the partitioned heritability enrichment in five SNP categories, comprising coding SNPs in each missense constraint quintile. Quintiles were computed over OE at sites of potential missense variants in the high-coverage MCRs of the 17,841 high-quality transcripts. These quintile bins on missense OE were: 0-0.78, 0.78-0.89, 0.89-0.96, 0.96-1.02, and 1.02+. Heritability enrichment was computed for each of 268 traits and each of these five bins as previously described[8]. Briefly, for each trait, per-SNP heritability was calculated using LD score regression, and heritability enrichment was calculated for each category of SNPs as the proportion of heritability ascribed to SNPs in the category divided by the proportion of SNPs in that category (**Supplementary Fig. 10, Supplementary Tables 9-10**). SNPs were filtered to those in high-coverage MCRs. The traits analyzed were those described in previous work[9] after removing traits with a maximum pairwise correlation greater than 0.2 for a total of 268 independent traits analyzed in this study. Trait name encodings are also as previously described[9]. The mean heritability enrichment for all coding SNPs in MCRs was 4.5 with a standard deviation of 9.7.

### Clinical calibration of regional constraint metrics

#### Datasets

We filtered to retain genes that had at least one pathogenic missense variant in ClinVar (Accessed November 9, 2025) and then selected missense variants in those genes that had at least a one-star review status and one of the following classifications: pathogenic, pathogenic/likely pathogenic, likely pathogenic, benign, benign/likely benign, or likely benign. Variants with an allele frequency (AF) ≥ 1% in gnomAD v4.1.1 or with conflicting classifications were removed. This process yielded a total of 122,269 ClinVar variants from 4,334 genes. We also retrieved missense variants from gnomAD v4.1.1 exomes in the genes where we collected ClinVar variants as described above. We retained only QC-pass variants ("PASS" in the "FILTER" column") and removed variants with an AF ≥ 1%. This resulted in a set of 3,847,501 gnomAD variants from 4,302 genes.

#### Calibration

We calibrated MCR missense OE and the missense tolerance ratio (MTR)[10] metric to evaluate their utility in clinical variant classification. We first annotated the ClinVar and gnomAD variants from the datasets described above with scores from each metric. Using the previously developed method[7] and the prior probability of pathogenicity of 4.41% established therein, we computed the local posterior probability curves for all score values given by each method. ClinVar variants were used with their classification labels to estimate the local values of the posterior probability of pathogenicity, and gnomAD variants were used to smooth out the posterior estimates (≥1% gnomAD variants required in each local window). Consequently, we defined score threshold ranges for each strength of evidence (supporting, moderate, strong, and very strong) for pathogenicity and benignity based on the ACMG/AMP criteria. For each metric, we plotted the local posterior probability curve estimate along with the one-sided 95% confidence interval, calculated on the more stringent side, determined using 10,000 bootstrapping iterations (**Fig. 4b**). Calibration found that MCR missense OE ≥ 0.9704 met moderate and OE ≥ 1.2254 met supporting evidence for BP1 (**Supplementary Fig. 12**). The final thresholds for each tool as determined with bootstrapping are provided in **Supplementary Table 13**.

| | Benign (BP1) | | | | Pathogenic (PP2) | | | |
|---|---|---|---|---|---|---|---|---|
| **Method** | Very Strong | Strong | Moderate | Supporting | Supporting | Moderate | Strong | Very Strong |
| MCR OE | - | - | 1.2254 | 0.9704 | 0.5936 | 0.3624 | - | - |
| MTR | - | - | - | - | 0.7601 | 0.5685 | - | - |

**Supplementary Table 13**
Thresholds on MCR missense OE and MTR meeting each tier of evidence for missense variant pathogenicity or benignity, calibrated based on the ACMG/AMP variant classification guidelines.

### Development of a constraint-informed variant-level missense deleteriousness predictor (MPC)

#### Training and test datasets

To generate independent training and test sets, we selected 80% (14,263 transcripts) of the 17,841 MANE Select or canonical coding transcripts to comprise the overall possible training transcript set and the remaining 20% (3,578 transcripts) the overall possible test transcript set. To ensure these had similar distributions of features that may impact constraint estimates, we used stratified randomization to match these transcript sets on $s_{het}$[10] coefficients (as a measure of selection) and number of potential missense sites (as a measure of power to detect transcript-wide constraint changes). The training set was used for MPC model training including hyperparameter tuning with cross-validation, and the test set was used for evaluation of MPC model architectures. From these transcript sets, we further extracted transcripts that were likely to contain severely deleterious missense variants as those corresponding to 2,987 predicted-haploinsufficient genes (probability of haploinsufficiency or pHaplo ≥ 0.86[11]) or to 359 genes with DD associations in G2P through non-LoF mechanisms (total 3,161 genes). For the classification task, the “pathogenic” variant set consisted of high-quality ClinVar P/LP variants in these genes. The “benign” variant set consisted of high-quality ClinVar B/LB or common (AF > 0.1%) variants in these genes from gnomAD. Variants matching criteria for both the “benign” and “pathogenic” sets were excluded. We further removed variants with %AN < 20 in the gnomAD v4.1.1 exomes and variants that were not in the 17,841 MANE Select/canonical transcripts. Finally, we only included variants with all 17 features evaluated in model training (**Supplementary Table 15**). In total, we extracted 20,931 “pathogenic” variants and 93,638 “benign” variants, of which 17,373 “pathogenic” variants and 83,330 “benign” variants were in the training set.

#### Variant classification model architecture and feature selection

In training the model to classify “pathogenic” vs. “benign” variants, we evaluated 17 features: two at the variant-level, five at the amino acid substitution class-level, six at the MCR-level, and four at the gene-level (**Supplementary Table 15**). “Upstream” and “downstream” MCRs refer to the MCRs immediately upstream or downstream, respectively, of the MCR housing the given variant. If not applicable (i.e. only one MCR in a transcript, or the first/last MCR in a transcript), this was instead assigned to be the MCR housing the given variant.

The “amino acid substitution class OE”, “amino acid substitution class OE second derivative”, and “missense badness” features are calculated using our MCR missense OE metric to quantify the increased deleteriousness of amino acid substitution classes (e.g., Met to Tyr) in functionally important areas of proteins (**Supplementary Table 14**). To calculate “amino acid substitution class OE”, we divided the total number of rare, high quality single-nucleotide variants observed in gnomAD (see **Methods**) resulting in the given amino acid substitution class by the total number of expected variants resulting in that substitution class. To calculate “amino acid substitution class OE second derivative”, we aggregated the OEs of each substitution class by MCR OE bin in 10 bins from 0 to 1.0+ (i.e., for the 0-0.1 OE bin, we calculated all of the observed substitutions that occurred within regions with a OE between 0 and 0.1 and divided that number by the total number of expected substitutions occurring in those regions). We then took the second derivative of the regression line, observing that this correlated with commonly used metrics of amino acid deleteriousness like BLOSUM (**Supplementary Fig. 13**). Missense badness was calculated similarly as previously described[12]. Briefly, for each amino acid substitution class, we summed all possible high-quality single-nucleotide variants that could result in that substitution class and the subset that were observed at rare frequency in gnomAD (see **Methods**). We split these counts by whether variants were in the most constrained MCR missense OE decile (OE ≤ 0.64) or the remaining nine MCR missense OE deciles (OE > 0.64), calculated across high-coverage MCRs. Then, for each substitution class, we calculated the fold-difference between observed divided by possible counts in the most constrained MCR missense OE decile vs. the remaining deciles. These were then normalized with the fold-difference for all synonymous substitutions as a floor (set to 0) and the fold-difference for all nonsense substitutions as a ceiling (set to 1), producing “missense badness” values between 0 and 1.

We evaluated two model architectures: logistic regression with L1 regularization and XGBoost (gradient-boosted tree) with recursive feature elimination and cross-validation. Simple random forest was unsuitable for our purposes, as the number of trees limits the number of emitted probabilities upon which the MPC score calculation is based. The XGBoost model was chosen over logistic regression based on performance on the test set and then retrained on all data for the final model, whose features are given in **Supplementary Table 15**.

| Feature Type | Feature Name | Selected for MPC |
|---|---|---|
| Variant | PolyPhen-2 | Yes |
| Variant | phyloP | Yes |
| Amino acid substitution class | BLOSUM | No |
| Amino acid substitution class | Grantham | No |
| Amino acid substitution class | Amino acid substitution class OE | No |
| Amino acid substitution class | Amino acid substitution class OE second derivative | Yes |
| Amino acid substitution class | Missense badness | Yes |
| MCR | MCR missense OE | Yes |
| MCR | MCR missense expected | Yes |
| MCR | Upstream MCR missense OE | No |
| MCR | Upstream MCR missense expected | Yes |
| MCR | Downstream MCR missense OE | Yes |
| MCR | Downstream MCR missense expected | No |
| Gene | Gene missense OE | Yes |
| Gene | Gene missense expected | Yes |
| Gene | Gene pLoF OE | Yes |
| Gene | Gene pLoF expected | Yes |

**Supplementary Table 15**
Model features evaluated in classification of “pathogenic” vs. “benign” variants and whether they were selected for the final XGBoost classification model.

#### Conversion of variant classification prediction probabilities to MPC scores

We applied the final XGBoost model across the 70,313,598 possible exome-wide missense variants in the Ensembl VEP table with all features to obtain a fitted probability of pathogenicity value for each missense variant *i*. The MPC score for missense variant *i* is given as:

$$d_i = -log_{10}(m_i/M)$$

where $d_i$ is the MPC score, $m_i$ is the number of “benign” missense variants in the training set with a fitted probability of pathogenicity value that is less than the fitted value for variant *i*, and *M* is the total number of “benign” missense variants in the training set. When $m_i$ is 0, i.e. when the fitted score for variant *i* is more severe than for all “benign” variants, this introduces a log-zero error so we set $d_i$ to 6 (the maximum real value for $d_i$ is just over 5 when $m_i$ is 1). Larger values of $d_i$ indicate stronger predicted-deleteriousness (**Supplementary Fig. 14**).

We note that the MPC scores in our data release are transcript-specific. This means that if a base pair substitution causes a missense change in the MANE Select or canonical transcript for two genes, it will be listed twice, with one score corresponding to each transcript. This is because the model uses gene-/transcript-specific annotations in variant deleteriousness prediction. For annotation of the rare and *de novo* variants in the developmental disorder cohorts analyzed, we assigned a single value to each unique variant by taking the maximum (i.e., most deleterious) transcript-specific MPC score.

### Evaluating the performance of MPC

As a sanity check, we observed that as gnomAD AF of a variant increases from 0% (not observed in gnomAD) to >5%, MPC scores decrease, predicting lower variant deleteriousness (**Supplementary Fig. 15**). Additionally, the MPC scores for *de novo* missense variants in individuals with DD tend to be higher than autistic individuals (Wilcoxon $p < 10^{-45}$), which in turn tend to be higher than unaffected siblings (Wilcoxon $p = 5.2 \times 10^{-6}$; **Supplementary Fig. 16**).

#### Association testing in neurodevelopmental disorders (NDDs) with MPC

To further pinpoint enrichment across specific regimes within the spectrum of predicted-deleteriousness, we compared case and control missense rates stratified by MPC bins (**Supplementary Fig. 17**). We additionally stratified by localization of variants to 373 known NDD-associated genes[6] in order to probe the genetic architecture of NDDs that remains to be mapped. We created three bins of MPC scores: <2, 2-2.5, ≥2.5. The MPC thresholds defining each of these bins were selected as the thresholds at which the NDD case-control *de novo* rate ratio of missense variants met the rate ratio of protein-truncating variants in pLoF-constrained genes. To determine these thresholds, we calculated the DD and ASD *de novo* rate ratio of pLoF variants in genes in the first two LOEUF deciles and averaged them together (DD RR = 5.6, ASD RR = 2.6, averaged RR = 4.1). Then, for each sliding MPC threshold starting from 0 and incrementing by 0.1, we averaged together the DD and ASD *de novo* rate ratios of missense variants with MPC greater than or equal to that threshold. The first MPC threshold at which the averaged missense rate ratio met or exceeded that of the pLoF variants at 4.1 was MPC ≥ 2.5 (96th percentile of MPC). We repeated this for the next most damaging tier of variants, comparing the averaged DD and ASD rate ratio for pLoF variants in the third and fourth LOEUF deciles (DD RR = 1.8, ASD RR = 1.4, averaged RR = 1.6) to those for missense variants with MPC greater than or equal to each sliding threshold but less than 2.5. This resulted in the selection of the second MPC threshold of 2 (92nd percentile of MPC). In the joint evaluation of MPC and AlphaMissense in **Supplementary Fig. 19**, “MPC-high” variants are defined with the most stringent cutoff of MPC ≥ 2.5, and “MPC-low” variants are defined as MPC < 2. The same methods were used to define the “AlphaMissense-high” and “AlphaMissense-low” score thresholds of 0.9985 and 0.761 (incrementing by 0.0005).

The confidence interval of the relative difference statistics is calculated based on a binomial test, with probability of the binomial distribution equal to (a) for *de novo*, number of affected individuals divided by the total number of affected plus unaffected individuals; (b) for inherited,

0.5; and (c) for case-control, the number of cases divided by the total number of cases plus controls in the case-control studies.

#### MPC evaluation against other missense deleteriousness prediction metrics

We compared MPC to the following other *in silico* missense deleteriousness predictors: AlphaMissense[13], popEVE[14], REVEL[15], CADD[16,17], PolyPhen-2[18], and SIFT[19] (**Supplementary Fig. 18**). To assess each predictor's ability to stratify case and control variation, we annotated the case and control *de novo* missense variants from each developmental cohort. Variants were filtered to only those scored by all predictors and in high-coverage MCRs. For each developmental cohort and each *in silico* predictor, we then ranked the case and control variants based on their predicted scores. We assessed the proportion of case to control variants among the variants with the top 1%, 5%, or 10% and compared to the remaining 99%, 95%, and 90% of variants, respectively, to obtain an odds ratio using Fisher's exact test.

### Supplementary Figures

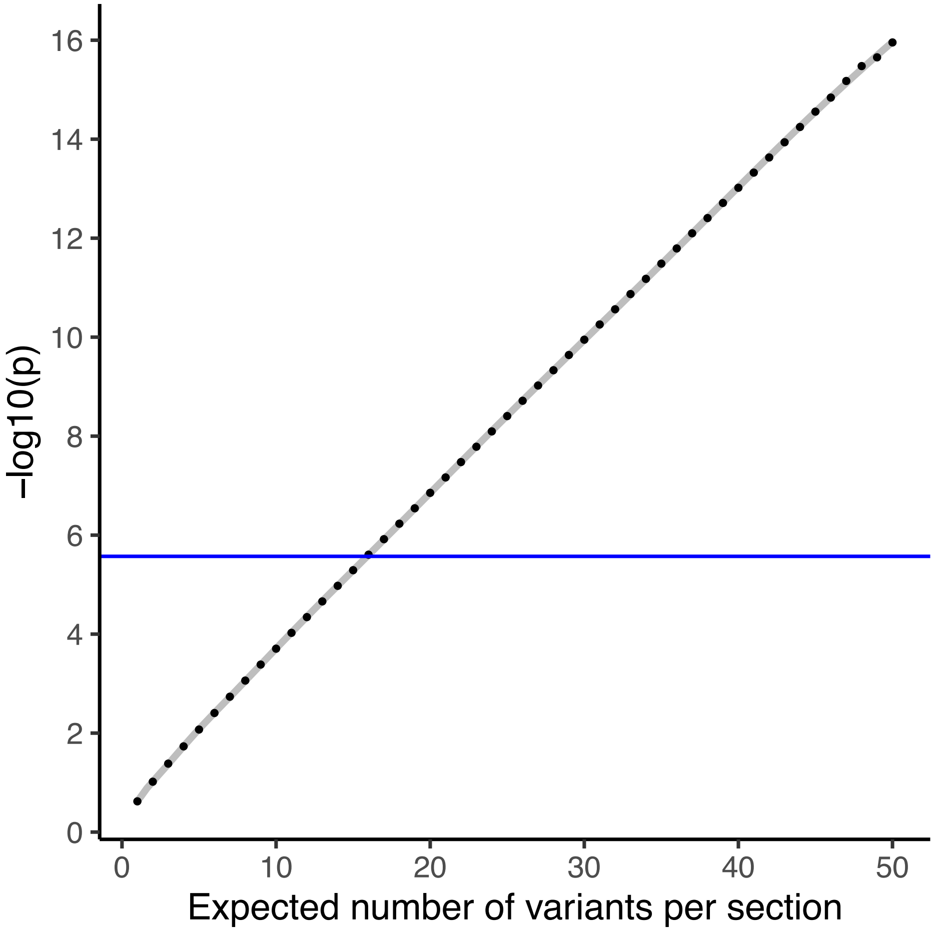

**Supplementary Fig. 1**

Statistical significance corresponding to the number of expected rare missense variants in each of two MCRs in the scenario where there are an equal number of expected variants in both MCRs, the OE of one MCR is 1.0, and the OE of the other MCR is 0.0. Blue line at $2.8 \times 10^{-6}$ corresponds to exome-wide significance (0.05 / 17841 transcripts).

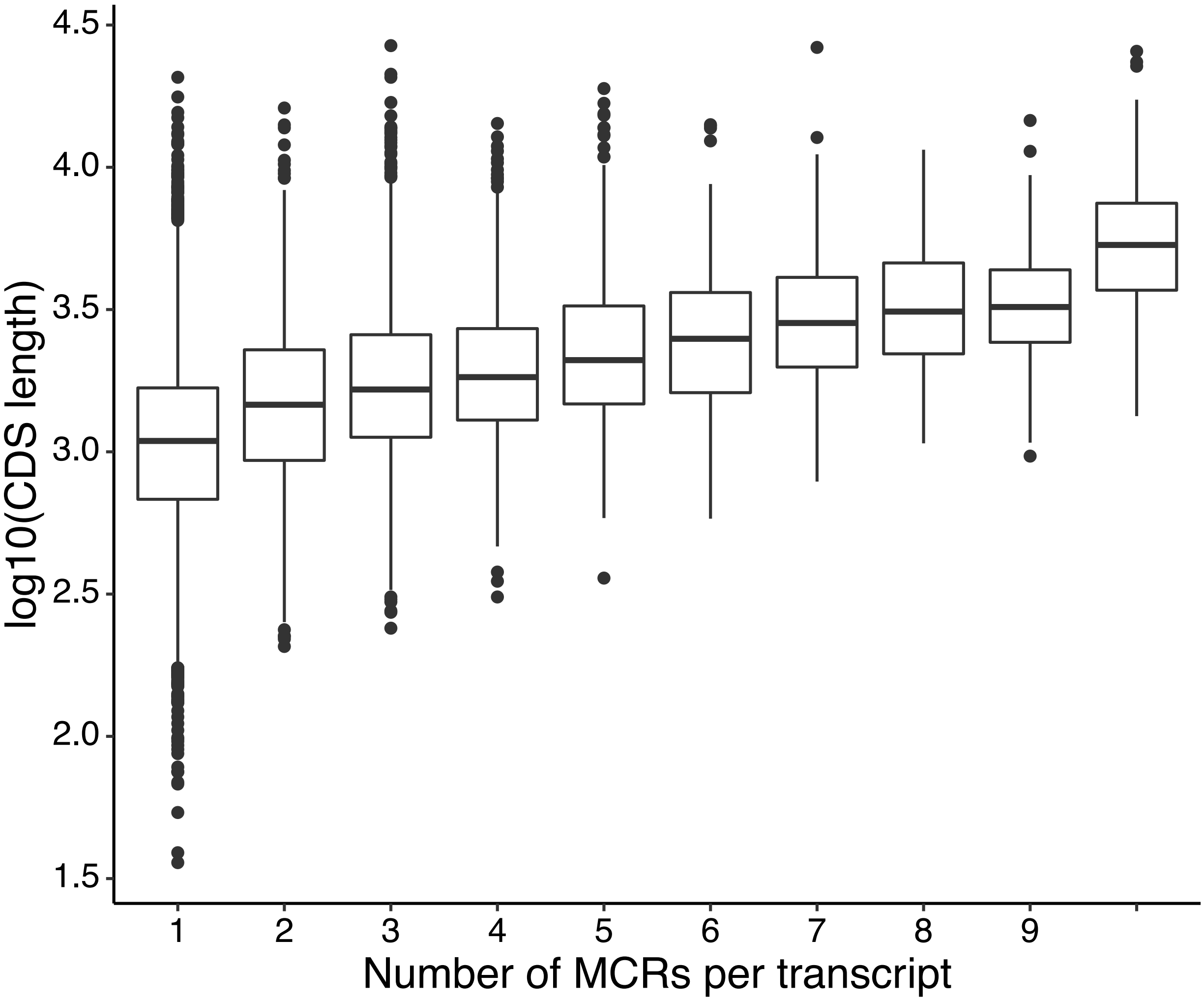

**Supplementary Fig. 2**

Transcript length is correlated with missense constraint region (MCR) count (Spearman ρ = 0.41, $p < 10^{-18}$). Coding sequence (CDS) length is measured in base pairs.

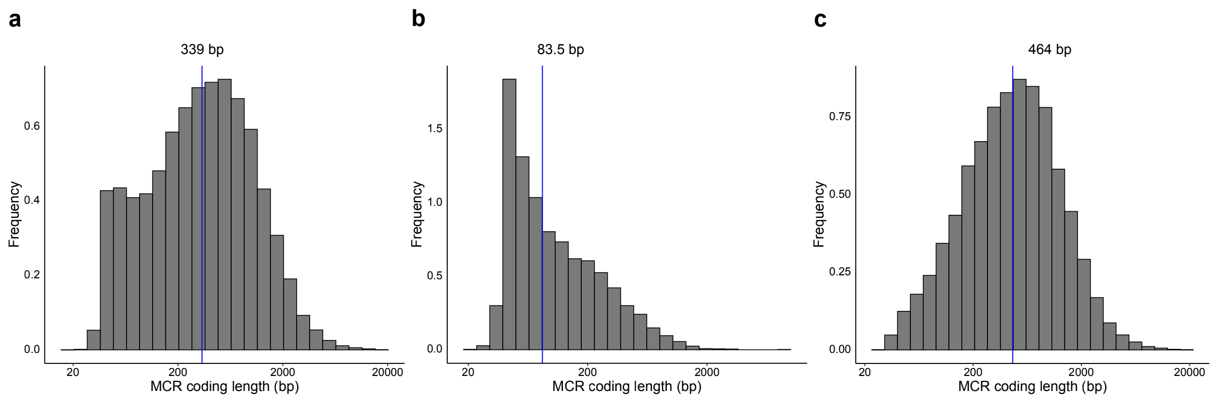

**Supplementary Fig. 3**

Missense constraint region (MCR) coding length across 6,361 transcripts harboring regional variability in missense constraint. **a**, Distribution of coding length. Median is 339 base pairs. **b**, Distribution of coding length for MCRs with strong missense constraint (missense OE < 0.4). Median is 83.5 base pairs. **c,** Distribution of coding length for remaining MCRs (missense OE ≥ 0.4). Median is 464 base pairs.

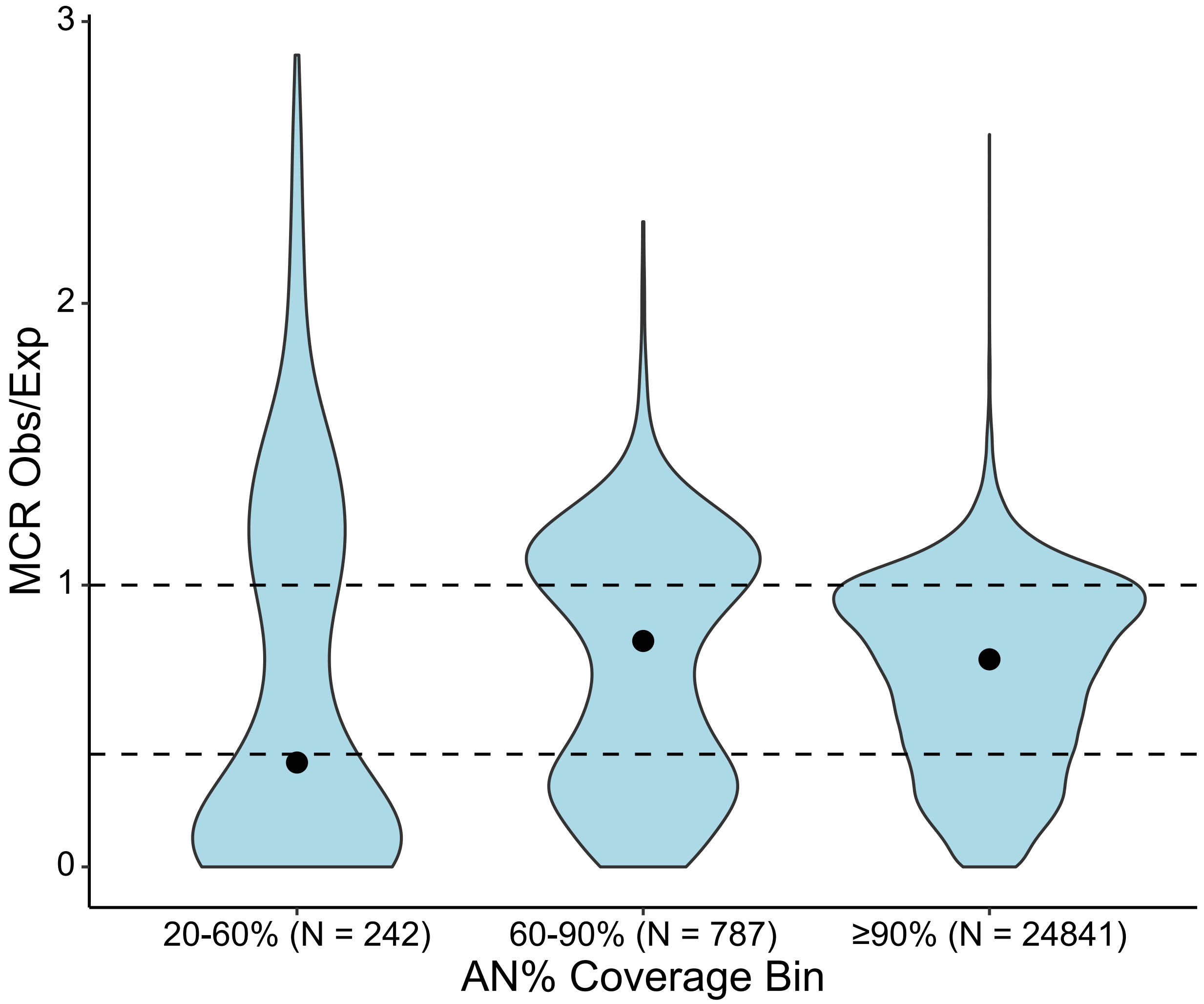

**Supplementary Fig. 4**

For the 6,361 transcripts with 2+ MCRs, the missense OE distribution over MCRs categorized by median %AN within each MCR (proxy for sequencing coverage). Dashed lines correspond to a neutral OE of 1.0 and a severely depleted OE at 0.4. Dots indicate the median value.

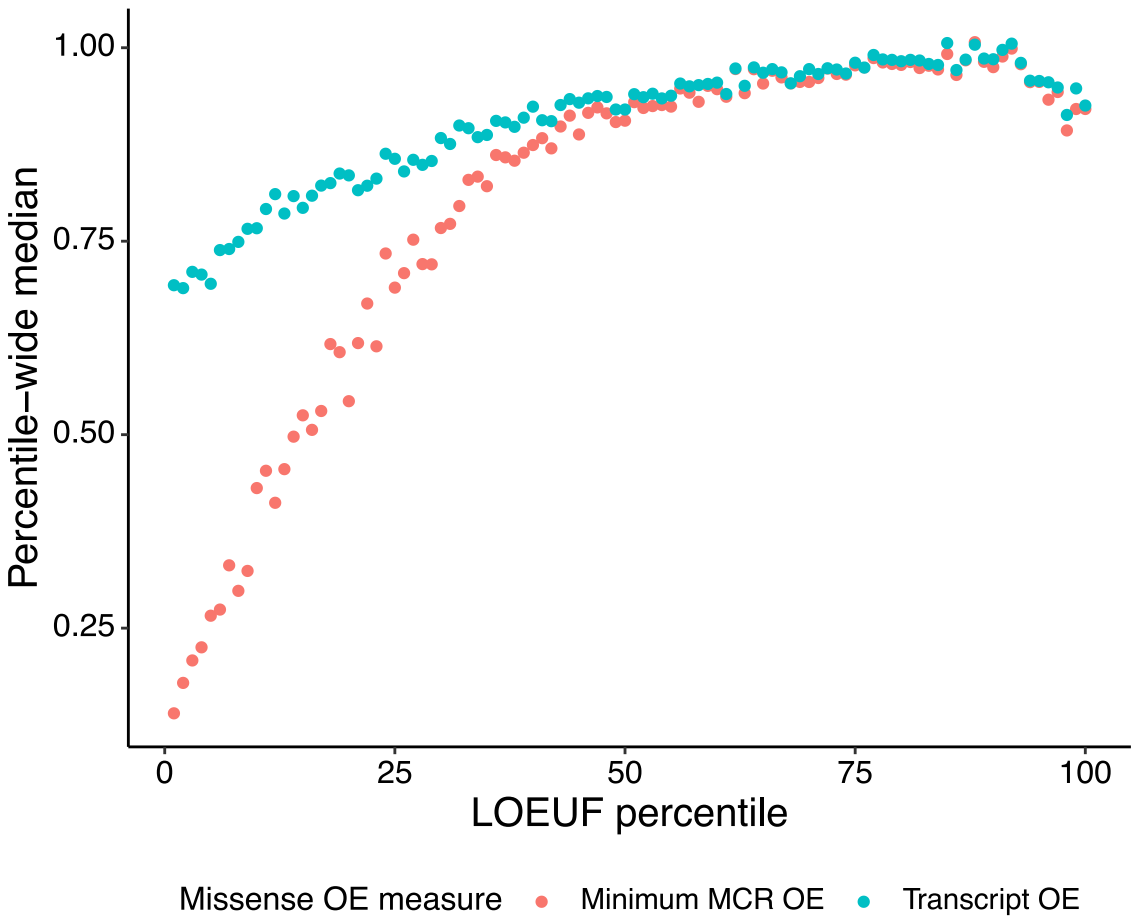

**Supplementary Fig. 5**

Loss-of-function (LoF)-intolerant transcripts tend to be intolerant to missense variation. Red: Minimum missense constraint region (MCR)-level missense OE; blue: transcript-level missense OE. The medians over transcripts within each LOEUF percentile are shown for each missense OE measure. For transcripts with one MCR, minimum MCR OE is the same as the full transcript OE.

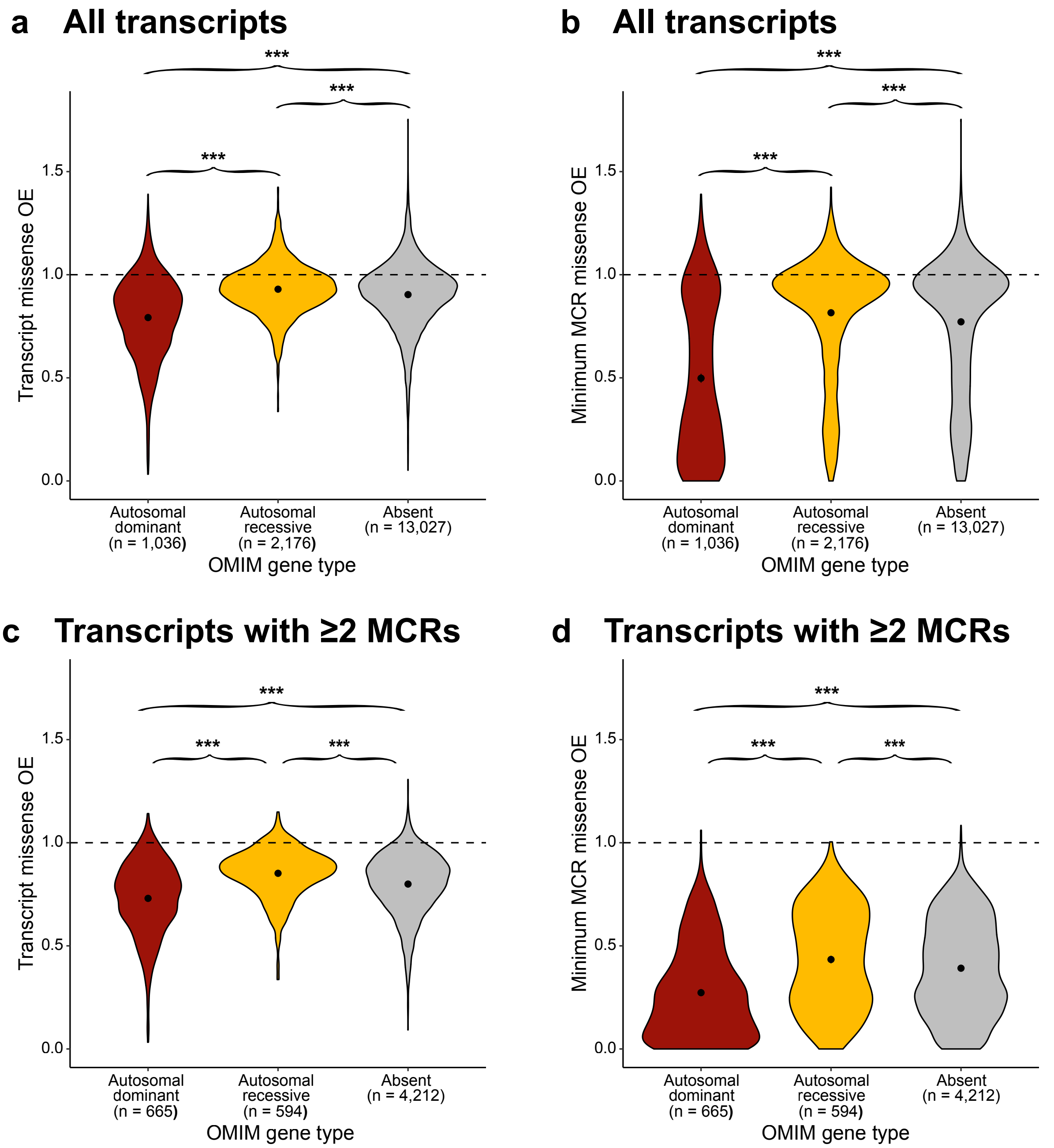

**Supplementary Fig. 6**

Genes without disease associations in OMIM are less missense-constrained than genes with autosomal dominant disease inheritance but more missense-constrained than genes with autosomal recessive disease inheritance. Missense OE per transcript measured for **a,** all transcripts and quantified over the whole transcript, **b,** all transcripts and quantified within the MCR with minimum OE, **c,** only transcripts with ≥2 MCRs and quantified over the whole transcript, and **d,** only transcripts with ≥2 MCRs and quantified within the MCR with minimum OE. One transcript with minimum MCR missense OE > 4 is excluded. Means and their 95% confidence intervals are marked. The Welch's t-test was used to compare transcript missense

OE distributions, and the Wilcoxon test was used to compare MCR missense OE distributions. One-sided tests were performed for the alternative hypotheses that genes with autosomal recessive inheritance are less constrained than non-OMIM genes, which in turn are less constrained than genes with autosomal dominant inheritance. *** = $p < 0.001$.

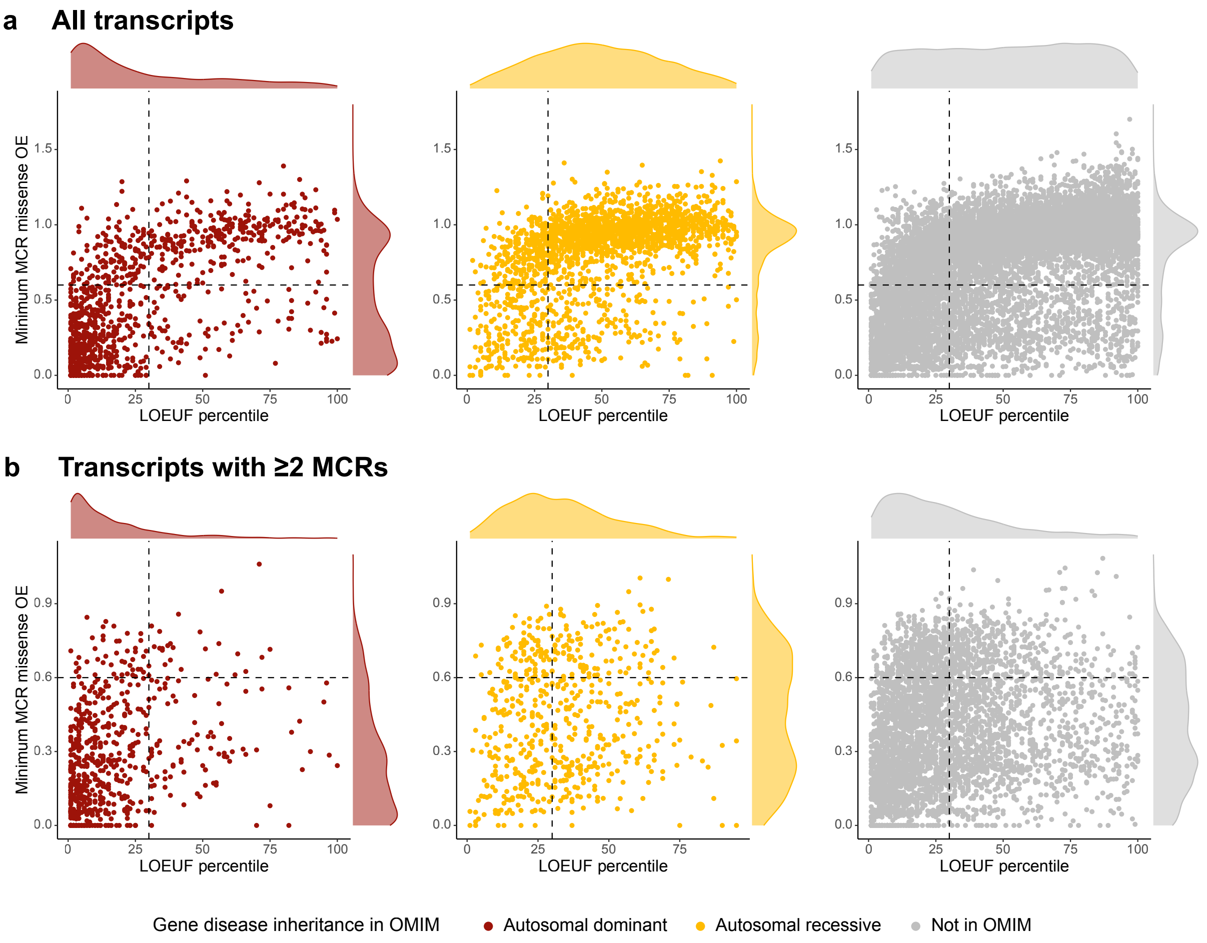

**Supplementary Fig. 7**

Loss-of-function (LoF) and missense constraint suggest additional potential for disease gene discovery. Missense OE in the MCR with minimum OE for each transcript across **a,** all transcripts (n=1,036, 2,176, and 12,623 for genes with autosomal dominant inheritance, autosomal recessive inheritance, or no disease annotation in OMIM, respectively) or **b,** only transcripts with ≥2 MCRs (n=665, 594, and 4,083 for genes with autosomal dominant inheritance, autosomal recessive inheritance, or no disease annotation in OMIM, respectively). For transcripts with only one MCR in **a**, this is equivalent to the transcript-level missense OE. Black dashed lines at 30th percentile of LOEUF and minimum MCR missense OE of 0.6 demarcate LoF- and missense-constrained transcripts. One transcript with minimum MCR missense OE > 4 is excluded.

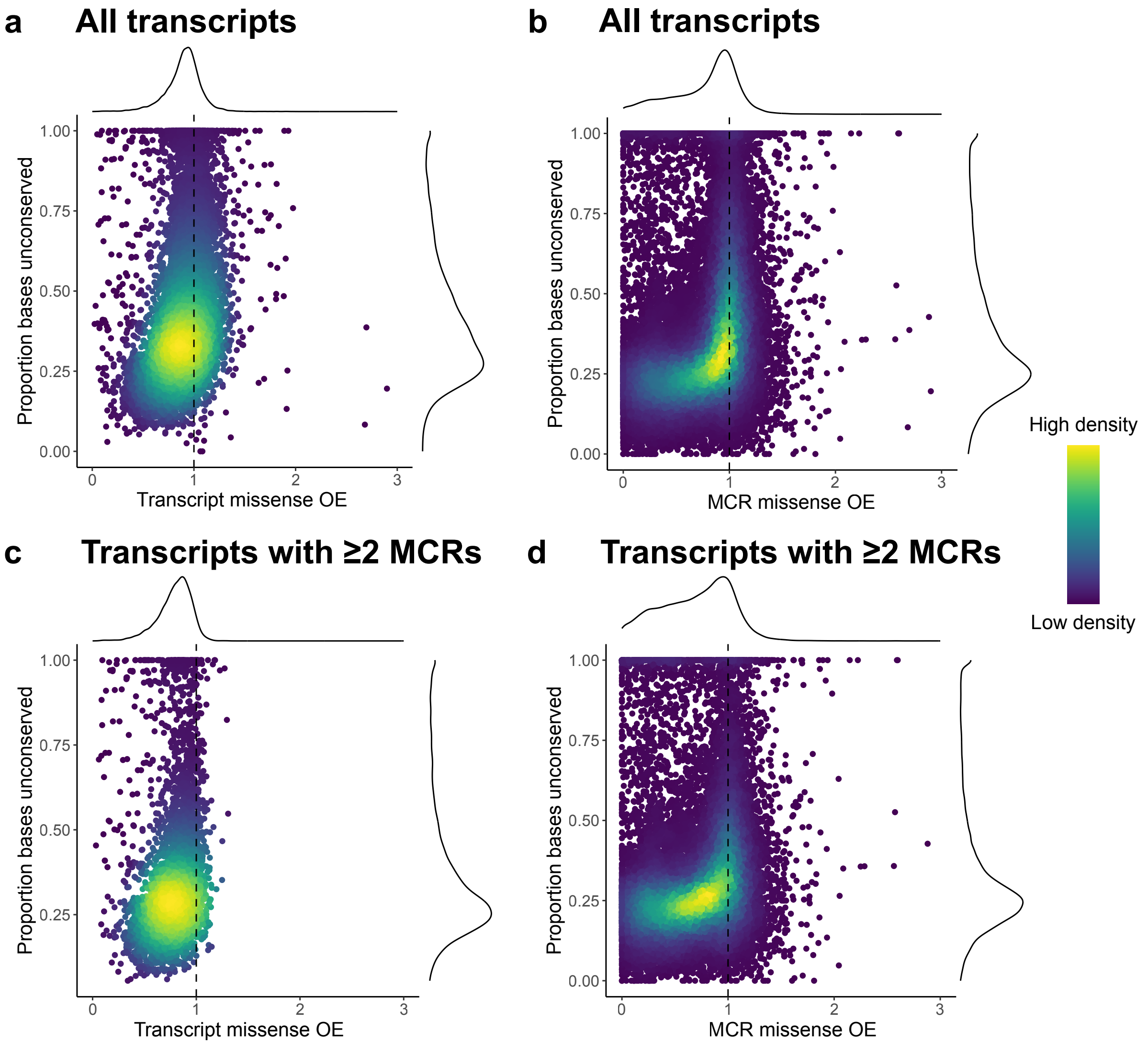

**Supplementary Fig. 8**

Divergence of missense constraint and evolutionary conservation suggests human-specific selective pressures. Missense observed/expected (OE) vs. proportion of coding bases that are unconserved (phyloP threshold < 2.7)[3] across **a**, entire transcripts (n=17,839 with phyloP scores; Spearman ρ = 0.50; $p < 10^{-50}$), **b**, missense constraint regions (MCRs) in these 17,839 transcripts (Spearman ρ = 0.449; $p < 10^{-50}$), **c**, entire transcripts (transcripts with ≥2 MCRs only; n=6,361; Spearman ρ = 0.45; $p < 10^{-50}$), and **d**, MCRs in these 6,361 transcripts (Spearman ρ = 0.32; $p < 10^{-50}$). Lighter colors indicate greater density of points.

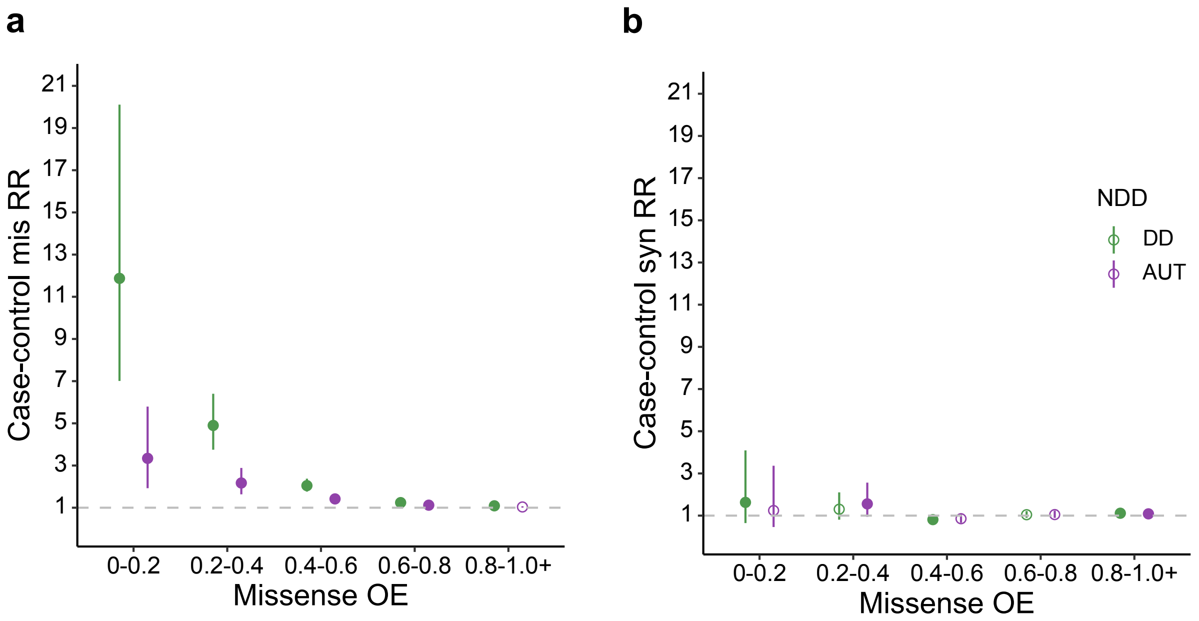

**Supplementary Fig. 9**

Rate ratios of *de novo* **a,** missense or **b,** synonymous variants over missense constraint region (MCR) observed/expected (OE) bins in individuals with developmental disorders (DD; green) or autism (AUT; purple) relative to unaffected siblings. Points are solid colored if the difference from 1 is statistically significant under Bonferroni correction (Poisson two-sided $p < 0.0025$).

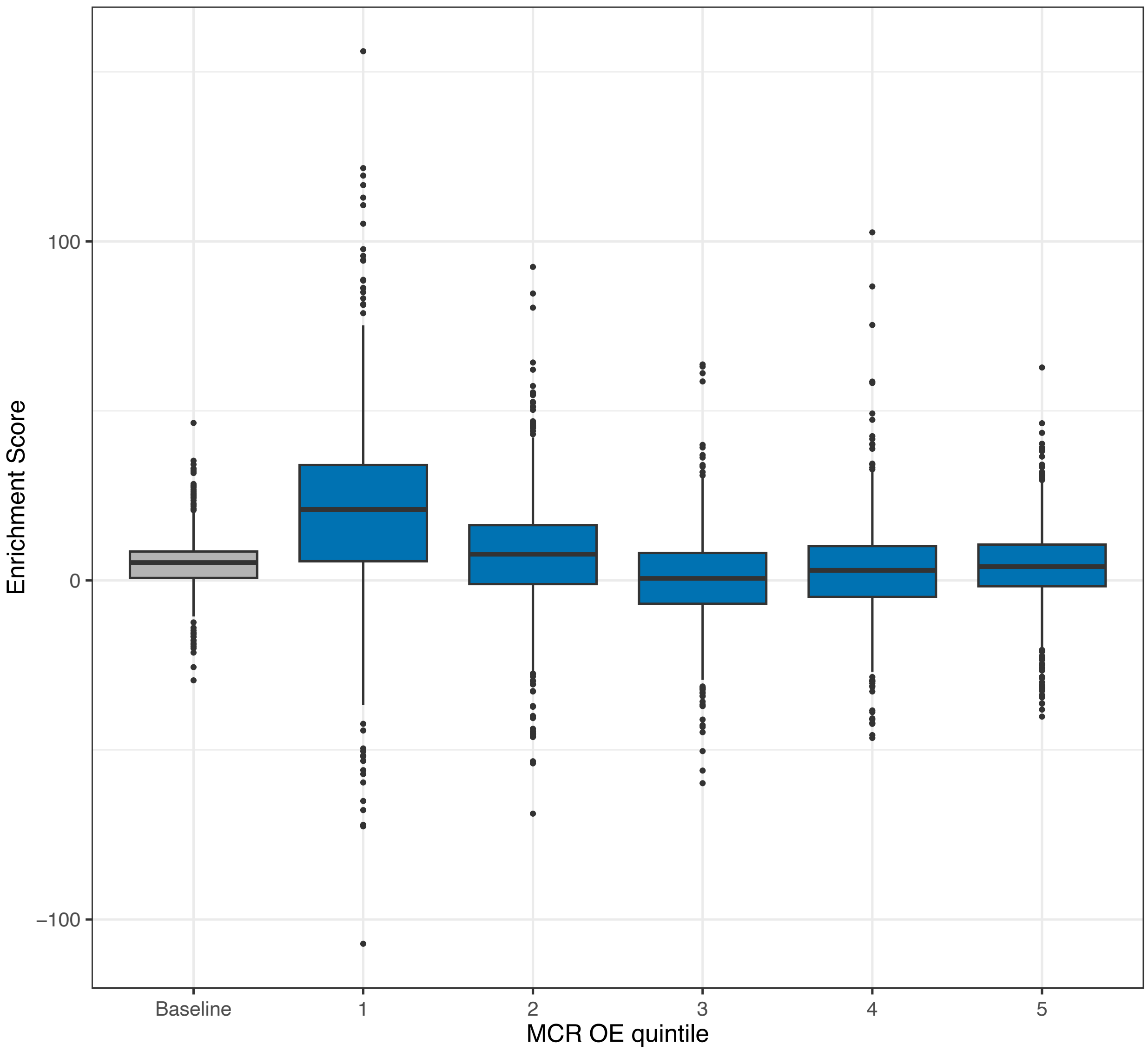

**Supplementary Fig. 10**

SNP heritability enrichments relative to the average genome-wide SNP across 268 independent traits in each MCR quintile by OE compared to the baseline enrichment across all MCRs.

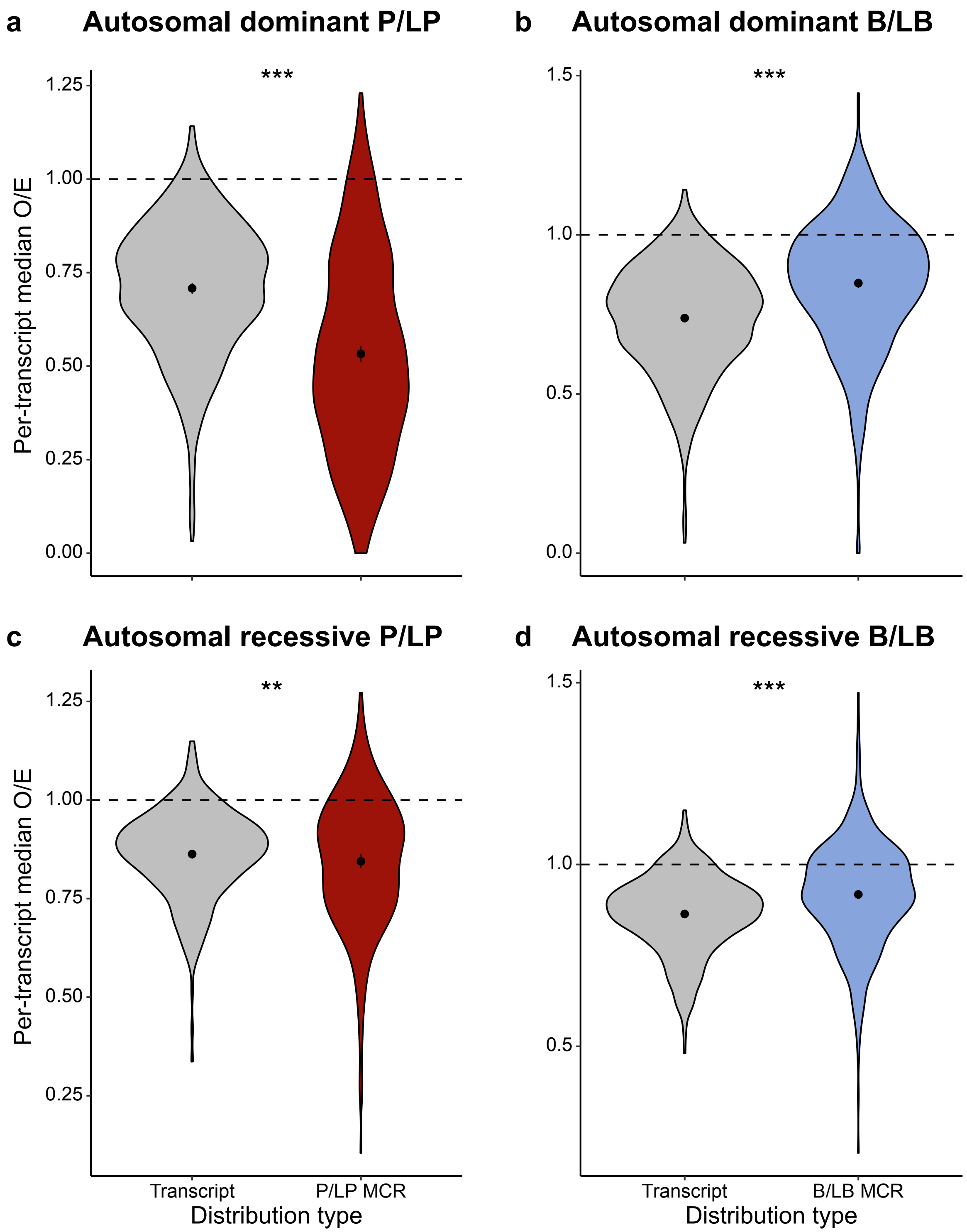

**Supplementary Fig. 11**

ClinVar pathogenic/likely pathogenic (P/LP) vs. benign/likely benign (B/LB) localization across missense observed/expected (OE) in disease-associated genes. **a,** P/LP variants in autosomal dominant genes (529 genes); **b,** B/LB variants in autosomal dominant genes (n=615 genes); **c and d,** P/LP variants in autosomal recessive genes (n=333 genes); **d,** B/LB variants in autosomal recessive genes (n=532 genes). For the P/LP and B/LB distributions, we annotated each variant with the missense OE across the MCR they fell in and collapsed these values within each transcript by taking the respective medians. Means of these distributions and their 95% confidence intervals are marked. ** = Wilcoxon $p < 0.01$; *** = Wilcoxon $p < 0.001$.

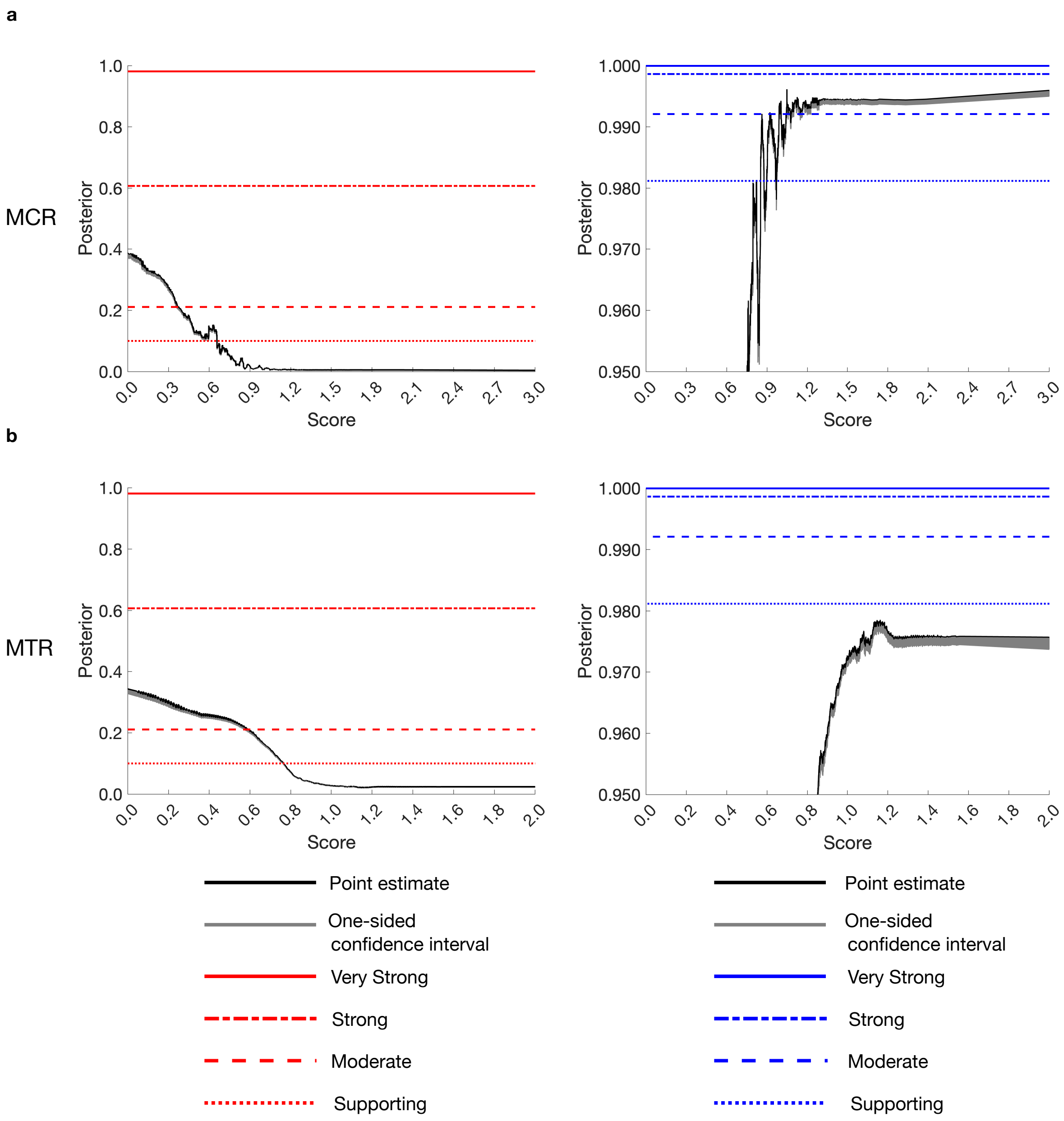

**Supplementary Fig. 12**

Clinical calibration of regional constraint metrics. Clinical calibration for pathogenic (red, left) and benign (blue, right) variation of **a**, Missense constraint region (MCR) missense observed/expected (OE); **b**, Missense tolerance ratio (MTR)[10]. Horizontal lines indicate thresholds required to meet ACMG/AMP evidence levels.

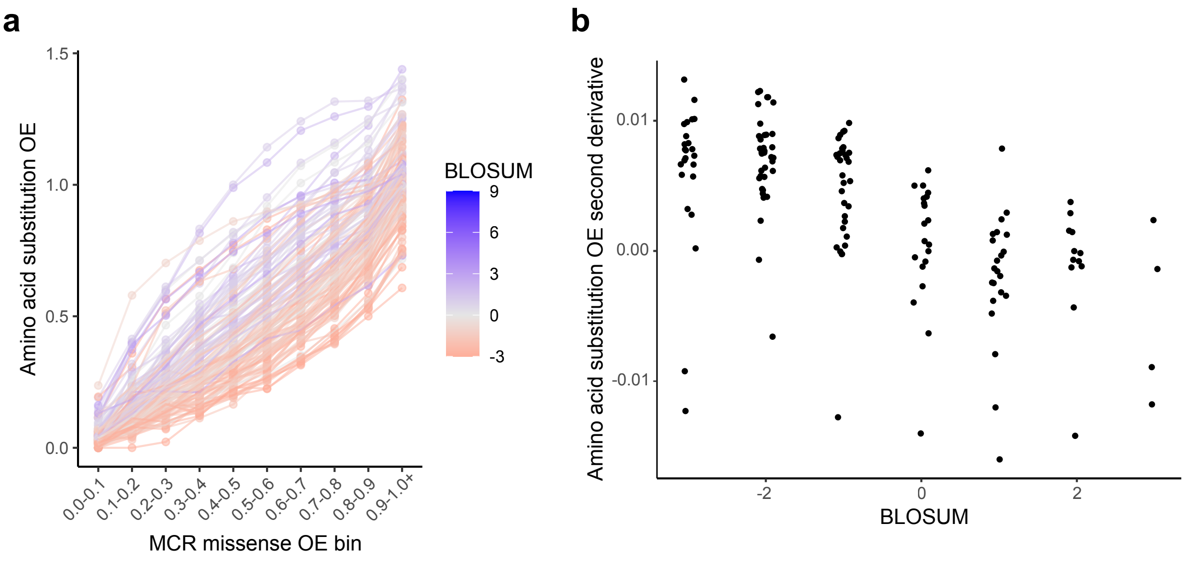

**Supplementary Fig. 13**

MCR missense OE-derived metrics quantifying amino acid substitution class severity vs. BLOSUM[20]. **A,** Missense OE per amino acid substitution class calculated within each MCR missense OE bin, colored by BLOSUM score for each amino acid substitution class. Blue indicates lower severity and red higher severity. **B,** Second derivative of linear regressions applied to **a** (termed “amino acid substitution OE second derivative”) vs. BLOSUM scores (Pearson r = -0.575, $p = 7.1 \times 10^{-15}$).

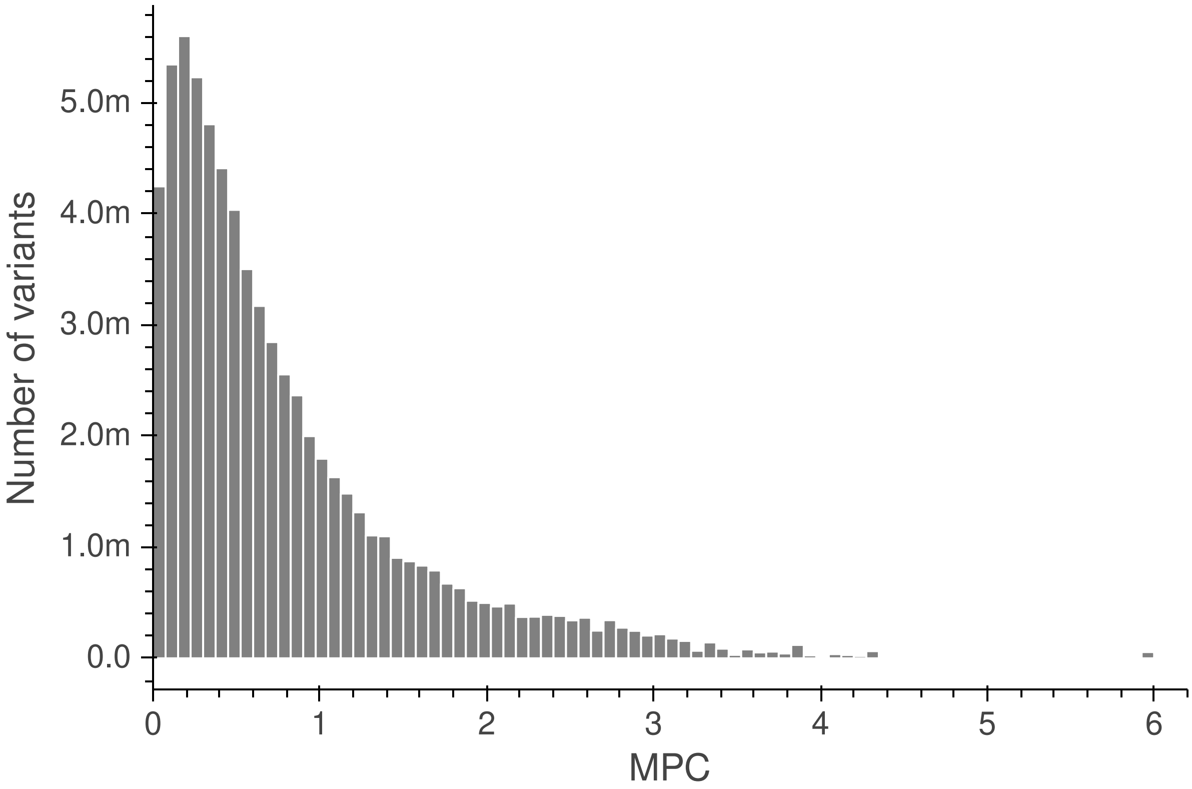

**Supplementary Fig. 14**

MPC distribution over missense variants in 17,841 transcripts. The “m” indicates million.

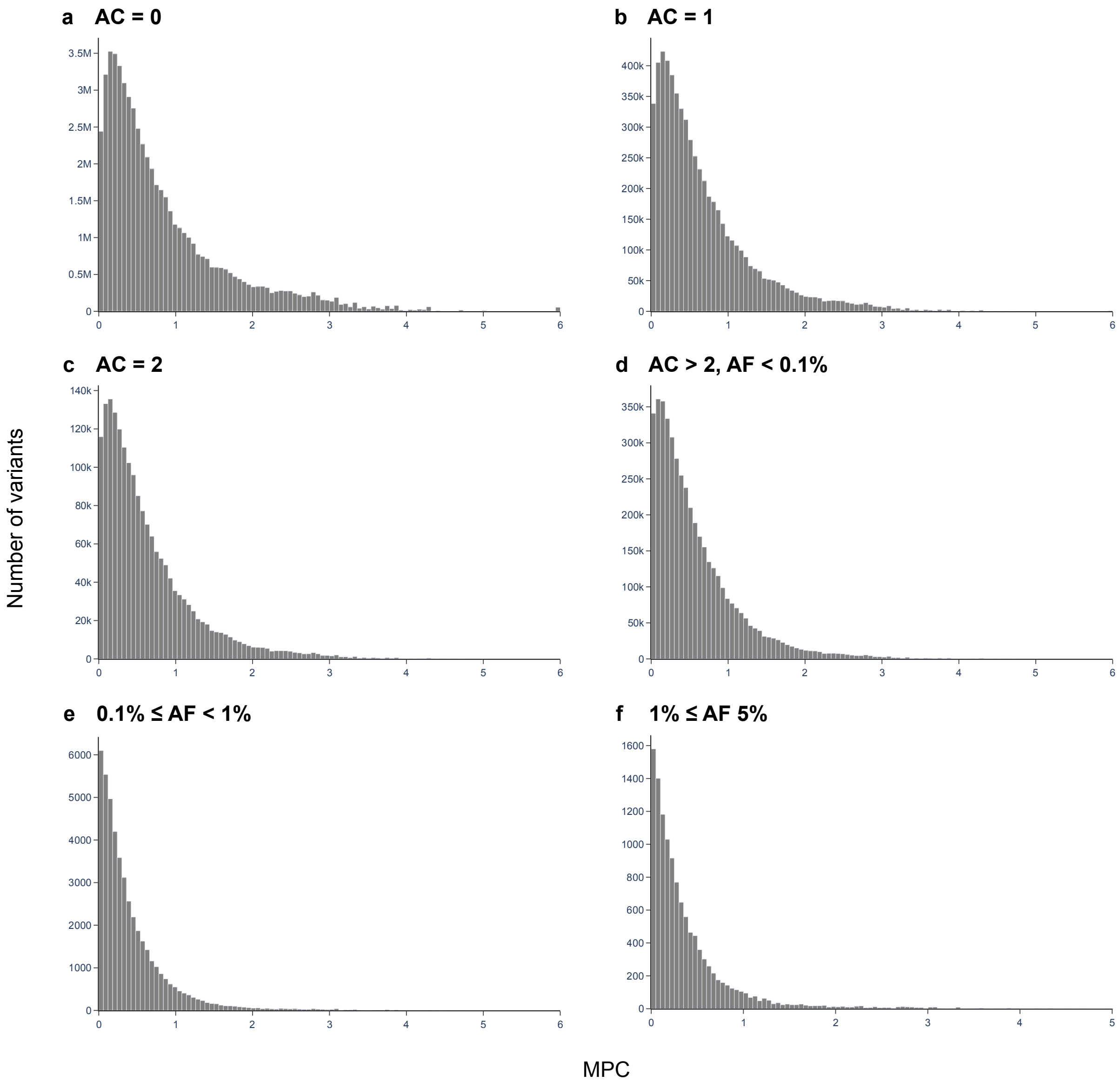

**Supplementary Fig. 15**

MPC distribution over missense variants by gnomAD allele frequency. Here, “M” indicates million and “K” indicates thousand.

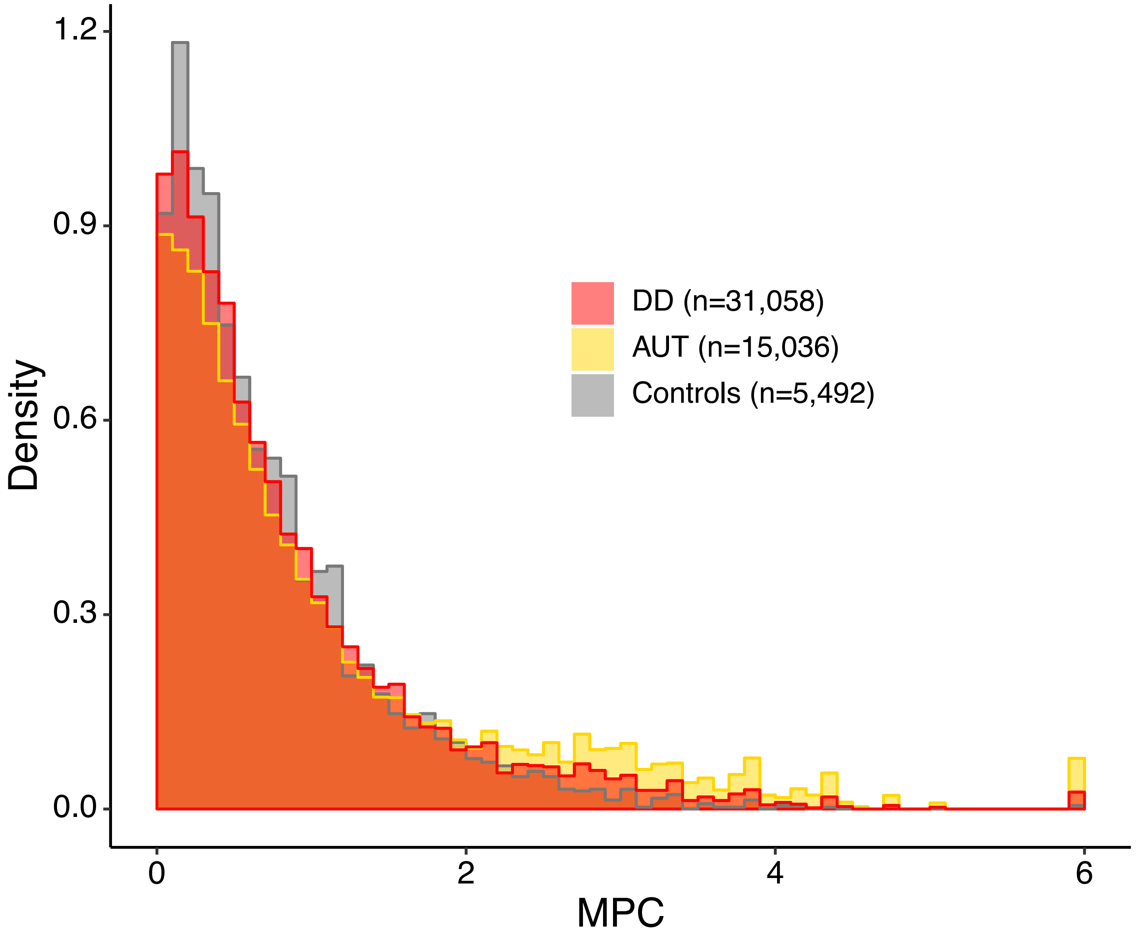

**Supplementary Fig. 16**

MPC distribution for *de novo* missense variants in developmental disorder (DD), autistic (AUT), and unaffected sibling cohorts. MPC scores for DD and AUT *de novo* missense variants are significantly higher than in unaffected siblings (Wilcoxon $p < 10^{-43}$ and $= 5.2 \times 10^{-6}$, respectively). MPC distribution medians are 0.68, 0.57, and 0.53 for DD, AUT, and unaffected siblings, respectively. The area under each curve sums to 1.

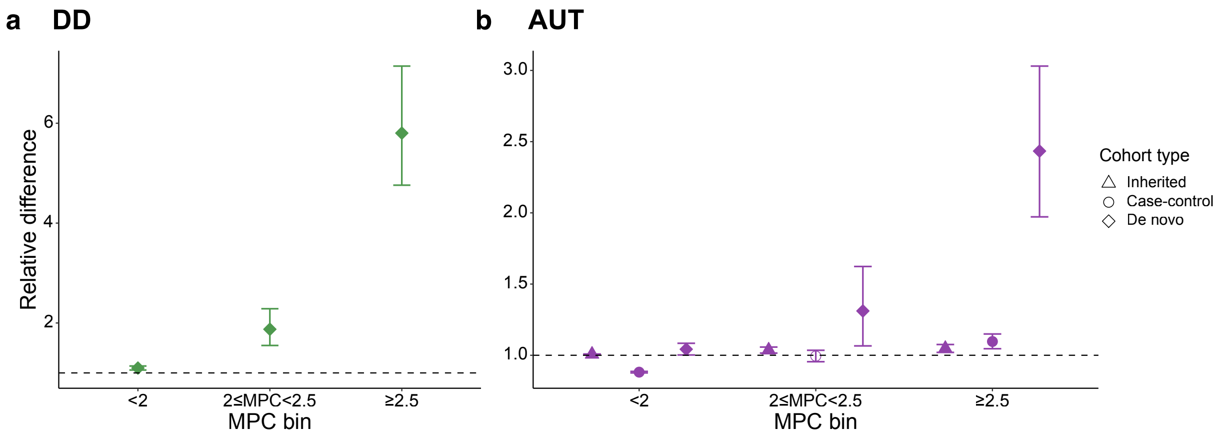

**Supplementary Fig. 17**

MPC effectively stratifies case and control variation. The difference relative to controls of missense variants stratified by MPC score without partitioning by gene for **a,** individuals with DD and **b,** autistic individuals (AUT). Relative difference is calculated as: for *de novo* variants, the average rate of variants in probands divided by that in sibling controls; for case-control, the average rate of variants in cases divided by that in controls from case-control data; for inherited, the average rate in probands of transmitted variants divided by that of untransmitted variants. Error bars represent 95% confidence intervals calculated from a binomial test. Points are solid colored if the difference from 1 is statistically significant (binomial or Fisher's exact $p < 0.05$).

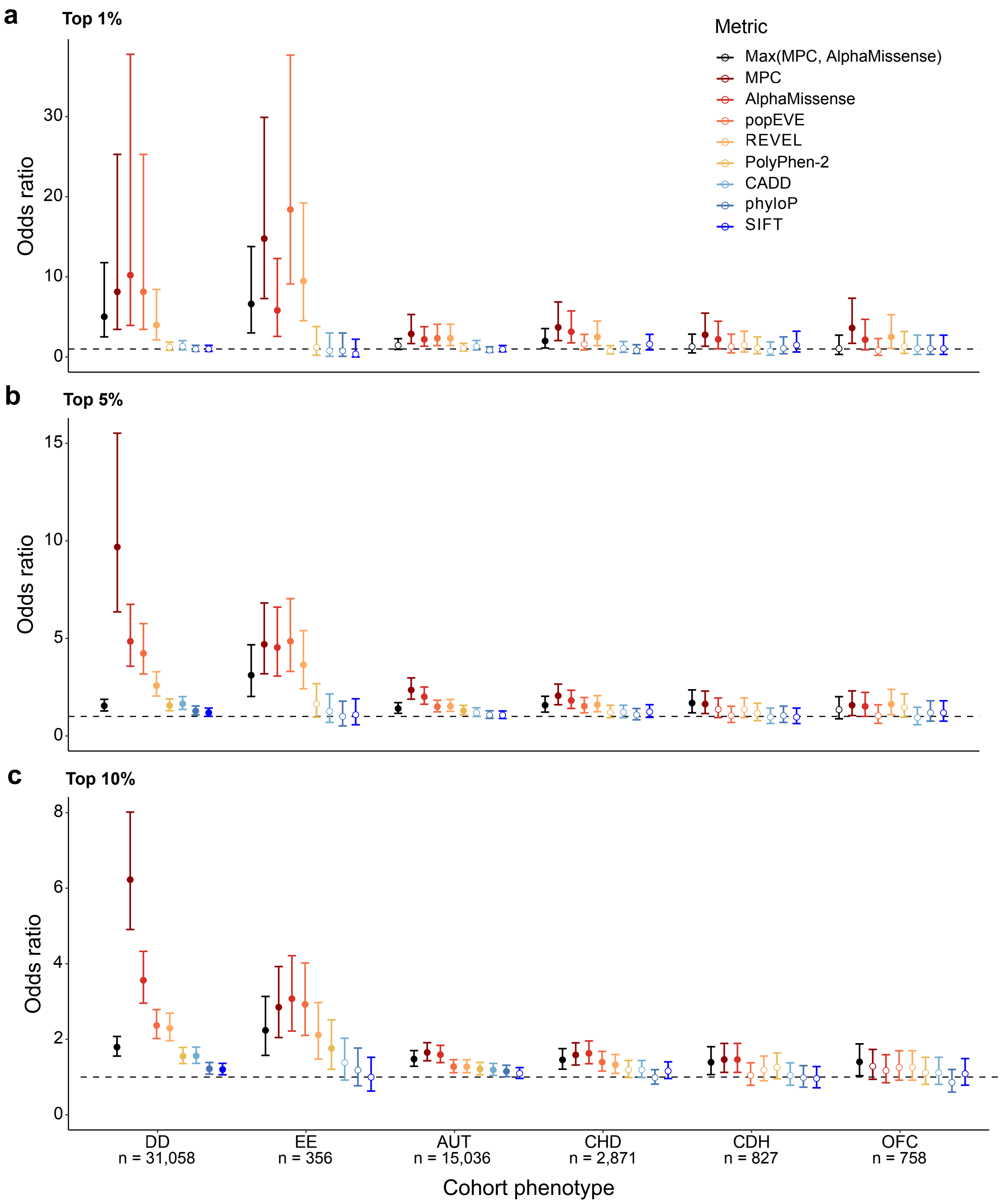

**Supplementary Fig. 18**

MPC stratifies case and control variation at different thresholds. The odds ratio of case to control *de novo* missense variants in the top **a,** 1%, **b,** 5%, and **c,** 10% of respective rankings. *De novo* missense variants from each case cohort are ranked against those in the 5,492 controls for each predictor. DD: developmental disorders, EE: epileptic encephalopathy, AUT: autism, OFC: orofacial cleft, CHD: congenital heart disease, CDH: congenital diaphragmatic hernia. Error bars represent 95% confidence intervals. Only variants scored by all predictors are included. Points are solid colored if the difference from 1 is statistically significant (Fisher's exact $p < 0.05$). "Max(MPC, AlphaMissense)" takes the maximum ranking of each variant for MPC or AlphaMissense before calculating enrichments.

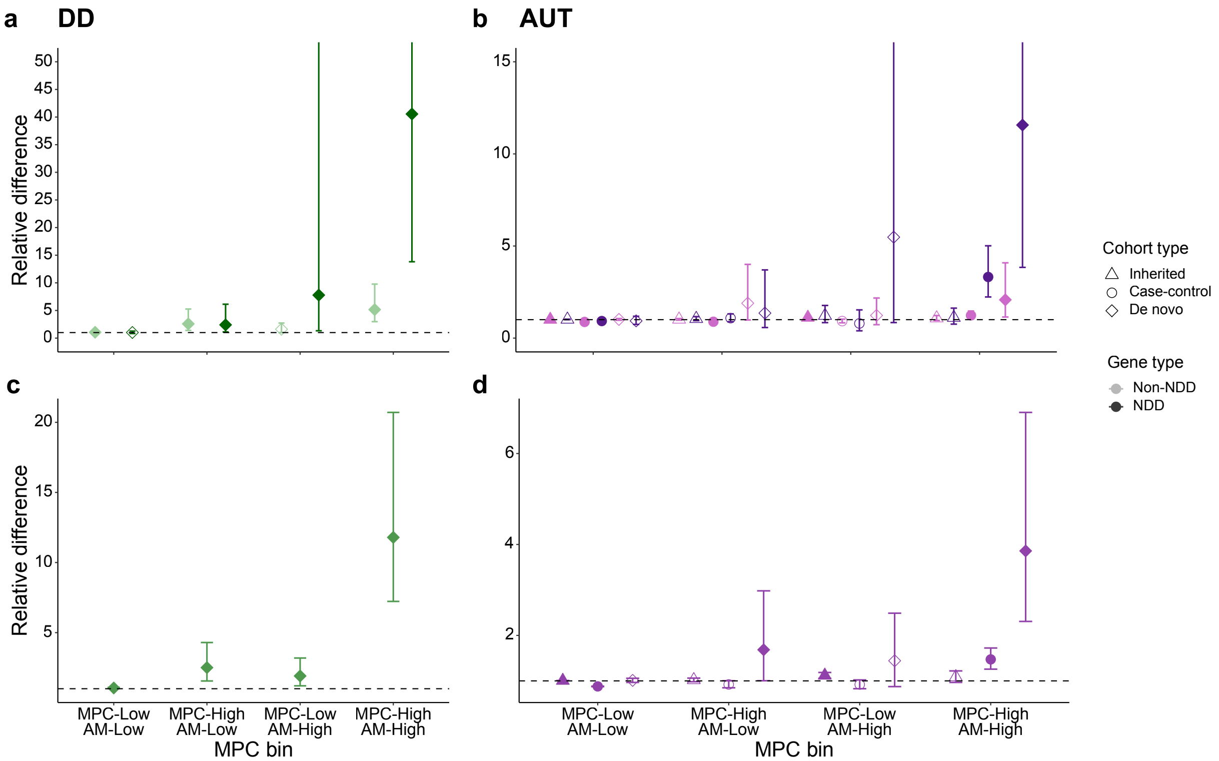

**Supplementary Fig. 19**

AlphaMissense and MPC contain complementary information for missense variant prioritization. The difference relative to controls of missense variants stratified by MPC and AlphaMissense score for DD **a,** with and **c,** without partitioning by gene and for AUT **b,** with and **c,** without partitioning by gene. “MPC-High” is defined as MPC ≥ 2.5 and “MPC-Low” is MPC < 2; “AM-High” is defined as AlphaMissense score ≥ 0.9985 and “AM-Low” is AlphaMissense score < 0.761. Only variants scored by both predictors are included. Y-axis is truncated in **a,** at 50 and in **b,** at 15 for readability. Points are solid colored if the difference from 1 is statistically significant (binomial or Fisher’s exact $p < 0.05$).

### Legends for Supplementary Tables in separate file

**Supplementary Table 1**
Annotations for transcripts used in analysis. Filtered to high-coverage MCRs.

**Supplementary Table 2**
Percentiles of missense OE across possible sites of missense variants in 17,841 transcripts.

**Supplementary Table 5**
*De novo* missense variants in the individuals with DDs, autistic individuals, and control siblings annotated with MCR missense OE and MPC. Filtered to those in high-coverage MCRs. DD = developmental disorder.

**Supplementary Table 6**
*De novo* synonymous variants in the individuals with DDs, autistic individuals, and control siblings annotated with MCR missense OE and MPC. Filtered to those in high-coverage MCRs. DD = developmental disorder.

**Supplementary Table 7**
Enrichments of DD and AUT *de novo* missense variants in high-coverage MCRs compared to controls, by MCR missense OE bin. DD = developmental disorder. AUT = autism.

**Supplementary Table 8**
Enrichments of DD and AUT *de novo* synonymous variants in high-coverage MCRs compared to controls, by MCR missense OE bin. DD = developmental disorder. AUT = autism.

**Supplementary Table 9**
SNP heritability enrichments across 268 independent traits partitioned by MCR missense OE. Filtered to high-coverage MCRs.

**Supplementary Table 10**
Summary of SNP heritability enrichments across 268 independent traits partitioned by MCR missense OE. Filtered to high-coverage MCRs.

**Supplementary Table 11**
Per-transcript median MCR missense OE for ClinVar P/LP or B/LB variants in genes with autosomal dominant inheritance. Filtered to high-coverage MCRs.

**Supplementary Table 12**
Per-transcript median MCR missense OE for ClinVar P/LP or B/LB variants in genes with autosomal recessive inheritance. Filtered to high-coverage MCRs.

**Supplementary Table 14**

Amino acid substitution-level severity metrics for missense variant classes. OE-based metrics are calculated using high-coverage MCRs.

**Supplementary Table 16**

Enrichments of rare and *de novo* variants in DD and AUT by MPC bin and NDD gene association. Filtered to high-coverage MCRs. DD = developmental disorder. AUT = autism.

**Supplementary Table 17**

Odds ratio enrichments of *de novo* variants in developmental disorder (DD) vs. control cohorts across *in silico* missense deleteriousness predictors. Filtered to variants in high-coverage MCRs scored by all predictors.

**Supplementary Table 18**

Enrichments of rare and *de novo* variants in DD and AUT by MPC and AlphaMissense bin and NDD gene association. Filtered to high-coverage MCRs. DD = developmental disorder. AUT = autism.

16. Kircher M, Witten DM, Jain P, O'Roak BJ, Cooper GM, Shendure J. A general framework

for estimating the relative pathogenicity of human genetic variants. *Nat Genet*. 2014;46(3):310-315.

17. Rentzsch P, Schubach M, Shendure J, Kircher M. CADD-Splice-improving genome-wide variant effect prediction using deep learning-derived splice scores. *Genome Med*. 2021;13(1):31.

18. Adzhubei IA, Schmidt S, Peshkin L, et al. A method and server for predicting damaging missense mutations. *Nat Methods*. 2010;7(4):248-249.

19. Ng PC, Henikoff S. SIFT: Predicting amino acid changes that affect protein function. *Nucleic Acids Res*. 2003;31(13):3812-3814.

20. Henikoff S, Henikoff JG. Amino acid substitution matrices from protein blocks. *Proc Natl Acad Sci U S A*. 1992;89(22):10915-10919.
